# Genomic repeats for single-cell molecular recording

**DOI:** 10.64898/2026.08.06.743335

**Authors:** Rachel K. Dveirin, Jessica D. Lin, Preeti Vyas, Jiaxi Lu, Yuqing Yan, Jennifer J. Lee, Xinzhong Dong, Sujatha Kannan, Ben Langmead, Sashank K. Reddy, Dingchang Lin, Reza Kalhor

## Abstract

Genomic recording enables transient biological signals to be indelibly captured through DNA alterations, creating a permanent record of cellular history retrievable by sequencing. However, current methods are limited by scarce writing space, typically targeting only one or a few amenable genomic sites and requiring large cell populations for signal reconstruction. Here, we establish Repeats for Genomic Recording (RGRs): sequences with up to 400 copies targetable by a single CRISPR guide RNA, readable with a common primer pair, and predicted to have minimal functional impact. We demonstrate that RGRs enable both signal deconvolution in single cells and high-resolution recording in cell populations. Individual RGR sites exhibit distinct response kinetics; thus, combining them improves recording resolution beyond what redundancy alone provides, analogous to diversity reception in wireless communication. We develop a computational pipeline for systematic RGR identification, revealing 15,000 to 25,000 candidates per species across human, mouse, and zebrafish, thereby markedly expanding recording capacity and enabling cell-type-specific applications. Finally, we validate RGRs in live mice by recording long-term immediate early gene activity across the brain following epilepsy induction. This work establishes genomic repeats as a high-capacity platform for single-cell molecular recording in vivo.

## Introduction

Capturing the temporal dynamics of biological processes typically requires destructive serial sampling: sacrificing cells or organisms at successive timepoints to reconstruct a timeline. This approach demands large sample numbers, misses transient events between timepoints, and cannot follow the same cells longitudinally^1,2^. Genomic recording offers a promising solution by leveraging DNA as a continuously present medium to document biological events over time: biological signals are converted into permanent, heritable changes in genomic DNA, enabling reconstruction of temporal dynamics retrospectively through sequencing^3–14^.

Genomic recording strategies have shown remarkable versatility, with diverse biological processes (gene expression^12,15^, cis-regulatory element activity^14^, signaling pathway activity^3^, and cell lineage^4,7,8,10^) successfully linked to DNA writing systems including nucleases, base editors, prime editors, and compound architectures such as DNA Typewriter, peCHYRON, TRACE, and CAMERA^13,14,16^. These approaches promise to transform our understanding of embryonic development^4,17^, cancer progression^18–21^, and immune response^3^. Yet, despite this progress, a fundamental bottleneck has limited the resolution and scale of genomic recording: the scarcity of efficiently accessible and retrievable writing space in the genome.

The core challenge is information capacity. Most recording approaches write to only one or a handful of genomic sites, and per-site capacity is inherently modest. Standard CRISPR base-editing and prime-editing systems can record approximately one bit per target site per cell (edited vs unedited). CRISPR nuclease systems can generate 3–5 bits^22^ due to the semi-random nature of mutational outcomes^23–25^. Higher-bandwidth systems such as homing guide RNAs (hgRNAs) and DNA Typewriter expand storage to 5–10 bits per site per cell^8,13,16^. These constraints in information capture per cell necessitate populations of tens to thousands of cells for meaningful signal reconstruction, precluding single-cell recording and missing cellular heterogeneity.

Multiplexing across additional sites might seem an obvious solution, but practical constraints emerge: few validated genomic targets exist (e.g., safe harbors), and expanding target numbers increases the complexity of the recording system while complicating coordinated writing and readout. A recent approach leveraging mitochondrial genomes, which exist in hundreds of copies per cell, represents a promising step^26^. However, mitochondria remain inaccessible to CRISPR-based strategies due to the unsolved challenge of guide RNA delivery, and mitochondrial genomes lack the permanence of nuclear DNA: copies may be lost during organelle turnover and do not segregate reliably to daughter during cell division. Repetitive nuclear elements offer an alternative for expanding recording capacity. They have hundreds of copies per cell, all of which segregate stably to daughter cells. They are accessible to CRISPR as they have been leveraged for multiplexed genome engineering^27–29^ and lineage tracing^30^ in cell lines. However, their potential for high-resolution molecular recording remains unrealized, since repetitive elements constitute nearly half of vertebrate genomes^31,32^, creating a complex landscape of variable sequences that elude easy access.

To address these constraints, here we identify Repeats for Genomic Recording (RGRs) as targetable repetitive loci with three key features (**Figure 1a**). First, they are editable by a single CRISPR guide RNA, enabling multichannel recording within a single cell. Second, they are amplifiable by common primer pairs, facilitating parallel readout via sequencing. Third, they are depleted in functional DNA, minimizing the consequences of recording-induced alterations. We use a prototype with over 300 copies in the haploid human genome to achieve recording that accurately resolves four distinct signal exposure levels in single cells. Importantly, we find that this performance derives not from redundancy alone but from the diverse response kinetics of individual sites, analogous to diversity reception in signal processing, where channels with distinct characteristics outperform identical ones. We then develop a computational pipeline to identify and characterize candidate RGRs across human, mouse, and zebrafish genomes, revealing 15,000–25,000 candidates per species—a surprisingly large repertoire with suitable properties, which we validate experimentally. Finally, we leverage RGRs to record immediate early gene activity across tissues in mice by delivering an RGR-targeting sensor. We record long-term immediate early activity changes in an epilepsy model in multiple brain regions, thereby demonstrating that RGRs can capture transient signaling events over time in native cells of live animals. This work establishes both a conceptual framework and a practical resource for robust and high-capacity single-cell recording across species.

**Figure 1:**
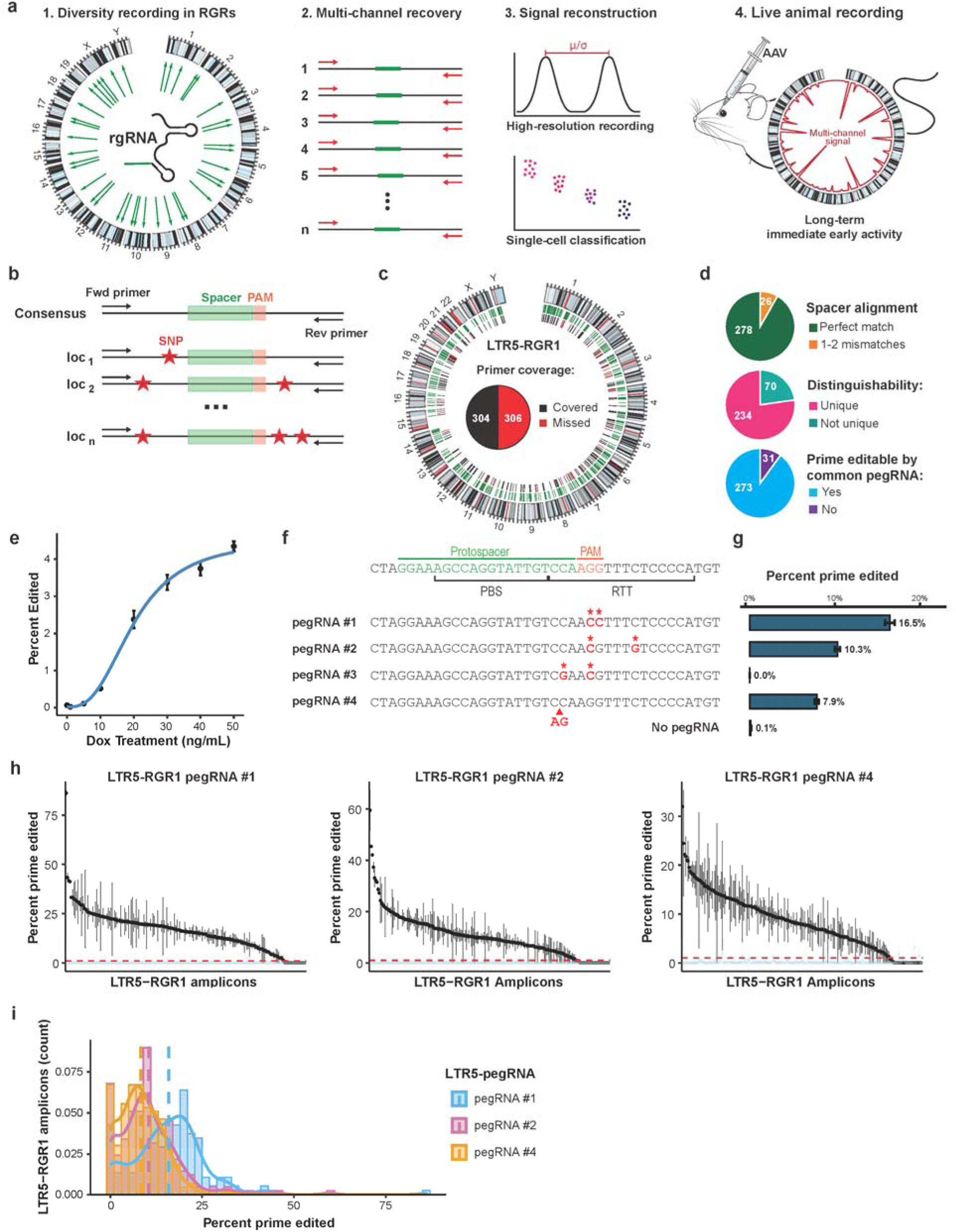
LTR5-RGR1 as a prototype for recording in genomic repeats. **a**, Graphical abstract. **b**, Schematic of ideal RGR structure. Black lines represent DNA sequences; the top line shows the consensus sequence, and subsequent lines represent individual loci. Elements labeled on the consensus sequence are conserved across copies. Spacer (green) and PAM (orange) are indicated. Forward (Fwd) and Reverse (Rev) primer binding sites are shown as arrows. Red asterisks denote single-nucleotide polymorphisms (SNPs) that distinguish individual copies from the consensus and each other. **c**, Genomic distribution and recovery of LTR5-RGR1 rgRNA target sites. First inner ring: sites targeted by LTR5 rgRNA (green). Second inner ring: amplicon coverage, with amplicons covering a target site (black) or not covering a target site (red). Pie chart (center) indicates the proportion of target sites expected to be covered by sequencing. **d**, Pie charts showing the proportion of covered target sites that perfectly match the rgRNA sequence (top), that contain at least one SNP distinguishing them from all other target-containing amplicons (middle), or that are predicted to be editable via prime editing by a common pegRNA (bottom). **e**, Dose–response curve of Cas9 mutagenesis with increasing doxycycline (dox)-induced Cas9 expression. Cells were treated with 0–50 ng/mL doxycycline for 3 days. Data points represent mean editing efficiency; error bars, s.e.m. **f**, LTR5 consensus sequence at the target site, with Cas9 target and pegRNA components labeled. Protospacer (green) and PAM (orange) are shown above; primer binding site (PBS) and reverse transcription template (RTT) are indicated below. The four candidate pegRNAs tested are shown with encoded edits (red, asterisks). pegRNAs #1–3 encode two substitutions; pegRNA #4 encodes a 2-nt insertion. **g**, Prime editing efficiency at LTR5-RGR1 targeted loci for the four pegRNAs compared to negative control (no pegRNA), 2 days after transient lipofection. Dots represent individual replicates; error bars indicate standard deviation (s.d.). **h**, Editing efficiency per LTR5-RGR1 target amplicon for pegRNA #1, #2, and #4. Each dot represents one amplicon. Error bars indicate standard error of the mean (s.e.m.). pegRNA samples (black); untreated samples (light blue, negative control). The y-axis shows the percentage of reads aligned to each amplicon that were prime edited. Red dashed line indicates 1% threshold for editing. **i**, Histogram of editing efficiency per amplicon for the pegRNAs showing non-zero editing: pegRNA #1 (blue), pegRNA #2 (pink), and pegRNA #4 (orange).

## Results

### LTR5-RGR1 as a prototype for recording in genomic repeats

Writing to multiple targets with a single gRNA and reading with a common primer pair would simplify expanding single-cell recording bandwidth (**Figure 1b**). To test if genomic repeats can offer such targets, we initially focused on Long Terminal Repeat 5 (LTR5, DF000000540), an endogenous retrovirus exclusive to Hominoidea (apes)^33,34^. We reasoned that LTR5’s relatively recent emergence in evolution, combined with its length of 969 base pairs and approximately seven thousand full or partial copies in the human genome would allow us to identify a Cas9 target with hundreds of copies and conserved flanking sequences for amplification with one pair of primers. We identified a Cas9-targetable sequence (protospacer and PAM) in LTR5 with over 600 copies across the haploid human genome (**Figure 1c**, **Table S1**). These copies contain highly conserved sequences upstream and downstream, enabling us to design a pair of primers capable of amplifying about half of the target sites (304 of 610) with only 33 off-target amplicons (**Figure 1c, 1d, Table S1**). Of these 304 targetable and amplifiable sites, 278 match the spacer sequence perfectly, 25 have one mismatch, and 1 has two mismatches (**Figure 1d**). Moreover, 77% of these sites (234 of 304) contain at least one unique sequence variant between the primer and gRNA target sites (**Figure 1d**), making them distinguishable in amplicon sequencing results (**Figure 1b**). We call the combination of this gRNA and amplification primers LTR5 Repeat for Genomic Recording 1, or LTR5-RGR1 for short.

To determine whether LTR5-RGR1 is accessible to Cas9 for editing, we transduced a previously established clonal HEK293T cell line containing stably-integrated doxycycline (Dox) inducible *S. pyogenes* Cas9^22^ with a lentivirus expressing the LTR5-RGR1 gRNA. Cas9 expression was then induced with a range of Dox concentrations, cells were harvested after four days, and the LTR5-RGR1 amplicon was sequenced with Illumina. LTR5-RGR1 editing displayed a sigmoidal relationship with Cas9 induction levels (**Figure 1e**), demonstrating both accessibility and a dynamic range suitable for distinguishing different signal intensities. However, prolonged Cas9 induction caused extensive cell death, consistent with the large number of LTR5-RGR1 targets (>600 per haploid genome) and the well-documented toxicity of numerous double-strand breaks (DSBs)^35,36^.

To avoid this toxicity, we switched to prime-editing. Prime editors (PEs) consist of a nicking Cas9 (nCas9) fused to a reverse transcriptase (RT) guided by a prime editing guide RNA (pegRNA) that both specifies the target site and encodes a desired edit as an RT template^37^. After nCas9 cuts one DNA strand, the RT uses the pegRNA template to synthesize new DNA containing the intended edit, which is then incorporated into the genome. This system enables precise edits without requiring double-strand breaks; however, it requires the pegRNA to include homology with the targeted DNA downstream of the PAM to allow the intended edit to be incorporated. We found that 90% of amplifiable LTR5-RGR targets (273 of 304) possessed the minimum sequence conservation for prime editing with a common pegRNA (**Figure 1d, Methods**), making LTR5-RGR1 a suitable target for multiplexed prime editing.

pegRNA performance varies significantly with the encoded edit^37,38^; therefore, we first compared four pegRNAs for LTR5-RGR1. All pegRNAs contain the same primer binding site (PBS) (**Figure 1f**), but each encodes a different two-base alteration that is expected to abrogate prime-editor binding by disrupting either the PAM or the seed sequence (**Figure 1f**). Constructs encoding these pegRNAs were transiently cotransfected with the Prime Editor 2 (PE2) system^37^ into HEK293T cells. 56 hours post-transfection, cells were harvested, LTR5-RGR1 was amplified with its primers, and sequenced with Illumina. pegRNA#1 showed the highest editing rate (16.5%), followed by pegRNA#2 (10.3%) and pegRNA#4 (7.9%); no edits were detected for pegRNA#3 (**Figure 1g**). We then assessed the distribution of these edits across amplicons with distinguishable sequences. Over 200 distinct amplicon sequences could be confidently identified. Editing levels varied across sequences (0–86% for pegRNA#1, 0–59% for pegRNA#2, 0–32% for pegRNA#4), but 91%, 86%, and 87% of sequences showed editing above 1%, indicating most sites are writable (**Figure 1h,i**). We chose pegRNA#1, which encodes the conversion of the ‘AGG’ PAM sequence into ‘ACC’, for further tests, as it showed the highest efficiency of editing in the largest number of sites. These results establish LTR5-RGR1 as a prototype RGR, enabling writing with a single guide and readout with a single primer pair across hundreds of target sites.

### A controllable prime-editing system to test recording performance

Testing LTR5-RGR1 recording performance requires tightly coupling prime editing to a controllable signal. We therefore evaluated several strategies for small-molecule induction of prime editor activity (**Figure 2a**). We first tested transcriptional control using a Dox-inducible TRE3G promoter driving PE2 expression (TRE-PE2) (**Figure 2a**). We generated a HEK293T cell line stably expressing LTR5-RGR1 pegRNA#1 via PiggyBac transposition and transiently transfected these cells with TRE-PE2, which carries a puromycin resistance cassette. After 48 hours of puromycin selection to remove untransfected cells, we induced with Dox for 72 hours. Amplicon sequencing of LTR5-RGR1 revealed that Dox induction achieved 12.2% editing on average, approaching constitutively expressed PE2 controls (**Figure 2b**). However, substantial background editing (8.3%) persisted without induction, yielding only ∼1.5-fold induction over background (**Figure 2c**).

**Figure 2:**
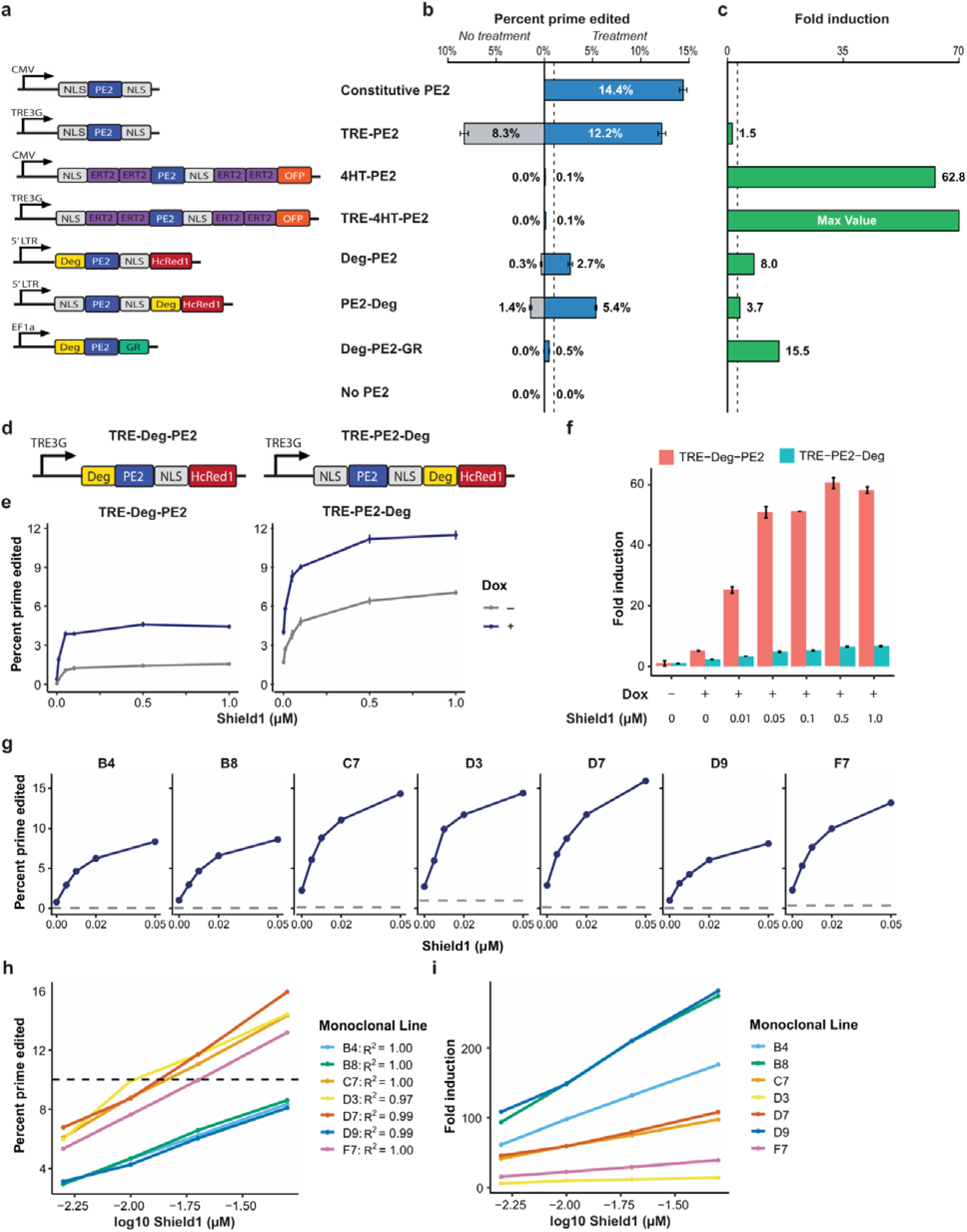
A tightly controllable dual-inducible prime editor. **a**, Schematic of inducible prime editor (PE2) constructs evaluated. Constructs were driven by constitutive (CMV, EF1a, *or 5’ UTR*) or Doxycycline-inducible (TRE3G) promoters. Fused regulatory and reporter domains include: nuclear localization signals (NLS), estrogen receptor T2 domains (ERT2), orange fluorescent protein (OFP), Shield1-responsive destabilizing domains (Deg), far-red fluorescent protein (HcRed1), and glucocorticoid receptor (GR). **b**, Prime editing efficiency of constructs shown in **a**. HEK cells were co-transfected with the indicated PE2 construct and LTR5-RGR1 pegRNA #1 and cultured without (gray bars) or with (blue bars) the corresponding inducer: 500 ng/mL doxycycline, 1 µM 4-hydroxytamoxifen (4HT), 1 μM Shield1, or 1 µM triamcinolone acetonide. Prime editing efficiency (percentage of LTR5-RGR1 target containing reads that had the encoded GG to CC substitution) was quantified by high-throughput sequencing 72 h post-transfection. Dashed line indicates 1% editing threshold. Error bars, s.e.m. (*n = 2* biological replicates). **c**, Fold induction for each inducible construct, calculated as the ratio of editing in treated versus untreated conditions. TRE-4HT-PE2 exhibited undetectable basal editing, yielding a maximum signal-to-noise ratio (denoted "Max Value"). Dashed line indicates signal-to-noise threshold of 3. **d,** Schematic of dual-inducible prime editor constructs. Deg-PE2 and PE2-deg from **a** were cloned into PiggyBac vectors under the TRE3G promoter, generating TRE-Deg-PE2 and TRE-PE2-Deg. **e,** Prime editing dose-response of dual-inducible constructs across Shield1 concentrations, in the presence (navy) and absence (gray) of 500 ng/mL doxycycline. HEK cell lines stably expressing the indicated PE2 construct and LTR5-RGR1 pegRNA #1 were induced for 72 h before sequencing. Error bars, s.e.m. (n = 2 biological replicates). **f,** Fold induction of TRE-Deg-PE2 (red) and TRE-PE2-Deg (turquoise) across induction conditions, calculated as the ratio of percent prime edited in the indicated treatment condition over the fully uninduced baseline condition (no Doxycycline, 0X Shield1). Dots represent individual replicates; error bars, s.e.m. (*n = 2* biological replicates). **g**, Prime editing dose-response of individual monoclonal HEK cell lines (B4, B8, C7, D3, D7, D9, F7) stably expressing TRE-Deg-PE2 and LTR5-RGR1 pegRNA #1 across Shield1 concentrations in the presence of 500 ng/mL doxycycline. Gray dashed line indicates baseline editing in the fully uninduced condition (no doxycycline, 0 μM Shield1). Error bars, s.e.m. **h**, Prime editing dose-response of monoclonal lines from **g** with data plotted against log₁₀ Shield1 concentration. Cells were cultured with 500 ng/mL doxycycline and the indicated Shield1 concentration for 4 days, with daily treatment media preparation and replacement. R^2^ values indicate goodness of fit for linear regression of editing efficiency against log10 Shield1 concentration. Dashed line indicates 10% editing threshold for clone selection. **i**, Fold induction of monoclonal cell lines across induction conditions. Fold induction again calculated as the ratio of editing in the indicated treatment condition over the fully uninduced baseline condition.

We next tested post-translational control through inducible nuclear localization. Fusing PE2 to estrogen receptor ligand-binding domains (ERT2) (**Figure 2a**) restricts nuclear entry until 4-hydroxytamoxifen (4HT) exposure. However, both constitutive (4HT-PE2) and Dox-inducible (TRE-4HT-PE2) versions performed poorly, neither exceeding 1% editing upon induction, despite high fold-change over background (**Figure 2b,c**).

We then evaluated protein stability control using an FKBP12-derived degron that is stabilized by the synthetic ligand Shield1^39^ (**Figure 2a**). Fusing this degron to either the N-terminus (Deg-PE2) or C-terminus (PE2-Deg) of the prime editor yielded promising results: 2.7% and 5.4% editing upon Shield1 induction, respectively, with at least fivefold lower editing when uninduced—yielding acceptable fold induction for both configurations (**Figure 2b,c**). Adding glucocorticoid receptor-mediated nuclear localization (Deg-PE2-GR) further improved fold induction upon double-induction with Shield1 and triamcinolone acetonide but yielded editing below 1% after induction (**Figure 2a–c**). Based on these results, we selected Shield1 as our recording signal and proceeded to optimize the degron-based constructs.

To further reduce leakage, we cloned the Deg-PE2 and PE2-Deg designs into PiggyBac vectors under the TRE3G Dox-inducible promoter, creating TRE-Deg-PE2 and TRE-PE2-Deg constructs (**Figure 2d**). After transposition into the pegRNA#1-expressing HEK293T cells and selection to generate stable lines, we tested a range of Shield1 concentrations with and without Dox induction for 3 days (**Figure 2e,f**). While TRE-PE2-Deg achieved higher absolute editing levels, TRE-Deg-PE2 exhibited superior fold induction along both axes: between Dox-induced and uninduced conditions at matched Shield1 concentrations (e.g., 2.8-fold versus 1.7-fold at the highest Shield1), and between the highest and lowest Shield1 concentrations in the presence of Dox (12-fold versus 3-fold) (**Figure 2e**). Combined, these differences yielded 58-fold total induction for TRE-Deg-PE2 (maximum Shield1 with Dox versus no Shield1 without Dox) compared to 7-fold for TRE-PE2-Deg (**Figure 2f**). We therefore selected TRE-Deg-PE2 for its superior fold induction and near-complete absence of background editing when both inducers were withheld.

Because the transposed cell population is heterogeneous^40^ with respect to TRE-Deg-PE2 and pegRNA#1 copy number and integration loci, factors that could produce non-uniform Shield1 responses, we derived seven monoclonal lines. To characterize the dose-response of each clone, we exposed them to a range of low Shield1 concentrations (0 μM, 0.005 μM, 0.01 μM, 0.02 μM, and 0.05 μM) with Dox for four days and quantified LTR5-RGR1 editing by amplicon sequencing. All clones showed dose-dependent editing across the tested Shield1 concentrations in the presence of Dox (**Figure 2g**) with a log-linear relationship that enables analog recording capable of distinguishing inputs across an order of magnitude (**Figure 2h**). However, clones varied considerably in absolute editing levels and fold induction (**Figure 2h,i**). Clones with the highest fold induction (i.e., B4, B8, D9) tended to have low absolute editing, limiting their utility in practice. Clone D7 offered a favorable balance: the highest editing levels across Shield1 concentrations (15.9% at highest induction), negligible background without treatment (<0.15%), a log-linear response (R^2^ = 0.99), and fold induction exceeding 100-fold at 0.05 μM Shield1 versus no Dox (**Figure 2g–i**). This clone provides a tightly controllable system for evaluating LTR5-RGR1 recording performance.

### High-resolution recording in LTR5-RGR1 in cell populations

To test RGR recording performance, we exposed the D7 monoclonal Shield1-inducible prime editing line with LTR5-RGR1 pegRNA#1 to five different Shield1 levels (0, 0.005, 0.01, 0.02, and 0.05 µM) over five weeks with Dox, harvesting an aliquot of cells every 3 to 6 days and sequencing their LTR5-RGR1 amplicon (**Figure 3a**). A sample without Dox or Shield1 served as a negative control. We quantified RGR mutagenesis as the fraction of protospacer-containing reads carrying the exact edit specified by pegRNA#1. Editing increased with time across all induction levels, and remarkably, LTR5-RGR1 editing rates distinguished twofold differences in Shield1 concentration after only three days of exposure (**Figure 3b**). As a measure of resolution, we calculated the standardized effect size (Cohen’s *d*) between adjacent Shield1 concentrations (**Figure 3c**). Cohen’s *d* ranged from 2.5 to 69, indicating complete separation of adjacent conditions. These results demonstrate that LTR5-RGR1 offers high-resolution signal discrimination.

**Figure 3:**
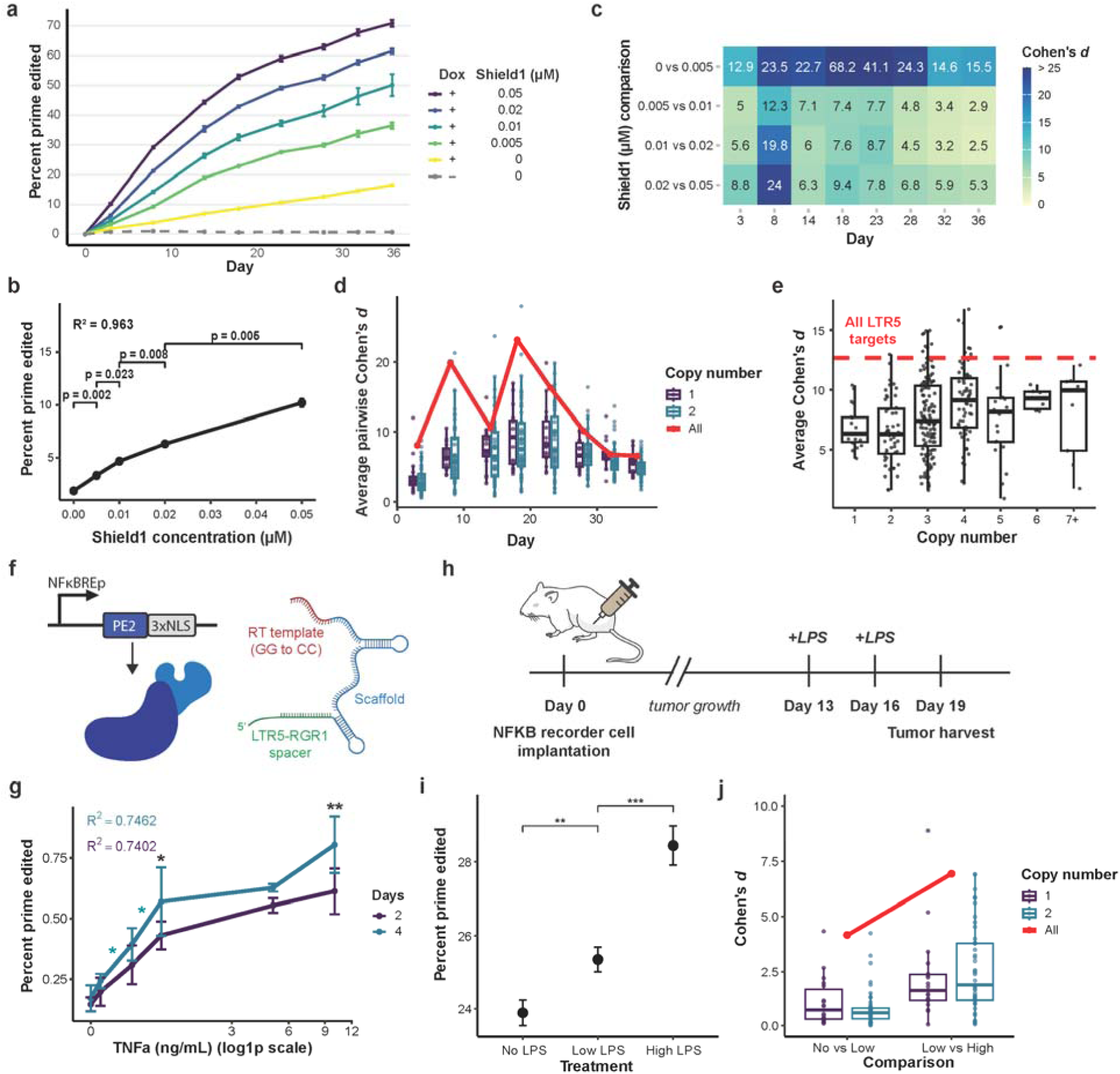
Analogue recording in cell populations using LTR5-RGR1. **a**, Prime editing accumulation over a 36 day time course. Monoclonal HEK cells (line D7) stably expressing TRE-Deg-PE2 and LTR5-RGR1 pegRNA #1 were treated with 500 ng/ml doxycycline and varying Shield1 concentrations (0 μM, 0.005 μM, 0.01 μM, 0.02 μM, and 0.05 μM), or left untreated (negative control). Treatment media was prepared and refreshed daily. Cells were harvested every 3-6 days and LTR5-RGR1 amplicons were sequenced. The y-axis shows the percentage of protospacer-containing reads with the encoded GG to CC substitution. Data points represent mean ± s.e.m. (*n = 3* biological replicates). **b**, Prime editing in response to Shield1concentration at day 3 (data from **a**). Data points represent mean editing efficiency; error bars, s.e.m. (*n = 3* biological replicates). R² value indicates goodness of fit for linear regression of dose-response relationship. Statistical significance between successive Shield1 concentrations was assessed using two-sided Student’s t-tests; *P* values are shown above brackets. **c**, Heatmap of Cohen’s *d* effect sizes for comparisons between adjacent Shield1 concentrations across time points. Rows represent pairwise concentration comparisons; columns represent measurement days. Cohen’s *d* values are displayed within each cell. **d**, Average Cohen’s *d* for pairwise comparisons between treatment conditions over time for single- and double-copy amplicons versus for using all LTR5-RGR1 targets. Single-copy amplicons (purple), double-copy amplicons (turquoise), and all LTR5-RGR1 targets combined (red line) are shown. Box plots: center line, median; box bounds, first to third quartiles (25th–75th percentiles); whiskers, 1.5× interquartile range. **e**, Average Cohen’s *d* for pairwise comparisons between treatment conditions across all time points, stratified by amplicon copy number. Red dashed line indicates average Cohen’s *d* for using all LTR5-RGR1 targets. Box plots are as described in **d**. **f**, Schematic for NF-κB PE2 recorder cell line. Inside each cell, is a NF-κB induced PE2 and a LTR5-RGR1 pegRNA encoding the CC to GG edit. **g**, Dose-dependent prime editing following TNF-α treatment in vitro. HEK cells expressing the NF-κB-inducible prime editing recorder and LTR5-RGR1 pegRNA #1 were treated with varying concentrations of TNF-α for 2 days (purple) or 4 days (turquoise). The y-axis indicates the percentage of LTR5-RGR1 amplicons containing the encoded edit. Data points represent mean ± s.e.m. R^2^ values indicate goodness of fit for linear regression of editing efficiency against log10 TNF-α concentration for day 2 (purple) and day 4 (turquoise). Black asterisks denote significant differences in editing efficiency between day 2 and day 4 at a given concentration; turquoise asterisks denote significant differences between consecutive TNF-α concentration treatments within day 4 samples. Statistical comparisons were performed using pairwise contrasts derived from a global linear model, with standard errors reflecting pooled variance across the entire dataset. **h**, Experimental design for in vivo recording of LPS-induced inflammation. Mice were xenografted with NF-κB recorder cells on day 0 and injected with low-dose (0.03 mg/kg) or high-dose (3 mg/kg) LPS on days 13 and 16. Tumors were harvested on day 19. **i**, Dose-dependent prime editing following LPS administration in vivo. The y-axis indicates the percentage of LTR5-RGR1 amplicons containing the encoded edit. Data points represent mean ± s.e.m. (*n = 2* tumors per condition, from one mouse per treatment group). Asterisks denote statistical significance between consecutive treatment conditions. **j**, Cohen’s *d* effect sizes for consecutive treatment comparisons (No LPS versus Low LPS; Low LPS versus High LPS), stratified by copy number. Single-copy amplicons (purple), double-copy amplicons (turquoise), and all LTR5-RGR1 targets combined (red points with connecting line) are shown. Box plots: center line, median; box bounds, 25th–75th percentiles; whiskers, 1.5× interquartile range. **g,i**, \**P* < 0.05, \*\**P* < 0.01, \*\*\**P* < 0.001 (**g**, pairwise contrasts from global linear model; **i**, two-sided Student’s t-test).

We next examined how individual LTR5-RGR1 copies contribute to recording performance. Using single-cell amplicon sequencing, we estimated the genomic copy number of each unique LTR5-RGR1 amplicon sequence in the D7 line (see below, **Table S2** and **Methods**). Each unique amplicon sequence is identified by sequence variation(s) around the protospacer (**Figure 1b**). We then compared the signal resolution using all LTR5-RGR1 amplicons versus that of amplicons with one or two copies in the genome. The single-copy sequences are comparable to an engineered landing pad inserted in one of the two homologous chromosomes^3,11,13,14,41^. The double-copy sequences are comparable to a native genomic site (e.g., a safe harbor) that is naturally in a diploid state^16,42^. For each subset (all sites, single-copy, and double-copy), we calculated the average Cohen’s *d* between adjacent Shield1 concentrations at each timepoint (**Figure 3d**). Combining all LTR5-RGR1 sequences yielded substantially better signal resolution, on average 2-fold higher than either single- or double-copy subsets alone. Amplicon sequences with even higher copy numbers improved resolution but did not exceed the resolution achieved using all sequences combined (**Figure 3e**).

Further analysis of resolution across exposure time revealed that the advantage of combining LTR5-RGR1 sequences emerges from their heterogeneous editing kinetics. Different amplicon sequences mutate at characteristic rates, creating distinct recording channels. Fast-editing amplicons better distinguish concentration differences over short exposures but saturate with prolonged exposure, losing discriminatory power (**Figure S1a**). Conversely, slow-editing amplicons better resolve differences over long exposures but accumulate insufficient edits after short exposures to distinguish conditions (**Figure S1b**). By combining sites with diverse kinetics, the RGR captures differences across a wide temporal dynamic range. These results demonstrate that recording in genomic repeats achieves finer signal discrimination across a broader dynamic range than conventional approaches.

### RGR-based recording of inflammatory signaling in vivo

The preceding results establish LTR5-RGR1 performance with a synthetic signal in vitro. Before testing LTR5-RGR1 in vivo, we first validated its ability to record a natural biological signal in cultured cells. We coupled PE2 expression to an NFκB-responsive promoter (NFκB-PE2), which combines tandem NFκB response elements with a minimal promoter to drive expression in response to the inflammatory cytokine tumor necrosis factor-α (TNF-α)^3^. We generated a HEK293T line with both NFκB-PE2 and LTR5-RGR1 pegRNA#1 stably integrated (**Figure 3f**) and exposed it to a range of TNF-α concentrations, sequencing LTR5-RGR1 at days 2 and 4 (**Figure 3g**). The system exhibited a log-linear dose-response between 0 and 10 ng/ml TNF-α and distinguished 2-day from 4-day exposures across the concentration range (marginal means *t*-test, *p* < 0.001, **Figure 3g**), confirming its ability to record analog inflammatory responses via NFκB activity.

We then tested this inflammation recorder in a xenograft model. We implanted the NFκB recording line bilaterally via subcutaneous injection into the flanks of three immunodeficient mice. We induced acute inflammation via intraperitoneal lipopolysaccharide (LPS) injection on days 13 and 16 (**Figure 3h**). LPS activates Toll-like receptor 4 (TLR4) on host immune cells and triggers release of circulating cytokines including TNF-α^43,44^. Mice received either low-dose LPS (0.03 mg/kg), high-dose LPS (3 mg/kg), or saline control for both injections (one mouse per condition). Mice were euthanized on day 19 and tumors were harvested (one injection site in the saline-treated mouse failed to engraft; two distinct pieces of its only tumor were used as replicates). We amplified LTR5-RGR1 from each tumor. Because the amplification primers are specific to human LTR5, which is absent from the mouse genome, no normalization for tumor fraction was required. The results revealed clear dose-dependent accumulation of edits in LTR5-RGR1 (**Figure 3i**), demonstrating successful in vivo recording of inflammatory signaling in a genomic repeat. As with our in vitro analysis, we compared signal discrimination of all LTR5-RGR1 sequences to single- and double-copy subsets (**Figure 3j**). The average Cohen’s *d* between adjacent treatment groups (saline vs. low-dose, low-dose vs. high-dose) for all amplicons was 2.9-fold higher than double-copy amplicons (one-sample t-test, p < 0.001) and 3.3-fold higher than single-copy sequences (p < 0.001). These results demonstrate that the resolution and dynamic range advantages of recording in repeats over conventional single- or double-site approaches extend to physiological signals in vivo.

### RGR-based recording enables single-cell signal classification

Single-cell recording of multiple signaling states has remained a significant challenge in genomic recording. Current approaches require at least 50 cells to distinguish binary states (signal ON vs. OFF)^9,42^ and thousands of cells to resolve multi-state or analog signals^3,14,16,45,46^. We reasoned that by expanding the writing space per cell, genomic repeats could enable multi-state recording at single-cell resolution.

To test this, we treated the D7 monoclonal line (carrying integrated TRE-Deg-PE2 and LTR5-RGR1 pegRNA#1; **Figure 2g,h**) with four signal levels for 23 days: no Dox or Shield1 (OFF state), or 1 µg/ml Dox with 0, 0.01, or 0.05 µM Shield1 (low, mid, and high ON states, respectively) (**Figure 4a**). We sorted 270 single cells from each treatment group and sequenced LTR5-RGR1 amplicons (**Figure S2a**). The distribution of editing levels across treatment groups showed significantly different means (two-sided Wilcoxon rank-sum test, *p* < 0.0001) but high variance (**Figure 4b**). This variance reflects both intrinsic noise in the recording system and the extensive cell-to-cell heterogeneity inherent in gene expression and cellular responses^47–49^.

**Figure 4:**
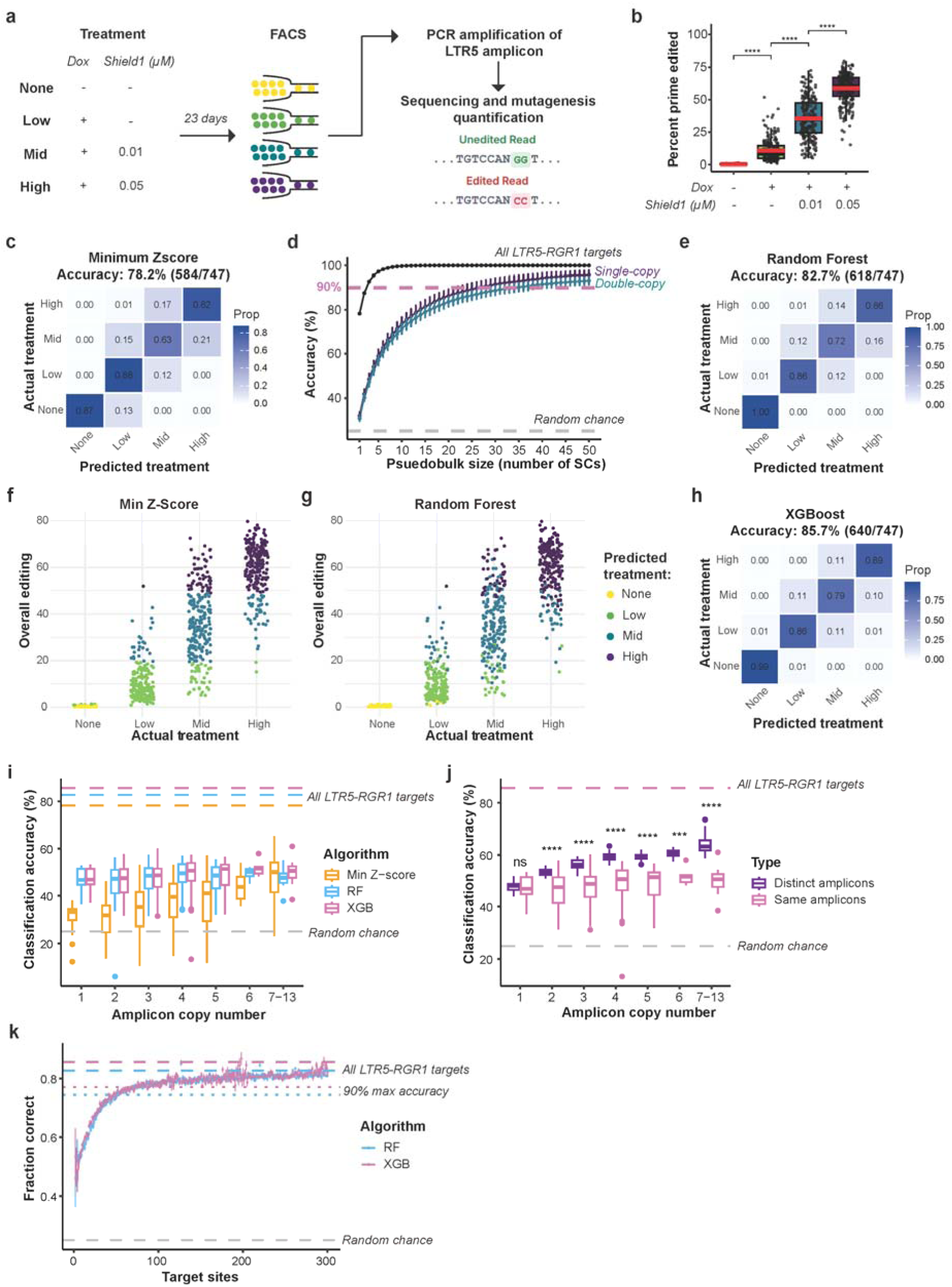
Single-cell recording and classification using LTR5-RGR1. **a**, Experimental layout for single-cell recording. Monoclonal HEK cells (line D7) stably expressing TRE-Deg-PE2 and LTR5-RGR1 pegRNA #1 were treated with 500 ng/mL doxycycline and one of three Shield1 concentrations (0 μM, 0.01 μM, or 0.05 μM), or left untreated (negative control), for 23 days to generate four distinct signal levels(None, Low, Mid, and High). Treatment media was prepared and refreshed daily. Single cells were isolated by FACS, followed by PCR amplification of LTR5-RGR1 loci, sequencing, and prime editing quantification. **b**, Prime editing across single cells in the four treatment groups (∼250 cells per treatment group). The y-axis indicates the percentage of protospacer-containing reads with the encoded GG to CC substitution. Box plots: red line, mean; black line, median; box bounds, 25th–75th percentiles; whiskers, 1.5× interquartile range. \*\*\*\**P* < 0.0001 for all pairwise comparisons (two-sided Wilcoxon rank-sum test). **c**, Confusion matrix for single-cell classification using the naive minimum Z-score framework. Rows indicate actual treatment group; columns indicate predicted treatment group. Cells are colored by the proportion of cells in each category. **d**, Classification accuracy as a function of the number of pooled single cells using all LTR5-RGR1 target sites (black), single-copy amplicons (purple), or double-copy amplicons (turquoise). Pink dashed line: 90% accuracy threshold; gray dashed line: 25%, random chance. Data represent mean ± s.e.m. (error bars may be smaller than symbols in some cases). **e**, Confusion matrix for single-cell classification using Random Forest. Overall accuracy: 82.7% (618/747 cells). **f**,**g** Single-cell prime editing levels colored by predicted treatment group (yellow: None; green: Low; blue: Mid; purple: High) using (**f**) the naive minimum Z-score framework or (**g**) Random Forest. **h**, Confusion matrix for single-cell classification using XGBoost. Overall accuracy: 85.7% (640/747 cells). **i**, Single-amplicon classification accuracy stratified by copy number. Box plots display classification accuracy when individual amplicons are used as single-feature predictors, evaluated using the naive minimum Z-score (orange), Random Forest (blue), or XGBoost (pink) frameworks. Each point represents the test accuracy of a single amplicon via leave-one-out cross-validation (Random Forest, XGBoost) or distance to global mean (Z-score). Dashed horizontal lines indicate accuracy achieved using all 291 amplicons (Z-score: 78.2%; Random Forest: 82.7%; XGBoost: 85.7%). Gray dashed line indicates random classification (25%). **j**, Comparison of classification accuracy using single amplicons with increasing copy number versus multiple distinct single-copy amplicons. Box plots display XGBoost classification accuracy using either individual amplicons with copy number *n* (pink) or *n* distinct single-copy amplicons (purple). For single amplicons with increasing copy number, each point represents the test accuracy via leave-one-out cross-validation. For multiple single-copy amplicons, each point represents the mean accuracy across 1,000 simulations in which *n* single-copy amplicons were randomly sampled for classification. Pink dashed line indicates accuracy achieved using all 291 amplicons (85.7%); gray dashed line indicates random classification (25%). \**P* < 0.05, \*\**P* < 0.01, \*\*\**P* < 0.001, \*\*\*\**P* < 0.0001 (two-sided Wilcoxon rank-sum test); ns, not significant. **k**, Classification accuracy as a function of cumulative copy number. Simulated classification accuracy of Random Forest (blue) and XGBoost (pink) models using randomly sampled subsets of LTR5-RGR1 amplicons. The x-axis indicates cumulative copy number of the randomly drawn amplicon subset; the y-axis indicates classification accuracy. Error bars represent standard error of the proportion. Dashed horizontal lines indicate maximum accuracy achieved using all 291 amplicons (Random Forest: 0.827; XGBoost: 0.857). Dotted horizontal lines indicate 90% of maximum accuracy. Gray dashed line indicates random classification (0.25).

We first evaluated single-cell classification accuracy with a naive approach: using the editing rate distribution of each treatment group as a reference and assigning each cell to the group with the lowest z-score distance (i.e., fewest standard deviations from the group mean). This approach correctly classified 78.2% of cells into their respective treatment groups (**Figure 4c**). Notably, this accuracy was achieved in single cells—a regime where prior methods lack discriminatory power. For the simplified binary case of distinguishing OFF from the high ON state (analogous to standard benchmarks in the field), accuracy reached 100% (**Figure 4c**). These results demonstrate that RGRs enable multi-state signal classification at single-cell resolution.

To compare RGR-based recording with conventional single- or double-site approaches, we estimated the genomic copy number of each LTR5-RGR1 amplicon sequence from the single-cell data. Briefly, we used the per-amplicon editing rates observed across single cells from each treatment group to estimate the probability that each copy of an amplicon would be edited in a given treatment group. We then assigned each amplicon the copy number whose expected editing distribution best matched the observed distribution across cells, accounting for estimated dropout rates (**Table S2**, **Methods**). For example, amplicon sequences observed exclusively as fully edited or unedited across all cells (i.e., never partially edited) were designated as single-copy. These amplicons likely represent copies with unique variation present on one chromosome only. Conversely, increasing copy number leads to a proportional increase in partially edited cells. The total copy number of LTR5-RGR1 targets in the D7 line was estimated at 1,120, within the expected range given our estimate of 304 targets in the haploid reference genome (**Figure 1c**) and the hypotriploid karyotype of HEK293T cells^50^. Using these estimates, we performed classification using individual single- or double-copy amplicons, finding that they accomplished only 32% and 31% accuracy, respectively—comparable to the 25% expected by chance.

To evaluate how classification improves when combining multiple cells, we generated pseudobulk samples by randomly pooling 2–50 cells within each treatment group (**Figure 4d**). Single- and double-copy amplicons required 13 and 15 cells, respectively, to match the single-cell classification accuracy of all LTR5-RGR1 amplicons (78.2%). To reach 90% accuracy, they required over 40 and 50 cells, respectively (**Figure 4d**). By contrast, classification using all LTR5-RGR1 amplicons exceeded 90% accuracy with just three cells. These results underscore the substantial advantage of RGR-based recording over single- and double-site alternatives, achieved while maintaining the simplicity of a single guide RNA and primer pair.

We next asked whether machine learning approaches that leverage the mutational states of individual amplicons could improve classification accuracy. The naive approach provides a robust baseline but fails to account for the differential recording kinetics of individual LTR5-RGR1 sites: some respond sensitively to low signals and saturate early, while others record more slowly (**Figure S1a,b**). We first implemented a Random Forest (RF) classifier^51^ that weighs the mutational state of 304 distinguishable amplicon sequences, improving accuracy from 78.2% to 82.7% (**Figure 4e**). The naive approach systematically misclassifies cells at the tails of overlapping editing distributions (**Figure 4f**), but the RF model correctly classified many of these outliers by recognizing amplicon-specific patterns, and eliminated false positives by achieving 100% accuracy for the OFF group (**Figure 4g**). However, the RF model also showed signs of overfitting to dominant features: removing the six most informative amplicons paradoxically improved accuracy to 83.0% (**Figure S2b**), suggesting these dominant features, which largely recapitulated overall editing, were obscuring subtler discriminatory signals across the remaining amplicons.

To better leverage information across all amplicons, we implemented an XGBoost (Extreme Gradient Boosting) framework^52^. Using a lower feature sampling rate to reduce the influence of dominant amplicons, this approach achieved 85.7% accuracy, with notable gains in distinguishing mid from high treatment groups (**Figure 4h**). Classification accuracy across the three approaches correlated with confidence separation between correct and incorrect predictions (Cohen’s *d*: naive = 0.38; RF = 1.08; XGBoost = 1.19; **Figure S2c-e**), indicating that models leveraging high-dimensional amplicon information not only classify more accurately but also better distinguish reliable from unreliable predictions. Moreover, coverage variation across cells did not substantially affect prediction accuracy for any model (**Figure S2f–k**), demonstrating RGR robustness to partial coverage and dropout, an inevitable challenge in single-cell analysis.

We next examined how classification accuracy scales with amplicon copy number. Using our copy number estimates, we repeated Z-score, RF, and XGBoost classification for each single cell, considering only amplicons with 1–13 genomic copies (**Figure 4i**). Classification accuracy increased with copy number (Spearman’s ρ = 0.303, 0.126, and 0.195 for Z-score, RF, and XGBoost, respectively), but gains were modest and average accuracy remained below 51% across all methods. To distinguish the contributions of redundancy versus diversity, we performed an additional analysis using the best-performing XGBoost classifier: we compared classification accuracy using 1–13 copies of a single amplicon versus 1–13 different single-copy amplicons (**Figure 4j**). While neither approach matched classification with the full RGR, two or more diverse single-copy amplicons outperformed equivalent numbers of identical copies by an average of 10.04 percentage points. These results demonstrate that the advantage of RGR-based recording over single-copy and diploid sites is substantial at single-cell resolution, deriving not from redundancy alone but from sequence diversity as well.

We next asked how many genomic sites, regardless of sequence distinguishability, are required for high-resolution recording. We performed classification using randomly sampled subsets of the 304 distinct amplicons, with combined genomic copy numbers ranging from 1 to 300, across 10,000 iterations per condition (**Figure 4k**). Classification accuracy increased rapidly with cumulative copy number, surpassing 75% with as few as 52 copies. An inflection point at approximately 100 copies marked where 90% of maximum accuracy was achieved for both RF and XGBoost. Beyond this threshold, gains were marginal, with accuracy asymptotically approaching the maximum. These results suggest that genomic repeats with 50 or more copies provide substantial benefit over conventional approaches, while approximately 100 copies capture most of the benefit of RGR-based recording.

Overall, these results establish that recording in genomic repeats enables accurate single-cell classification into multiple signaling states. The diversity of RGR kinetics provides higher resolution than conventional single- or double-site approaches, while the redundancy of recording across hundreds of sites confers robustness to the inherent stochasticity and incompleteness of molecular and sequencing readouts.

### Systematic identification of RGRs across species

While LTR5-RGR1 demonstrated how recording in genomic repeats increases resolution and enables single-cell recording, it is unlikely to support all applications. Some copies of LTR5-RGR1 overlap coding sequences (**Table S1**), and widespread adoption would require RGRs that avoid functional sequences. Moreover, LTR5-RGR1 is absent from common model organisms such as mouse and zebrafish. To assess the broader viability of RGR-based recording, we developed a computational pipeline (**Figure S3**) to systematically identify candidate RGRs that: (1) contain a common CRISPR target, (2) avoid functional sequences, and (3) share primer binding sites for amplification.Our pipeline begins with RepeatMasker annotations of repeats within the human, mouse, and zebrafish genomes (**Figure S3a**). RepeatMasker identifies repeat coordinates and classifies them into families^53^. For each genome, we counted distinct *S. pyogenes* Cas9 targets (20-nucleotide protospacer sequences adjacent to an NGG PAM), limiting the search to targets with up to 200 haploid copies (400 diploid) and moderate or better predicted gRNA effectiveness (≥0.55 CRISPRater score)^54^. This step identified 2,068,178 (human), 1,421,442 (mouse), and 1,980,974 (zebrafish) gRNAs that perfectly match multiple copies of a genomic repeat in each haploid genome (**Table S3**).

This initial count establishes a lower bound for target number as each gRNA may also target sites outside annotated interspersed repeats. Additionally, Cas9 tolerates up to two mismatches between gRNA and target depending on their location^55–60^. To identify all likely targets, we scanned each genome for each Cas9 target using CasOFFinder^61^ (**Figure S3b**), allowing up to two mismatches within the protospacer but not within the six-nucleotide seed sequence or the PAM, where mismatches are poorly tolerated^55,56^. This genome-wide scan increased median target counts 2.5-, 5.25-, and 2.75-fold for human, mouse, and zebrafish gRNAs, respectively. It also revealed exonic targets—an unambiguous indication of potential functional interference. We thus eliminated gRNAs with any target in or within 20 bases of an exon (**Table S3**), retaining those with 20–200 total genome-wide copies exclusively outside exons in each haploid genome. This filtering yielded 211,814 (human), 231,152 (mouse), and 287,438 (zebrafish) non-exonic gRNAs with 20–200 targets in each haploid genome (**Table S3**). **Figure 5a** shows the target count distribution of these gRNAs, which we term repeat-targeting guide RNAs (rgRNAs). We chose 20 haploid copies as a lower bound because our single-cell classification analysis showed that RGR advantages can be realized with approximately 50 genomic copies (**Figure 4j**); a gRNA matching 20 targets in a haploid reference genome would target 40 locations in a diploid cell.

**Figure 5:**
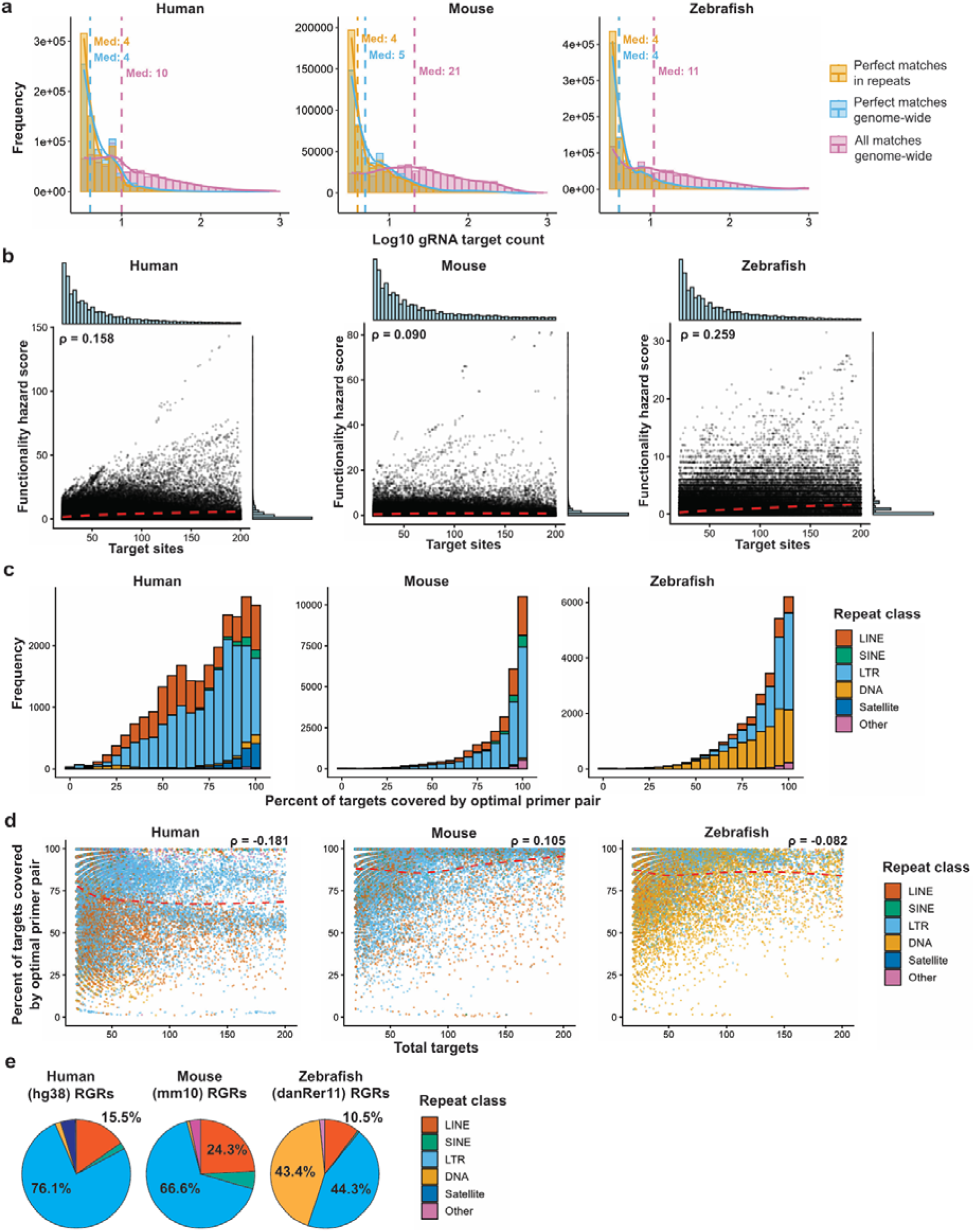
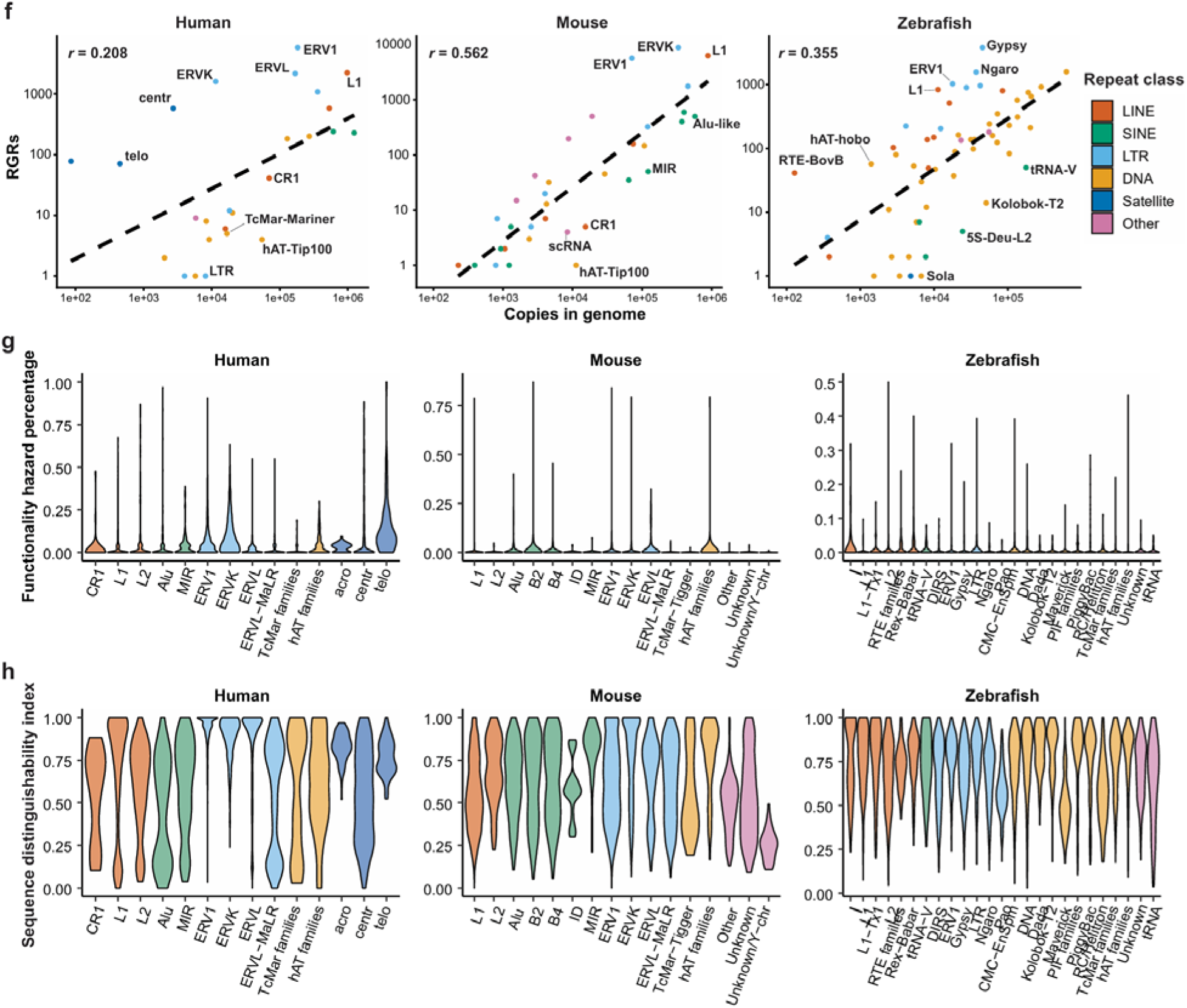
RGR properties across species. **a**, Distribution of target sites per non-exonic rgRNA in human, mouse, and zebrafish. Histograms show the frequency of target site counts (log10 scaled) quantified by three metrics: perfect matches strictly within annotated repeats (orange), perfect matches genome-wide (blue), and matches allowing up to two non-seed mismatches genome-wide (green). Vertical dashed lines indicate median values for each category. **b**, Relationship between functionality hazard score and number of sites targeted for non-exonic rgRNAs with 20-200 targets. Scatter plots display each rgRNA’s total target count (x-axis) versus calculated functionality hazard score (y-axis). Marginal histograms show the frequency distributions of both variables. Red dashed lines represent LOESS regression fits. Spearman’s ρ is indicated for each species. **c**, Distribution of optimal primer pair coverage stratified by repeat class. Stacked histograms show the frequency of rgRNAs across increasing primer coverage bins, colored by repeat class. **d**, Optimal primer pair coverage versus target site count for non-exonic rgRNAs. Scatter plots show the percentage of target sites expected to be recovered by the pipeline-identified optimal primer pair (y-axis) versus total sites targeted (x-axis). Data points are colored by their corresponding repeat class: LINE (red-orange), SINE (green), LTR (light blue), DNA (yellow), Satellite (dark blue), and Other (pink). Spearman’s ρ is indicated for each species. **e**, Distribution of RGRs across human (hg38), mouse (mm10), and zebrafish (danRer11) repeat classes. Pie charts show the relative proportion of each repeat class comprising total RGR content. **f**, Correlation between repeat copy number and RGR number across repeat families. Scatter plots show the relationship between repeat family copy number (x-axis) number of RGRs (y-axis) for human (left), mouse (middle), and zebrafish (right). Both axes are log10-scaled. Data points represent individual repeat families, colored by repeat class (LINE, SINE, LTR, DNA, Satellite, and Other). Black dashed lines indicate linear regression fits. Pearson’s *r* is indicated for each species. **g**, Distribution of functionality hazard percentage across repeat families. Violin plots show the distribution of functionality hazard percentage (y-axis) for rgRNAs within individual repeat families (x-axis), colored by repeat class (LINE = red-orange, SINE = green, LTR = light blue, DNA = yellow-orange, Satellite = dark blue, Other = pink, in listed order). **h**, Distribution of sequence distinguishability index (*SDI*) across repeat families. Violin plots show the distribution of sequence distinguishability index (y-axis) for rgRNAs within individual repeat families (x-axis), colored by repeat class. Higher values indicate greater sequence diversity among target amplicons, enabling more reliable locus-level resolution. **g-h**, Repeat families are ordered along the x-axis within each class alphabetically.

Beyond exons, defining sequence functionality is more complex and context-dependent, as the functional relevance of target loci will vary by cell type and application. To broadly assess potential interference with active regulatory elements, we compared rgRNA target coordinates to known promoter regions and candidate cis-regulatory elements (cCREs) from ENCODE for human and mouse^62^, and DANIO functional annotations for zebrafish^63^. We developed a functionality hazard score to quantify this risk. Targets overlapping promoter-like signatures (PLS), proximal or distal enhancer-like signatures (pELS/dELS), or potential promoter regions (500 bp upstream of a transcription start site) were assigned a weight of 1. Remaining targets overlapping DNaseI hypersensitive sites, H3K4me3-marked regions, or CTCF binding sites were assigned a weight of 0.5. These weights were summed across all targets to generate the functionality hazard score. For zebrafish, we assigned a weight of 1 to DANIO annotated consensus promoters, validated enhancers, and Consensus Predicted ATAC-seq-supported Developmental Regulatory Elements (cPADREs)^63^. Remaining targets overlapping DANIO Dynamic Orphan Predicted Elements (DOPEs) or Constitutive Orphan Predicted Elements (COPEs) were assigned a weight of 0.5. The functionality hazard score (sum of weights across targets) and functionality hazard percentage (score divided by target number) together quantify the absolute and relative potential for interference with functional elements, respectively.

As expected, the functionality hazard score correlated positively with target number across all three genomes (**Figure 5b**); however, this correlation was weak (Spearman’s ρ < 0.26) and the functionality hazard percentage remained steady across target counts (–0.10 ≤ ρ ≤ 0.13; **Figure S4a**). These observations indicate that rgRNAs with more targets are not disproportionately enriched for functional elements. Moreover, a substantial fraction of rgRNAs maintained low scores (**Figure S4b**), with 33.1% (human), 70.0% (mouse), and 62.6% (zebrafish) having a hazard score of 0, providing a pool of candidates for recording applications in each genome.

We next sought to identify conserved primer binding sites for amplification of rgRNA targets. Because many rgRNAs differ only by small shifts in spacer position and would share the same primers, we first merged rgRNAs with >60% target overlap (defined as ≥16 nucleotides of protospacers overlapping) into groups. This consolidation reduced the number of unique rgRNAs from 211,814 to 168,743 (human), 231,152 to 173,196 (mouse), and 287,438 to 112,092 (zebrafish). For each merged group, we selected a representative rgRNA by minimizing functionality hazard percentage while maximizing exact-match target count (**Table S3**, **Methods**).

We then assessed whether conserved primer binding sites exist for each unique rgRNA. We performed Clustal Omega multiple sequence alignment^64^ on a 520 bp window centered on each rgRNA’s targets and required a minimum of 15 bp conservation on both sides of the spacer across at least half of targets, yielding amplicons of at least 100 bp (**Figure S3c**, **Methods**). If the representative rgRNA for a merged group failed this screen, we evaluated alternatives from the same group. Only 28.3% (human), 27.6% (mouse), and 23.7% (zebrafish) of unique rgRNAs passed this conservation filter.

For passing rgRNAs, we designed forward and reverse primers based on the most conserved flanking regions, applying quality thresholds for GC content, sequence complexity, secondary structure, dimerization probability, and melting temperature match between primers (**Methods**). Of remaining rgRNAs, 53.0% (25,346; human), 60.7% (29,005; mouse), and 96.8% (25,676; zebrafish) had at least one primer pair passing all thresholds. To select optimal primers and characterize amplification performance, we identified all genomic loci expected to be amplified by each primer pair using findMotif^65^ (**Figure S3d**, **Methods**). This analysis defined two key performance metrics. Recovery fraction measures the proportion of rgRNA targets captured by the primer pair—critical for maximizing information recovery. On-target fraction measures the proportion of all amplicons containing the rgRNA’s spacer and PAM—important for efficient use of sequencing resources, though low values are tolerable given declining sequencing costs^66–68^.

Median recovery fractions were 76.7% (human), 95.12% (mouse), and 90.9% (zebrafish), with distinct distributions across repeat families in each species (**Figure 5c, Figure S4c**). LTRs showed the highest average recovery fraction across all three species. Recovery fraction did not show a strong relationship with rgRNA target count (–0.19 < Spearman’s ρ < 0.11 for the three genomes; **Figure 5d**), indicating that even rgRNAs with many targets can be efficiently captured with a single primer pair. Median on-target fractions were 23.9% (human), 65.6% (mouse), and 50.0% (zebrafish); fractions higher than 90% were common in all three species irrespective of total target count (**Figure S4d**). For rgRNAs with multiple primer options, we selected pairs maximizing recovery fraction, then on-target fraction (**Table S4**). We excluded rgRNAs with recovery fraction <50% or on-target fraction <10%.

A final consideration is the extent to which different copies of an RGR can be distinguished after sequencing. Both our bulk and single-cell experiments with LTR5-RGR1 demonstrated that the ability to distinguish genomic targets enhances signal deconvolution similar to diversity reception in signal processing (**Figures 3d and 5i,j**). We defined a sequence distinguishability index (*SDI*) as *SDI* = (*m* – 1)/(*n* – 1), where *m* is the number of distinct target-containing amplicon sequences and *n* is the total number of covered genomic targets. *SDI* = 1 indicates that all amplified targets in the haploid genome have at least one single-nucleotide variant (SNV) or indel beyond the spacer distinguishing them from all others; *SDI* = 0 indicates that all amplified targets are identical and thus indistinguishable; values between 0 and 1 represent intermediate scenarios. Mean RGR distinguishability indices were 0.80 (human), 0.65 (mouse), and 0.73 (zebrafish), largely independent of target count and amplicon length (–0.3 < Spearman’s ρ < 0.3 for both comparisons across all genomes; **Figure S4e,f**).

These filtering and characterization steps yielded 15,201 (human), 25,606 (mouse), and 18,822 (zebrafish) rgRNA-primer pairs with 20–200 haploid targets and shared amplification sites (**Table S4**). Mean amplicon sizes were 308 bp (human; range 103–500), 307 bp (mouse; range 105–500), and 272 bp (zebrafish; range 101–500) (**Figure S4g**). We term the genomic loci targeted by an rgRNA and amplified by its primers a Repeat for Genomic Recording (RGR).

To understand how RGRs are shaped by the repetitive element landscape of each genome, we analyzed their repeat family origins (**Figure 5e**). Strikingly, RGRs with 20–200 copies are dominated by LTR elements in all three species, despite dramatically different overall repeat compositions: SINE retrotransposons are most abundant in human and mouse, whereas DNA transposons dominate in zebrafish^31,32,69,70^. Comparing repeat subclass representation in RGRs to each genome’s background distribution revealed differential enrichment (**Figure 5f**). Specifically, ERV1, ERVL, and ERVK LTRs were most enriched among human RGRs; Alu elements, which are the most abundant interspersed repeat subfamily in the human genome and were previously used for multiplexed genome perturbation with Cas9^29^, were depleted. ERVK and ERV1 LTRs were most enriched among mouse RGRs, and Gypsy, Ngaro, and ERV1 LTRs together with L1 LINEs were most enriched among zebrafish RGRs. These results reveal that the RGR landscape is unique to each genome.

Beyond enrichment, repeat subfamilies show a rich diversity of properties relevant to genomic recording in each species. For example, ERVK RGRs in both human and mouse have a higher average functionality hazard percentage than L2 LINE RGRs (**Figure 5g**). On the other hand, sequence distinguishability is, on average, higher in ERVK RGRs than L2 LINE RGRs in both species (**Figure 5h**). Other recording-relevant features, including total targets, recovery fraction, on-target fraction, and amplicon length, similarly show mosaic patterns across repeat subfamilies in each species (**Figure S5**). These results underscore the importance of systematic cataloguing to facilitate selection of RGRs tailored to each application. **Table S4** catalogues all of these relevant parameters for each identified RGR in each species.

In summary, our computational pipeline identifies a surprisingly large catalog of candidate RGRs across human, mouse, and zebrafish genomes and characterizes their properties relevant for genomic recording (i.e., copy number, distinguishability, recovery fraction, on-target fraction, functionality hazard), establishing a resource for RGR-based recording across diverse applications.

### Experimental validation of predicted RGRs

To validate our RGR mining pipeline, we tested 20 predicted human RGRs and 14 predicted mouse RGRs with high recovery fractions across the target count range. We first tested primer performance by amplifying each RGR from genomic DNA (HEK293T cells for human, N2A cells for mouse) and comparing the observed number of distinct amplicon sequences to predictions. Observed and predicted values showed excellent agreement in both species (Pearson’s *r* = 0.984 and 0.999 for human and mouse, respectively; p < 5.4×10^-15^ and p < 1×10^-17^; **Figure 6a**), demonstrating that our primer design strategy performs as expected.

**Figure 6:**
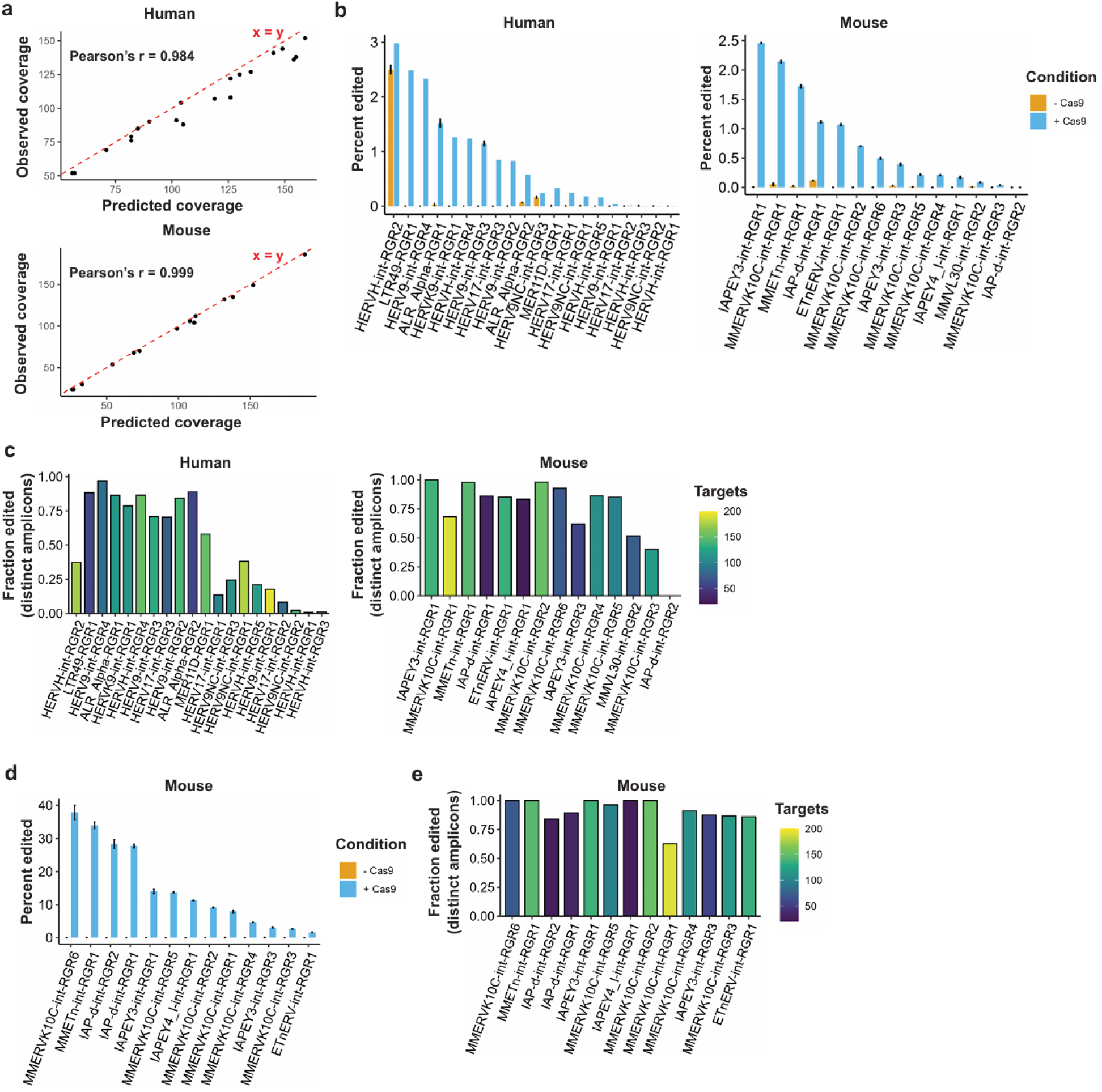
Testing performance of predicted RGRs. a, Observed versus predicted recovery of target sites using optimal primer pairs. Scatter plots compare the predicted number of targets covered by the pipeline-identified optimal primer pairs (x-axis) against empirically observed recovery (y-axis) for human (top) and mouse (bottom) rgRNAs. Each data point represents an individual rgRNA tested in vitro. Red dashed line indicates perfect agreement (x = y, labeled). Pearson’s correlation coefficient (r) is indicated for each species. b, RGR editing efficiency following transient transfection. Bar plots show the percentage of target-containing reads with successful edits 72 h post-transfection in HEK cells (human, left) and N2A cells (mouse, right). rgRNAs are sorted by descending editing. Cells were co-transfected with Cas9 and the respective rgRNA (+Cas9, blue) or transfected with rgRNA alone (-Cas9 control, orange). Bars represent mean editing efficiency; error bars, s.e.m. (n = 2 biological replicates). c, Barplots showing the fraction of distinguishable rgRNA amplicons showing significant editing. Amplicons were classified as "significantly edited" if their average editing in samples with Cas9 was strictly greater than the background noise limit, defined as the average editing in the no Cas9 samples plus one SEM (of the no Cas9 samples). rgRNAs are sorted as in b by descending overall editing, and colored by the number of sites targets. d, Enhanced editing efficiency in stably expressing mouse cell lines. Evaluation of mouse rgRNAs using an optimized experimental design to guarantee all cells expressing both Cas9 and rgRNA. The bar plot displays the overall percent editing achieved in NIH/3T3 cells engineered to stably express Cas9 and the indicated rgRNA (+Cas9, light blue) compared to negative controls (-Cas9, orange). rgRNAs are sorted by descending editing. Bars represent the mean percent editing +/- SEM for n=2 independent biological replicates. e, Bar plot indicating the fraction of distinguishable target amplicons demonstrating significant editing for the stable cell line experiments shown in d. Significance was determined using the identical background noise threshold criteria defined in c. rgRNAs are sorted as in d by descending overall editing, and colored by the number of sites targets.

We next tested CRISPR targeting by transiently transfecting HEK293T or N2A cells with vectors expressing Cas9 and one of the rgRNAs, then sequencing RGR amplicons after 72 hours. Of 20 human RGRs, 16 showed significant editing above non-transfected controls; of 14 mouse RGRs, 13 showed significant editing (**Figure 6b**). One apparent exception, HERVH-int-RGR2, showed high background signal without Cas9; this proved to be an off-target amplicon whose sequence resembled a Cas9-edited product. Total editing ranged from 0.1–2.5% of reads in both species, as expected given transient transfection without selection. The fraction of distinguishable amplicons with detectable editing varied across RGRs, averaging 49% for human and 74% for mouse (**Figure 6c**).

To circumvent the limitations of transient transfection, we used lentivirus to stably transduce mouse NIH/3T3 cells with Cas9 and each of the 13 mouse rgRNAs that showed editing in transient transfection. Selection markers (puromycin for Cas9 and hygromycin for rgRNA) enabled isolating a pure double-transduced cell population for each rgRNA. After 72 hours in culture, RGR amplicons were sequenced (**Figure 6d**). Total editing rates increased for all RGRs under stable transduction, ranging from 3% to nearly 40%. The fraction of distinguishable amplicons with detectable editing within each RGR also increased compared to transient transfection, with an average of 91% targets showing editing (**Figure 6e**). Close inspection of coverage and edit patterns within each RGR revealed a strong correlation between the predicted haploid copy number of each distinct sequence and its relative abundance in sequencing (median Pearson’s *r* = 0.838, **Figure S6a**). This observation supports the copy number predictions of the computational pipeline. Moreover, all RGRs showed a wide range of editing sensitivities across their distinct sequences (**Figure S6b**), preserving a feature of human LTR5-RGR1 critical for high-resolution and single-cell recording. These results validate our computational RGR predictions, demonstrating that identified rgRNAs effectively target editable sites that can be efficiently recovered with a single pair of amplification primers, and establish a foundation for translating these RGRs to in vivo recording applications.

### Recording long-term immediate early gene activity across brain in live mice

The identification of mouse RGRs and their validation in cultured cells paved the way for RGR-based recording in live animals. We focused on immediate early genes (IEGs), which are activated rapidly and transiently in response to cellular stimulation^71–74^. IEG-inducing stimuli vary by cell type and include electrical activity in neurons and metabolic signals in hepatocytes^75–78^. Because IEGs are activated during normal physiology, their cumulative activity should scale with elapsed time. We thus tested whether RGRs can capture this relationship.

For the recording locus, we selected MMERVK10C-int-RGR1 from the mouse RGRs validated in NIH/3T3 cells (**Figures 6b, 6c, S6**). This RGR has the largest copy number among those tested (188 copies; **Figure 7a**) with a high distinguishability index (*I_d_* = 0.67), leading to 132 recording channels. It also provides efficient editing together with high recovery (99%) and on-target fractions (65%). Critically, it has a functionality hazard score of zero—none of its targets overlaps with known functional elements.

**Figure 7:**
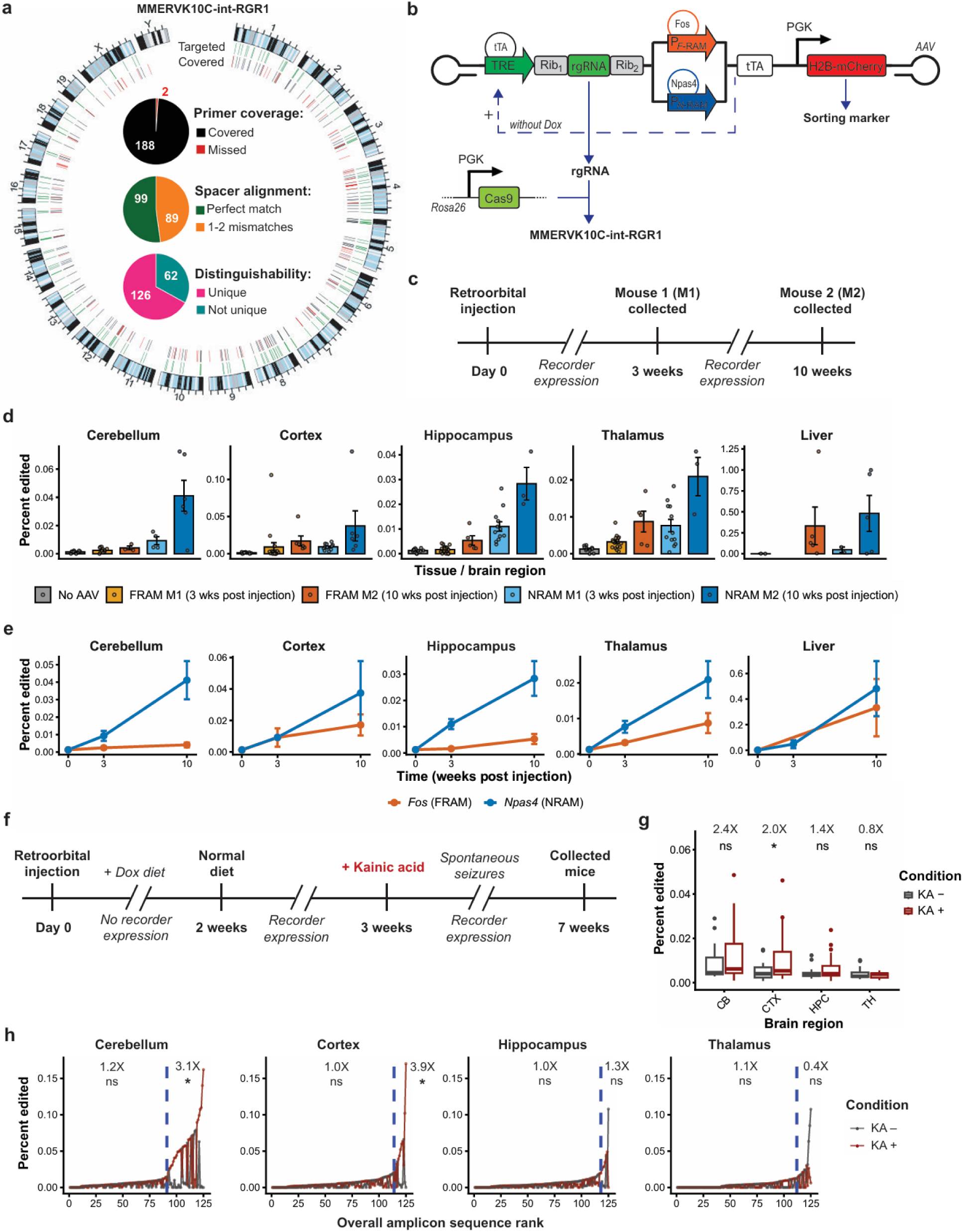
Recording immediately early gene activity in live mice. **a**, Genomic distribution and recovery of MMERVK10C-int-RGR1 target sites. First inner ring: sites targeted by MMERVK10C-int-RGR1 (green). Second inner ring: amplicon coverage, with amplicons covering a target site (black) or not covering a target site (red). Pie charts (center) indicate the proportion of target sites expected to be covered by sequencing (top), the proportion of covered target sites with perfect rgRNA sequence match (middle), and the proportion containing at least one SNP distinguishing them from all other target-containing amplicons (bottom). **b**, Schematic of FRAM and NRAM rgRNA recorder AAV vectors. P_FRAM_: Fos promoter (4X response elements); P_NRAM_: Npas4 promoter (4X response elements); Rib_1_ = hammerhead ribozyme; Rib_2_ = HDV ribozyme. In the absence of doxycycline, Npas4 or Fos activity is linked to expression of MMERVK10C-int-RGR1 rgRNA. **c**, Experimental timeline for in vivo recording. Mice received retroorbital AAV injection on day 0. Tissues were collected at 3 weeks (M1) and 10 weeks (M2) post-injection. **d**, In vivo editing efficiency of FRAM (orange) and NRAM (blue) recording systems across four brain regions (CB: cerebellum; CTX: cortex; HPC; hippocampus, TH; thalamus) and liver at 3 weeks (M1) and 10 weeks (M2) post-injection. "No AAV" (gray) represents an uninjected Cas9 mouse. Individual dots represent 100 sorted cells (NeuN^+^ neuronal nuclei for brain; mCherry^+^ for FRAM and NRAM samples. Bars represent mean ± s.e.m. **e**, Time-course analysis of edit accumulation from 0 to 10 weeks post-injection for FRAM (orange) and NRAM (blue) systems across tissues. The 0-week baseline represents pooled "No AAV" control data (shared across brain regions; separate for liver). Data points represent mean ± s.e.m. Pearson correlation coefficients (R) indicate the strength of the linear relationship between time and editing efficiency. Note: y-axes are scales independently for each tissue to accommodate varying editing ranges. **f**, Experimental timeline for in vivo Fos recording of kainic acid (KA) induced seizures. Mice received retroorbital AAV injection on day 0 and placed on a doxycycline diet for 2 weeks. Mice were taken off of doxycycline 1 week before kainic acid injections. Injected mice experienced recurring seizures for four weeks before mice were collected at 7 weeks post AAV injection (3 KA + mice, and 2 KA - mice). **g**, In vivo editing efficiency of Fos recording systems across four brain regions (CB: cerebellum; CTX: cortex; HPC; hippocampus, TH; thalamus) in kainic acid induced seizure model. Box plots display MMERVK10C-int-RGR1 editing, and individual data points represent 50 sorted cells (NeuN^+^ neuronal nuclei; mCherry^+^ for FRAM samples), for mice that received kainic acid induced seizures (dark red) vs mice that did not (dark gray). \**P* < 0.05, \*\**P* < 0.01, \*\*\**P* < 0.001, \*\*\*\**P* < 0.0001 (one-sided Wilcoxon rank-sum test); n.s., not significant. Average fold increases in KA-treated samples is labeled above significance. **h**, Rank plots of editing per MMERVK10C-int-RGR1 amplicon across regions in mice that received kainic acid induced seizures (dark red) vs mice that did not (dark gray). Amplicons ranked per region by maximum of the average editing across KA + and the average editing across KA - mice. Dashed blue vertical lines denote threshold for sites with editing rates above background. \**P* < 0.05, \*\**P* < 0.01, \*\*\**P* < 0.001, \*\*\*\**P* < 0.0001 (paired Wilcoxon signed-rank test); n.s., not significant. Average fold increases in KA-treated samples is labeled above significance.

We designed AAV vectors that express the MMERVK10C-int-RGR1 rgRNA in response to IEG activity, driven by either the *Fos* or *Npas4* promoter (**Figure 7b**)^79^. The rgRNA is expressed under a tetracycline-responsive element (TRE) and flanked by cis-cleaving ribozymes to ensure nuclear localization^80,81^. The tetracycline-controlled transactivator (tTA) is expressed under the IEG-responsive promoter; in the absence of doxycycline, tTA binds TRE and drives rgRNA expression, coupling IEG activation to rgRNA production. The vector also constitutively expresses H2B-mCherry to identify transduced cells (**Figure 7b**).We tested this system in transgenic mice constitutively expressing Cas9^82^. We injected four mice with 1–5×10^10^ viral genomes of the IEG-recording AAV via the retroorbital route (two mice each for *Npas4* and *Fos* recorders). We used the AAV-PHP.eB serotype, which traverses the blood-brain barrier, to achieve systemic transduction across the brain and different organs^83^. For each IEG construct, one mouse was euthanized 3 weeks after injection and the other at 10 weeks (**Figure 7c**). Mice never received doxycycline, ensuring recording remained active throughout. At each timepoint, we sorted 300–2000 mCherry-positive single nuclei from representative sites: liver, cortex, hippocampus, thalamus, and cerebellum (**Figure S7**). For the brain-derived samples, we used NeuN immunostaining to isolate neuronal nuclei. As a control, we sorted equivalent numbers of neuronal nuclei from untransduced mice. We amplified MMERVK10C-int-RGR1 using its corresponding primers (**Table S4**) and sequenced amplicons from each nucleus. Editing was detectable above untransduced controls in all tissue samples, confirming successful RGR targeting in vivo (**Figure 7d**).

Editing rates increased with time, correlating positively with elapsed time across all tissues and both IEGs (median Pearson’s *r* = 0.569, range 0.27–0.84; **Figure 7e**). Editing levels were higher in the liver compared to brain tissues, likely reflecting lower Cas9 activity in neurons in this mouse model^84,85^. These results show that MMERVK10C-int-RGR1 can record the cumulative IEG signal over weeks.

We next asked whether RGR-based recording could detect altered IEG activity in a disease model. We used kainic acid (KA)-induced *status epilepticus* (KASE), a well-established model of seizures and epilepsy. The temporal dynamics of KASE are well characterized over short and medium timescales. In the short term (hours), systemic KA injection induces *status epilepticus* (prolonged seizures) and IEG *c-Fos* mRNA peaks within 4 hours across the brain, with highest intensity in cortex and hippocampus^86,87^—thalamus and cerebellum show much weaker responses^86,88,89^ (e.g., ≥25-fold lower than hippocampus in the hours after KA injection^89^ **Methods**). In the medium term (days), the animal enters the epileptogenesis phase (latent epilepsy) during which the brain undergoes extensive neuronal degeneration in the thalamus, parts of the cortex, and most dramatically in the hippocampus^90^. IEG returns to the detection baseline during this period^86^. In the long term (weeks), chronic temporal lobe epilepsy develops, leading to spontaneous recurrent seizures. Long-term IEG activity during this process remains poorly characterized, as conventional methods (e.g., in situ hybridization) are better suited to capturing acute responses than activity that is either intermittent or sustained at lower levels over extended periods.

To investigate these long-term changes, we injected five Cas9 mice with the Fos IEG-recording AAV. Three weeks later, we administered KA to three mice to induce *status epilepticus*, with two receiving saline as controls (**Figure 7f, Methods**). Four weeks after KA injection, we harvested brains and sorted NeuN-positive, mCherry-positive neuronal nuclei from cerebellum, cortex, hippocampus, and thalamus. We collected 3–10 replicates of 50 nuclei per sample, amplified MMERVK10C-int-RGR1, and sequenced the amplicons. KA-treated mice showed 2.4-, 2.0-, 1.4-, and 0.8-fold increases in editing (cerebellum, cortex, hippocampus, and thalamus, respectively), with cortex reaching statistical significance (one-sided Wilcoxon rank-sum test, *P* < 0.05; **Figure 7g**). These fold changes represent cumulative IEG activity over the four weeks following KASE induction.

Deconvolving editing rates to individual sites provided additional resolution (**Figure 7h**). The subset of sites with editing rates above background in each region (**Methods**) showed 3.1-, 3.9-, 1.3-, and 0.4-fold increases in KA-treated compared to untreated samples for cerebellum, cortex, hippocampus, and thalamus, respectively. The increases in cerebellum and cortex were significant (paired Wilcoxon signed-rank test, *P* < 0.05). Overall, these results reveal stronger cumulative IEG activity in surviving neurons of cortex and cerebellum than in hippocampus and thalamus. Because the recorder is active in all four regions (**Figure 7e**)—and KASE induces acute IEG responses in all four—the divergent long-term signals likely reflect regional differences in neuronal survival. The low cumulative signal in the hippocampus, despite strong acute IEG induction^89^, is consistent with preferential loss of KASE-activated neurons during epileptogenesis. The low signal in the thalamus likely reflects both its weak acute response and subsequent neuronal degeneration^86,89,90^. The higher signal in the cerebellum, despite relatively mild acute IEG induction, is consistent with long-term survival of cerebellar granule neurons, which constitute more than 99% of NeuN+ neurons in this region^91^. These findings demonstrate that RGR-based recording in live mice can integrate neuronal activity across the brain over weeks, capturing cumulative IEG signals that reflect both region-specific activity levels and survival outcomes—dynamics that are inaccessible to snapshot methods. Compared to conventional longitudinal methods, such as GCaMP or electrode array recordings, RGR-based recording is not restricted to a limited number of regions and allows for simultaneous recording in the entire brain. Altogether, this method provides a flexible framework for longitudinal studies of neuronal activity across the whole brain.

## Discussion

In this work, we establish genomic repeats as a platform for expanding the capacity and resolution of molecular recording. By targeting hundreds of similar sequences with a single guide RNA and reading them out with a common primer pair, RGRs overcome a fundamental bottleneck in genomic recording: the scarcity of accessible writing space. Using LTR5-RGR1 as a prototype, we demonstrate that this approach enables accurate discrimination of multiple signal levels in single cells—a capability that has remained elusive with conventional single- or double-site approaches. We further develop a computational pipeline to systematically identify RGRs across human, mouse, and zebrafish genomes, revealing thousands of candidates with favorable properties and validating predictions experimentally. Finally, we leverage RGRs in live mice to record immediate early gene activity across tissues and in a disease model, demonstrating in vivo applicability.

The advantage of RGRs derives not from redundancy alone but from sequence diversity. Different RGR copies exhibit heterogeneous editing kinetics: some respond sensitively to low signals and saturate early, while others accumulate edits more slowly. This kinetic diversity enables RGRs to capture differences across a wide temporal and concentration range that would be inaccessible to conventional approaches. This principle parallels diversity reception, a classic concept in signal processing in which combining channels with distinct response characteristics—rather than duplicating identical ones—enables robust signal detection across varying conditions^92^. By analogy, we term this capability diversity recording: multi-channel signal capture through genomic loci with heterogeneous response kinetics. Diversity recording provides a boost to signal deconvolution capacity without added engineering complexity. Our single-cell classification analysis quantifies this advantage, showing that combining sites with diverse kinetics achieves 2-fold higher signal discrimination than single- or double-copy sites alone. Machine learning approaches that leverage amplicon-specific patterns further improve accuracy, demonstrating how the high-dimensional information encoded across RGR copies can be computationally exploited.

Repetitive elements constitute over half of mammalian genomes, yet our systematic analysis reveals that only a small fraction possess the properties required for effective recording: a shared CRISPR target, avoidance of functional elements, and conserved primer binding sites. Depending on the genome, only 0.7–1.8% of distinct Cas9 targets in interspersed repeats qualified as RGRs—rare needles in a genomic haystack. The computational pipeline we developed navigates this complexity, filtering millions of candidate gRNAs through successive criteria to identify the thousands that meet all requirements. Yet despite these stringent requirements, the total number of RGRs identified is surprisingly large: 15,000 to 25,000 per genome, providing a wealth of options for diverse recording applications. The curated catalogs we provide distill this vast repetitive landscape into an optimized resource, with metrics characterizing each RGR’s properties relevant to genomic recording facilitating selection of RGRs tailored to specific applications.

Several considerations will guide the selection and optimization of RGRs for particular use cases. Target number is a key parameter: fewer targets reduce potential off-target effects and cellular burden, while more targets increase recording capacity and robustness. Our resource offers flexibility across this range, with RGRs spanning 20 to 200 targets in the catalogs provided. The functionality hazard score provides a starting point for identifying RGRs that avoid interference with coding and regulatory elements, but context-dependent effects likely necessitate empirical validation in target cell types. Integration of RGR target positions with cell type-specific epigenomic signatures (e.g., ATAC-seq accessibility or histone modification profiles) could refine functionality hazard estimates given cell type context. Moreover, the diversity of RGRs enables functional screening to identify candidates that do not interfere with biological processes of interest. Additional metrics such as the distinguishability index and recovery fraction can further guide selection based on readout requirements. Finally, the choice of writing system (i.e., Cas9 nuclease, base editor, prime editor, or compound architectures) will depend on application-specific efficiency and performance requirements. Our in vivo experiments provide initial evidence that pipeline-identified RGRs can function across diverse biological contexts: a single mouse RGR recorded IEG activity across the liver and brain over 10 weeks; it also recorded differential cumulative IEG signal across brain regions in an epilepsy model. These results suggest that appropriately selected RGRs can support long-term recording of endogenous signals in living animals, addressing a long-standing challenge in genomic recording.

Looking forward, RGR-based recording opens several avenues for future development. The large number of identified RGRs provides opportunities for multiplexed recording of multiple signals simultaneously in single cells, with different RGRs dedicated to different biological inputs. Higher-capacity writing systems, such as peCHYRON and DNA Typewriter, could be adapted for RGRs to further expand per-site information storage. Our pipeline can also be readily extended to incorporate updated genome annotations, other Cas proteins (e.g., Cas9 from other bacteria, Cas12, and nucleases yet to be developed), and other species, potentially expanding the repertoire of targetable repeats and editing modalities. Integration with single-cell sequencing technologies could enable lineage-coupled recording of cellular states during development or disease progression.

In conclusion, this work establishes genomic repeats as a readily accessible, high-capacity platform for molecular recording. By leveraging the natural redundancy and diversity of repetitive elements, RGRs enable single-cell resolution recording with minimal engineering complexity. The computational pipeline and curated catalogs we provide distill the vast repetitive landscape into a practical resource, establishing a foundation for deploying this approach across species and applications.Materials and Methods

### Animal procedures

All animal procedures were approved by the Animal Care and Use Committee (ACUC) of Johns Hopkins University. All experiments were performed in accordance with relevant institutional and national guidelines and regulations.

### Cloning Plasmids

Standard molecular cloning techniques were used to assemble constructs in this paper, including standard NEB HiFi (Gibson) assembly, Golden Gate Assembly, and restriction–ligation cloning. Oligos for PCR amplification, spacers, and 3’ extension sequences (all oligos used for cloning) were ordered from IDT (Integrated DNA Technologies). All primers and oligos described below were ordered from IDT and are shown in 5’ to 3’ orientation (**Table S5**). All transformations were done in NEB Stbl cell (C3040) at 37°C unless otherwise noted. All resulting plasmids were prepared with Qiagen Mini Prep or Zymo Midi/Maxi Prep kits, and were sequence verified using PlasmidExpress (Quintara Bio), Plasmidsaurus, or Sanger sequencing (Genewiz). All plasmids are available along with full sequences at Addgene (*link to deposited plasmids on Addgene*).

To make the lentiviral construct expressing the LTR5-RGR1-gRNA, sgRNA oligos (sequences in Table S5) were annealed and cloned into LentiGuide-hygro^93^. LTR5 pegRNA constructs were cloned using Golden Gate Assembly into the backbone plasmid pU6-pegRNA-GG-acceptor (Addgene plasmid 132777), following the protocol outlined in Doman et al^94^. Single-stranded oligos for scaffold, spacer, and 3’ extension sequences (Integrated DNA Technologies) were annealed with 4 bp overhangs, and the scaffold phosphorylated (PNK, NEB). Cloning backbones were digested with either BsaI-HFv2 or BsmBI-v2 (NEB). Ligation of digested backbone and annealed oligos with T4 DNA ligase (NEB), and 2 uL of ligation products were added to an NEB Stbl cell (C3040) for transformation with cells grown at 30 °C for plasmid DNA preparation (Qiagen Miniprep). For integration of pegRNA constructs, puromycin resistance was replaced with hygromycin in the backbone plasmid pPBT-peRNA_GG-Puro (Addgene plasmid 173220), then used the same Golden Gate Assembly to clone pegRNAs as described above.

Inducible PE2 constructs were cloned by a combination of NEB HiFi (Gibson) Assembly and ligations. Fragments were either pcr amplified or enzyme digested from existing plasmids then gel extracted. 4HT-PE2 was cloned by replacing the Cas9 in Addgene plasmid 84232 (expression of Cas9 fused to 4X ERT2 proteins, for nuclear localization control by 4-hydroxytamoxifen)^95^, with PE2 (from pCMV-PE2, Addgene plasmid 132775), by NEB HiFi Assembly. 4HT-PE2 insert was digested from plasmid with BstBI and NotI (11 kb), then ligated with annealed oligos to add homology arms for assembly, and backbone was digested with EcoR1 and Acc65I (9 kb). TRE-4HT-PE2 was cloned by putting the 4HT-PE2 insert into a Piggybac doxycycline inducible backbone by assembly of the backbone and oligo annealed insert fragments. TRE-PE2 was cloned in a similar manner, by putting the PE2 from pCMV-PE2 by itself into a Piggybac doxycycline inducible backbone. In both cases, replacing Cas9 with either the 4HT-PE2 insert or PE2 insert.

DD-PE2-GR constructs were cloned by NEB HiFi (Gibson) Assembly. DD-PE2-GR was cloned by replacing the IsceI in 150911_pLEX_DD-IsceI-GR (source: Reza Kalhor), by PCR amplification of the entire backbone outside of IsceI, with PE2 (again pcr amplified from pCMV-PE2). TRE-DD-PE2-GR was cloned by replacing the IsceI in 150911_pLIX403_DD-IsceI-GR (source: Reza Kalhor), by double digestion of the backbone with NheI and SalI (8.5 kb fragment containing entire backbone except DD and IsceI), pcr amplification of the destabilizing domain from the same plasmid, with PE2 (again pcr amplified from pCMV-PE2).

The N-terminal and C-terminal FKBP destabilizing domain fused PE2 plasmids were cloned by NEB HiFi (Gibson) Assembly to replace the YFP in Addgene plasmids 31763 and 31766 with PE2 (again pcr amplified from pCMV-PE2). Backbones were digested with EcoRI and XhoI (N-terminal FKBP-DD) or BamHI (C-terminal FKBP-DD), and the ∼7.3 kb fragment was gel extracted to use as a backbone. To clone the dox inducible version of both plasmids, the FKBP-DD fused PE2 was PCR amplified out of the FKBP-DD-PE2 and PE2-FKBP-DD constructs, and inserted into a Piggybac doxycycline inducible backbone.

NFκB-PE2 was cloned by replacing Cas9 in NFκBRp_Cas9_3xNLS_p2a-puroR (Addgene plasmid #81255) with PE2 by NEB HiFi (Gibson) Assembly. To get the backbone, NFκBRp_Cas9_3xNLS_p2a-puroR was digested with AgeI and BamH1, and the 8.9kb fragment was gel extracted and purified. The PE2 insert was again pcr amplified from pCMV-PE2, adding homology arms overlapping the backbone.

Mouse and human selected RGRs were cloned using Golden Gate Assembly into the pU6-tevopreq1-GG-acceptor backbone (Addgene plasmid #174038) or a hygromycin resistant epegRNA backbone. The hygromycin resistant epegRNA backbone plasmid lenti-tmpknot-epegRNA-hyg was cloned by replacing the puromycin resistance in lenti-tmpknot-epegRNA-puro (Addgene plasmid #214089) with hygromycin, as was done with the pU6-pegRNA-GG-acceptor plasmid. Single-stranded oligos for scaffold, spacer, and 3’ extension sequences (Integrated DNA Technologies, **Table S5**) were annealed with 4 bp overhangs, and the scaffold phosphorylated (PNK, NEB). Cloning backbones were digested with either BsaI-HFv2 or BsmBI-v2 (NEB). Ligation of digested backbone and annealed oligos with T4 DNA ligase (NEB), and 2 uL of ligation products were added to an NEB Stbl cell (C3040) for transformation with cells grown at 30 °C for plasmid DNA preparation (Qiagen Miniprep).

Mouse RGRs were cloned into the lentiCRISPR v2 backbone (Addgene plasmid #52961)^96^ via Golden Gate Assembly to insert the rgRNA spacers at the BsmBI cut site. Spacer oligos used for the cloning are listed in **Table S5**.

FRAM and NRAM rgRNA AAV constructs were cloned from pAAV-F-RAM-d2tTA-TRE-mKate2 (Addgene plasmid #140274) and pAAV-N-RAM-d2tTA-TRE-mKate2 (Addgene plasmid #140275) by NEB HiFi (Gibson) Assembly of five fragments. The overall cloning scheme was to replace mKate2 with ribozyme flanked rgRNA, and to add a constitutively expressed nuclear mCherry outside of the dox controlled elements. The AAV backbone and F/NRAM enhancer modules were PCR amplified from the parent plasmids, adding homology arms for Gibson assembly. Ribozyme flanked rgRNAs were ordered as gblocks from IDT. The PGK promoter placed in front of mCherry was PCR amplified from another in-house partially complete plasmid, and the H2B-mCherry was PCR amplified from R26-H2B-mCherry HR donor vector (Addgene plasmid #137928), adding homology arms to both.

### Cell culture (General mammalian cell culture conditions)

HEK cells were purchased from ATCC and cultured in Dulbecco’s modified Eagle’s medium (DMEM), supplemented with 10% fetal bovine serum (Gibco, qualified, tet-free) and 1× penicillin streptomycin (Corning). Cells were incubated, maintained, and cultured at 37 °C with 5% CO2. Cell lines were authenticated by their respective suppliers and tested negative for mycoplasma. All cells treated with doxycycline received 500 ng/uL doxycycline unless otherwise specified. In time course experiments media was exchanged daily with freshly prepared treatment (dox and/or Shield1) or non-treatment containing media. Puromycin selection media was prepared by diluting 10 µL of 10 mg/mL stock per 50 mL media, to get 2 µg/mL puromycin media. Hygromycin media was prepared by diluting 100 µL of 50 mg/mL stock per 50 mL media, to get 100 µg/mL hygromycin media).

### HEK tissue culture transfection protocol

HEK cells were seeded on plates (Corning) coated with poly-D-Lysine. Between 16 and 24 hours after seeding, cells were transfected at approximately 60% confluency with lipofectamine 2000 (Thermo Fisher Scientific) according to the manufacturer’s protocols. Unless otherwise stated, media was refreshed 18-24 hours post transfection, and when applicable selection media was added 48 hours post transfection. Cells were cultured for 2-5 days following transfection, after which the medium was removed, the cells were washed with 1X phosphate-buffered saline (PBS) solution (Thermo Fisher Scientific), and genomic DNA was extracted following the bulk/population level protocol below.

### Lentivirus production

Lentiviruses were packaged in HEK cells using a second generation system with VSV.G as the envelope protein. Lentiviral supernatant was filtered with 0.45 micron filters, and viral particles were purified and concentrated using polyethylene glycol precipitation and resuspension in PBS. Purified and concentrated lentivirus was stored at −70 °C until use.

### Lentiviral transduction protocol

Cells were seeded on PDL-coated plates 24 hours prior to transduction at 30-40% confluency, to be at 60-70% confluency 24 hours later. Cell media was refreshed 30-60 minutes before transduction. Cells were infected with lentiviral particles packaged as described above, in the presence of 6 μg/ml polybrene. Polybrene and virus containing media was freshed 18-20 hours post infection. 48 hours post infection, cells were placed under puromycin (2 µg/mL) or hygromycin (100 µg/mL) selection and passaged and maintained under selection at least one passage past when non infected cells treated with the same level of selection had all died to assure genomic integration before use in any experiments. Cells were maintained under respective selection throughout all experiments.

### Cas9 editing of LTR5-RGR1

Previously established clonal HEK cell line with Dox inducible S. Pyogenes Cas9^22^ were transduced with lentivirus expressing the LTR5-RGR1-gRNA. Hygromycin selection was started 48 hours post transduction as described above. After selection, cells were seeded in a 48-well plate and treated with 0 to 50 ng/mL of doxycycline. Media with respective doxycycline concentration was prepared and refreshed daily. To get the appropriate doxycycline concentrations, 1 mg/mL stock solution of doxycycline in DMSO was first diluted 1:500 in media. Then, the 2 µg/mL doxycycline media was diluted 1:10 to 200 ng/mL. 1 to 50 ng/ml doxycycline media was then prepared by further diluting the 200 ng/mL doxycycline media to the respective concentration. Serial dilutions were performed so the volume pipetted never dropped below 5 uL. Cells were collected and the LTR5-RGR1 locus amplified and sequenced after 4 days of treatment with Illumina with at least 10,000 reads per sample.

### Testing LTR5-RGR1 pegRNAs

HEK cells were seeded at ∼30-40% confluency on PDL-coated 24 well plates. 24 hours later, 60-80% confluent cells were co-transfected with a constitutively expressed prime editor (pCMV-PE2, Addgene plasmid 132775) and either an LTR5-RGR1 pegRNA or no pegRNA. Each well was transfected with 750 ng PE2 and 250 ng pegRNA (3:1 PE2:pegRNA ratio). Cells were passaged 18 hours later, and collected 38 hours after passaging (56 hours post transfection). LTR5-RGR1 amplicons were amplified and sequenced with Illumina with at least 5000 reads per sample. Amplicons were called as seen if >= 10 reads aligned to that amplicon.

### Establishing HEK cell line stably expressing LTR5-RGR1-pegRNA#1

HEK cells were co-transfected with transposase and LTR5 peg-rgRNA (1:3 ratio) for PiggyBac mediated transposition of the LTR5 peg-rgRNA (hygromycin resistance) plasmid. Hygromycin (100 µg/ml) selection began 48 hours after transfection. Cells were passaged three times over a period of 8 days, with selection media being replaced every 24-48 hours, until a negative control well of un-transfected HEK cells was dead from receiving the same hygromycin selection.

### Testing inducible prime editor constructs

To test inducible prime editor constructs, HEK cells stably expressing LTR5-RGR1-pegRNA#1 were transiently transfected in a 6 well plate with 2 µg of one of the inducible constructs and induced as described below. Puromycin selection and induction began 48 hours after transfection. Selection and induction media was prepared and refreshed daily. Cells were collected after 3 days of max induction (5 days post transfection), and the LTR5-RGR1 amplicon sequenced with ≥5,000 reads per sample.

Induction media for TRE-PE2 contained 500 ng/mL doxycycline. 500 ng/mL doxycycline media was prepared as 1 µL of 1 mg/mL stock solution in DMSO per 2 mL media. Induction media for 4HT-PE2 contained 1 µM 4-hydroxytamoxifen (4HT), and for TRE-4HT-PE2 contained both 500 ng/mL doxycycline and 1 µM 4HT. 1 mM 4HT solution was prepared from 10 mg powder (H6278-10MG), and 1 µL of this 1mM stock was diluted in 1 mL media to get 1 µM 4HT media for induction. Induction media for Deg-PE2 and PE2-Deg contained 1 µM Shield1. 1 µM Shield1 media was prepared by adding 2 µL of 0.5mM Shield1 stock solution (Takara Bio 632189) per 1 mL media. Induction media for Deg-PE2-GR contained 1 µM Shield1 and 0.1 µM triamcinolone acetonide (TA). 20 mg/mL TA in DMSOR was prepared from 1g powder (PHR1701-1G), and 2.2 µL of this 20 mg/mL stock was diluted in 1 mL media to get 0.1 µM TA media for induction. 0.1 µM TA has been shown to be sufficient to achieve rapid and complete translocation from the cytoplasm to the nucleus within 30 to 60 minutes^97^, since it has a very high affinity for the glucocorticoid receptor (GR)^98,99^.

To test TRE-Deg-PE2 and TRE-PE2-Deg, transfection and selection were performed as described above. 24 hours post transfection, cells were passaged to a 24 well plate. When puromycin selection began 48 hours post transfection, cells were treated with one of six Shield1 concentrations (0, 0.01, 0.05, 0.1, 0.5, and 1 µM Shield1) in the presence or absence of 500 ng/mL doxycycline. As described above, selection and induction media was prepared and refreshed daily. Cells were collected after 3 days of max induction (5 days post transfection), and the LTR5-RGR1 amplicon sequenced with ≥5,000 reads per sample.

### Obtaining monoclonal HEK cell lines with LTR5-RGR1-pegRNA#1 and double-inducible prime-editor

HEK cells were co-transfected with transposase, TRE-Deg-PE2, and LTR5-RGR1 pegRNA #1 (4:12:3 ratio: transposase 4, TRE-Deg-PE2 12, LTR5-RGR1 pegRNA #1 3) for PiggyBac mediated transposition of the TRE-Deg-PE2 (Puromycin resistance) and LTR5-RGR1 pegRNA #1 (hygromycin resistance) plasmids. Puromycin (2 µg/ml) and hygromycin (100 µg/ml) selection began 48 hours after transfection. Cells were passaged twice before FACS sorting single cells into a 96 well plate with 150 uL selection containing media per well. Selection media was exchanged every 3-4 days after sorting for ∼ 2 weeks, until wells with surviving cells were clearly visible. These wells with monoclonal populations were passaged and expanded to larger wells until there were ∼1E6 cells per cell line. Half of the cells were frozen down (5% DMSO in media, -140°C) and the other half were seeded for a short signal induction time course experiment to identify monoclonal lines with optimal induction characteristics as described in the text.

Of the 18 out of 96 wells with surviving cells expanded first to a 24 well plate, only 14 wells contained growing cell populations when media was refreshed 2 days after passaging. 2 days later, the 14 wells containing growing cell populations were passaged to larger plates; the 7 lower confluency wells were passaged to a 12 well plate for further expansion, and the remaining 7 that were fully confluent were each split between one well of a 12 well plate to continue expanding to freeze down, and 7 wells of a 48 well plate to begin testing varying levels of induction to measure dose responsiveness and background noise. These 7 monoclonal cell lines (B4, B8, C7, D3, D7, and D9) were exposed to a range of low Shield1 concentrations (0 μM, 0.005 μM, 0.01 μM, 0.02 μM, and 0.05 μM) with Dox for four days, with one well that received no treatment (no doxycycline or Shield1) to measure background editing. After four days, cells were collected and the LTR5-RGR1 amplicon was sequenced with ≥10,000 reads per sample.

### LTR5 RGR bulk/population level library preparation and sequencing

Genomic DNA was extracted by the addition of freshly prepared lysis buffer (10 mM Tris-HCl, pH 7.5; 0.05% SDS; 25 µg/ml proteinase K (ThermoFisher Scientific)) directly into each well of the tissue culture plate. The genomic DNA mixture was incubated at 37 °C for 60 minutes, followed by an 80 °C enzyme inactivation step for 20 minutes. Extracted DNA was diluted 10-100X in ultra-pure-water. 2 uL of diluted DNA was used as template for PCR1 reaction, with a PCR1 master mix containing 1X Forget-Me-Not PCR Master Mix (Biotium) and 0.5 uM each of LTR5 RGR locus amplifying forward and reverse primers, with SBS3 (forward primer) and SBS9 (reverse primer) sequence adaptors. Reactions were denatured at 95°C for three minutes, and then cycled at 95°C for 20 seconds, 60°C for 25 seconds, and 72°C for 12 seconds. PCR reactions were stopped when the real-time PCR curve reached early-to-mid-exponential phase. Reactions were then diluted 10-50X with ultra-pure-water. 2 ul of diluted PCR product was used as template for a subsequent PCR reaction in another 1X Forget-Me-Not PCR Master Mix (Biotium) and 0.25 uM each dual indexing primer pair for Illumina (P5 and P7). Reactions were denatured at 95°C for three minutes, and then cycled at 95°C for 15 seconds, 64°C for 20 seconds, and 72°C for 20 seconds. PCR reactions were stopped when the real-time PCR curve reached early-to-mid-exponential phase. Libraries were purified with DNA Clean & Concentrator-5 columns (Zymo), sequenced on an Illumina MiSeq or NovaSeq X Plus instrument, and analyzed.

### LTR5 RGR single cell library preparation and sequencing

Single cells from the iiPE2 LTR5 peg-rgRNA HEK cell line with varying levels of induced signal over time were dissociated using 0.05% Trypsin-EDTA (Life Technologies), resuspended in PBS + 10% FBS, then flow sorted on a MA900 Sonysorter for single cells. Briefly, debris was removed using FSC-A vs BSC-A to remove large events and events with high complexity. Then singlets were selected based on FSC-A vs FSC-H and FSC-A vs FSC-W.

Single cells were sorted into a 96-well plate with each well containing 1.5 ul of QuickExtract DNA Extraction Buffer (Lucigen). Plates were spun down immediately after sorting to ensure cells were in the QuickExtract. DNA was extracted from single cells by incubating the 96-well plates for 10 minutes at 65°C followed by 5 minutes at 98°C to inactivate the QuickExtract. 9.5 uL of PCR1 master mix containing 1X Forget-Me-Not PCR Master Mix (Biotium) and 0.5 uM each of LTR5 RGR locus amplifying forward and reverse primers, with SBS3 (forward primer) and SBS9 (reverse primer) sequence adaptors. Reactions were denatured at 95°C for three minutes, and then cycled at 95°C for 20 seconds, 60°C for 30 seconds, and 72°C for 12 seconds. PCR reactions were stopped when the real-time PCR curve reached early-to-mid-exponential phase. Reactions were then diluted 10-50X with ultra-pure-water. 2 ul of diluted PCR product was used as template for a subsequent PCR reaction in another 1X Forget-Me-Not PCR Master Mix (Biotium) and 0.25 uM each dual indexing primer pair for Illumina (P5 and P7). Reactions were denatured at 95°C for three minutes, and then cycled at 95°C for 15 seconds, 64°C for 20 seconds, and 72°C for 20 seconds. PCR reactions were stopped when the real-time PCR curve reached early-to-mid-exponential phase. Libraries were purified with DNA Clean & Concentrator-5 columns (Zymo), sequenced on a NovaSeq X Plus instrument, and cells with at least 50,000 reads were analyzed.

### Quantifying recording resolution of signal reconstruction - statistical analysis

To quantify the resolving power and practical significance of separation between adjacent treatment levels, a time-point-specific effect size analysis was performed. For each dataset, the data was first subsetted by collection day. Within each day, pairwise comparisons were made exclusively between neighboring treatment groups (e.g., yDx-0XShd vs. yDx-005XShd, yDx-005XShd vs. yDx-01XShd, etc.). For each of these neighboring pairs, the standardized effect size was calculated using Cohen’s *d*. Cohen’s *d* is a dimensionless metric that quantifies the magnitude of the difference between two groups relative to their pooled standard deviation. Unlike *p*-values, which indicate whether a difference exists, Cohen’s *d* provides a measure of how large that difference is in units of standard deviation, allowing for comparisons across different experiments or measurements. The statistic is calculated as:

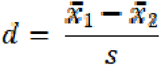

Where *x_1_* and *x_2_* are the group means and *s* is the standard deviation. The analysis assumed unequal variance and utilized the Welch’s *t*-test denominator for stabilization, calculated via the effectsize package in R. The absolute values of these Cohen’s *d* statistics were then averaged to generate a single "Average Pairwise Effect Size" score for that specific time point. This composite score serves as a metric for the dataset’s overall resolution, where a higher score indicates a greater magnitude of separation between consecutive treatment doses.

### NFκB recorder cell line responsiveness to TNF-α in vitro

To establish the NFκB recorder cell line, NFκB-PE2 lentivirus was produced as described above. HEK cells stably expressing LTR5-RGR1 pegRNA #1 were transduced with NFκB-PE2 lentivirus. Cells were treated with puromycin to select for cells expressing the NFκB-PE2 construct 48 hours post transduction, and passaged with puromycin selection 24 hours later. After three days of puromycin selection, all cells in an untransduced control well treated with the same puromycin selection were dead. The NFκB recorder cell line was maintained under puromycin and hygromycin selection throughout culture.

The NFκB recorder cell line’s responsiveness to NFκB signalling was evaluated by exposing it to various concentrations of Tumor Necrosis Factor-Alpha (TNF-α) in vitro. NFκB recorder cells were seeded in a 48 well plate and treated with one of six concentrations of TNF-α (0, 0.1, 0.5, 1, 5, and 10 ng/mL) for 4 days. Cells were collected after 2 or 4 days of each level of induction, and the LTR5-RGR1 amplicon sequenced with ≥10,000 reads per sample. Prime editing (expressed as the percentage of edited CRISPR target containing reads) was analyzed across two time points (Days 2 and 4) for each of the six TNF-α concentrations.

To determine the effect of TNF-α concentration on prime editing across different time points, a general linear model was fitted to the editing quantification data. The model included an interaction term between Day and Concentration to account for potential time-dependent variations in the dose-response relationship. To robustly estimate background variance, data from all biological replicates across all evaluated time points and TNF-α concentrations were pooled into a single model. The model evaluated the percent prime edited as the response variable, with time point (Day) and TNF-α concentration treated as interacting categorical fixed effects. Following the linear model fit, estimated marginal means (EMMs) were computed for each concentration within each time point^100^. To evaluate step-wise, dose-dependent effects, pairwise comparisons were performed to test for statistically significant differences between adjacent TNF-α concentration levels within each respective day, as well as comparisons between day 2 and day 4 within each concentration. By utilizing a global linear model, the standard error used for these specific pairwise comparisons reflects the pooled variance across the entire dataset rather than only the isolated variance of the two groups being compared.

Additionally, to assess the overall effect of time on prime editing efficiency independent of dose, the main effect of the time point was evaluated. EMMs for each day were calculated by averaging across all tested TNF-α concentrations, and an overall pairwise comparison between Day 2 and Day 4 was conducted using a marginal means t-test. Significance thresholds were defined as * p < 0.05, ** p < 0.01, *** p < 0.001, and **** p < 0.0001.

### Xenograft model

For the in vivo NFkB recorder xenograft model, HEK cells expressing a NFKB controlled prime editor and the LTR4-RGR1 pegRNA were resuspended in PBS and mixed with Matrigel® (Corning, 356231) in a 1:2 volume ratio. 5x10^6^ cells were planted subcutaneously in the flank regions of J:NU athymic nude mice. After implantation, tumor size was monitored. LPS (Sigma, 437627) was administered via i.p. injection at designated time points at designated doses. We amplified LTR5-RGR1 from each tumor and sequenced with at least 500,000 reads per sample.

Cohen’s *d* was calculated as described above to evaluate relative resolution of recording with LTR5-RGR1. To determine if the combined effect size (’All’) differed significantly from the individual copy number site distributions, one-sample t-tests were performed. For each comparison, the Cohen’s *d* value of the ‘All’ group was compared against the distribution of values for the 1-copy and 2-copy groups independently. All statistical analyses were performed in R.

### Initial processing of sequencing data (for input to data analysis)

Paired end reads were merged using usearch fastq_mergepairs^101^. Reads without the forward and reverse primers were thrown out, and primers were trimmed from reads. Reads were merged to a list of sequences with the number of reads aligning to that sequence, and sorted in descending order by abundance. Sequences with less than two reads were discarded.

### Building LTR5 RGR1 reference amplicon list

We evaluated the top 5000 most abundant amplicons across all 24 of the negative control samples in the bulk LTR5 RGR experiment performed in the monoclonal TRE-deg-PE2 + LTR5-RGR1 pegRNA cell line. For each amplicon, we found its closest higher abundance sequence. First, if it had a Hamming distance (HD) of less than 2 with a more abundant amplicon, the amplicon with highest abundance with that HD was considered its closest amplicon. If the smallest HD was 2 or greater, we found the closest higher abundance sequence by finding its maximum alignment score (local pairwise alignment). We considered the top 10 most abundant amplicons as reference amplicons. After that, we removed amplicons as most likely resulting from sequencing error or template switching if: (1) its abundance ratio to its closest sequence was less than 0.01 and its HD with the closest sequence was 1 or its alignment score with its closest sequence was >= the length of the shorter amplicon minus 1.5, or (2) if its abundance ratio to its closest sequence was less than 0.05 and its HD with the closest sequence was 1 and the single mismatch was located in the first or last 5 nucleotides of the amplicon. This resulted in 329 out of the next 355 most abundant amplicons being kept as new reference amplicons. For amplicons with an abundance less than 1% of the abundance of the second most common amplicon, we additionally removed amplicons as most likely resulting from sequencing error or template switching if its abundance ratio to its closest sequence was less than 0.1 and its HD with the closest sequence was 1. This resulted in 39 out of the next 131 most abundant amplicons being added as new reference amplicons. Finally, after the top 500 most abundant amplicons, we removed amplicons as most likely resulting from sequencing error or template switching if its abundance ratio to its closest sequence was less than 0.05, regardless of its minimum HD or maximum alignment score. This resulted in 9 out of the next 61 most abundant amplicons being added as new reference amplicons. We stopped considering amplicons for being added to the reference list once their abundance dropped to less than 0.1% of the most abundant amplicon, which occurred after the 561st most abundant amplicon. The final list contained 387 reference amplicons, 335 of which contained the CRISPR target and 42 of which did not (**Table S2**).

### Copy number prediction for LTR5 rgRNA target sites

As described above, a set of reference amplicons was built from deep sequencing of wild type DNA. Reads from all single cell and bulk samples were aligned to this set of reference amplicons, and reads were called as edited if they contained the encoded GG to CC edit. Per amplicon editing observed for single cells in all treatment groups outside of the no treatment group were used to calculate copy number.

First, we calculated the average editing rate across single cells for each amplicon for each treatment group, to give us *p*, the probability that for a cell in a given treatment group we would see that amplicon edited. Then, assuming 100% detection rate, for an amplicon with a copy number of 1 (*c = 1*), we would expect to see all of the reads aligning to that sequence to be edited in a single cell in a treatment group with a probability of *p*, and to see all reads be unedited with a probability of *q* (where *q = 1-p*). For an amplicon with a copy number of 2 (*c=2*), we expect reads aligning to the amplicon to be fully edited *p^2^* of the time, fully unedited *q^2^* of the time, and partially edited *2pq* of the time (meaning one copy is edited, one copy is unedited); and so on. For each amplicon, single cells were classified into three discrete observable states based on their editing frequency: fully edited (>0.96), fully unedited (<0.04), and partially edited (0.04 <= x <= 0.96).

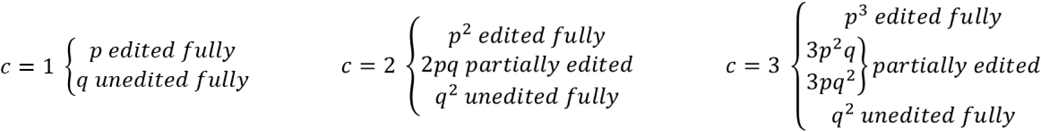

In this manner, as the copy number c increases, the number of “partially edited” cells expected increases, while the expected number of cells where that amplicon is either fully edited or unedited decreases.

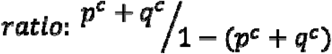

To account for technical dropout during sequencing, a detection rate parameter *d* was introduced, representing the probability of successfully detecting an existing copy. The observed state of a cell is thus conditional on both the true number of edited copies and the detection rate. Because an amplicon is included in the dataset only if at least one copy is detected, the probabilities are normalized by the factor *1 - (1-d)^c^*. Based on the observed sequencing coverage, we predicted a dropout rate of about 40%; as a result, we made our calculations using a detection rate of 0.6. For a copy number of 2 (*c = 2*), we would expect sequencing to capture each copy of the amplicon in 60% of single cells sequenced (*d = 0.6*). Factoring that detection rate in, our new predicted distribution of observed reads became:

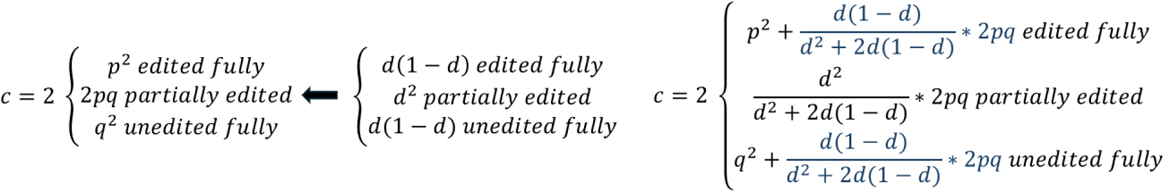

For any given copy number *c*, this equation becomes (where *k* represents the true number of edited copies (ranging from *1* to *c-1*):

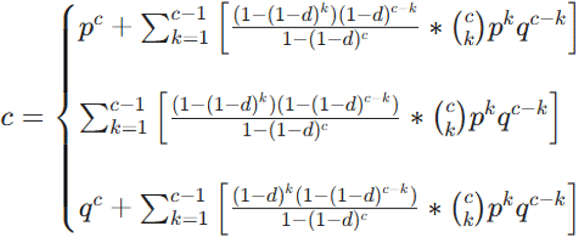

The expected number of cells for which an amplicon was fully edited (*e_exp_*), partially edited (*p_exp_*), or fully unedited (*u_exp_*) in each treatment groups if it had copy number *c* was calculated as follows. In these equations, *N* is the total number of single cells in that treatment group for which the amplicon was covered, and *p* is the editing rate for that amplicon in that treatment group:

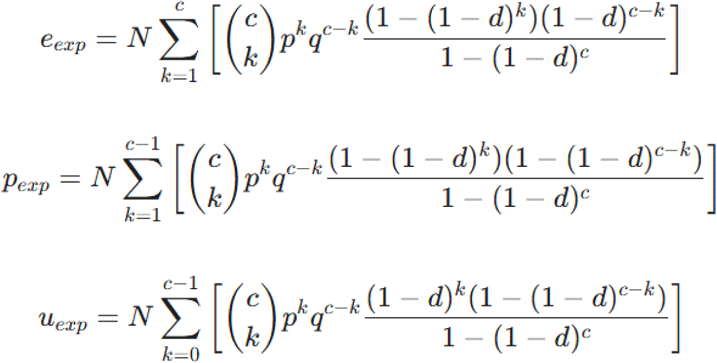

To evaluate the fit of each hypothesized copy number *c*, a modified Chi-square (□^2^) statistic comparing the observed and expected counts was used. In a strict mathematical model of a single-copy amplicon, the expected count for partially edited cells is exactly zero. However, standard single-cell data contains noise and bias/technical confounders (e.g., droplet doublets, alignment artifacts) that might result in amplicons being incorrectly called as partially edited. Standard □^2^ calculations result in an undefined or mathematically inflated penalty when dividing by an expectation of zero, leading to the systematic misclassification of true single-copy amplicons. To resolve this, we used a modified penalty function implementing a soft-penalty threshold when calculating □^2^ for *c = 1*.

The chi-squared statistic for the number of cells with the amplicon fully edited (*e_obs_*), partially edited (*p_obs_*), or fully unedited (*u_obs_*), with copy number c, was calculated as:

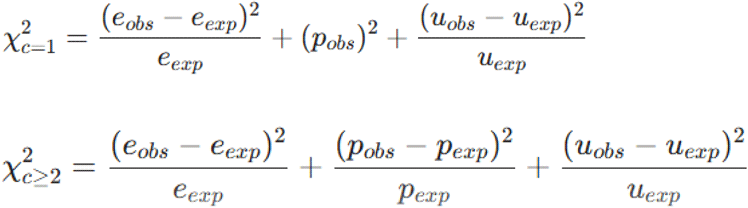

These □^2^ are calculated for each treatment group, and the sum across all five treatment groups is taken as the final □^2^ for that amplicon having copy number *c*. The mathematically assigned copy number for the amplicon is the copy number *c* that minimized the final □^2^.

In a subset of amplicons, editing efficiency was negligible across all experimental groups (e.g., >99.5% of cells remained fully unedited). Under these conditions, the observed data perfectly matches the mathematical expectation of the fully unedited state for any hypothesized copy number, resulting in the □^2^ for all copy numbers to be equal; as a result, the current algorithm fails to estimate the copy number for these amplicons. To address this, if the aggregate fraction of fully and partially edited cells across all subpopulations for an amplicon was below 0.5%, the binomial modeling was bypassed. In these cases, the copy number of that amplicon was instead estimated by calculating the integer median of the final consensus copy numbers from the 5 confidently assigned amplicons closest in sequencing abundance. This localized abundance proxy allowed for robust copy number estimation even in the complete absence of an active editing signal.

### LTR5 prime-editing analysis (aka processing of in vitro experimental data)

A set of unedited reference sequences/amplicons was built from deep sequencing of DNA from a nontreated population of cells from the D7 monoclonal iiPE2 + LTR5 peg-rgRNA cell line. An edited set of reference sequences was built by making an edited version of each unedited reference sequence with the specified prime edit. Reads were aligned to the combined set of unedited and edited reference sequences, and were called as “edited” or “unedited” based on whether they aligned with a higher score to an edited or unedited reference sequence. Reads that aligned equally to multiple reference amplicons were discarded, and sequences with an alignment score less than the length of the shorter sequence minus four were discarded. This alignment score cut off of the length of the shorter sequence minus four was set to reduce misaligned reads contributing to skewing the per amplicon editing fractions for low copy number amplicons that might negatively impact classification per amplicon editing fraction based classification algorithms, while keeping reads with one or two errors likely from sequencing.

### Single cell classification using average editing across all LTR5 RGR target amplicons - aggregate Z-score method

Prime editing of LTR5 RGR target amplicons for each single cell was quantified as described above. The average editing per cell for all cells in each treatment group was used as the population mean for that treatment group. To classify each single cell, the Z-score was calculated between the cell and each treatment population, and the cell was assigned to the treatment group for which it had the smallest Z-score.

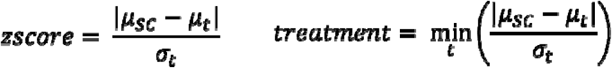

Normalized confidence scaling for aggregate editing: As a measure of confidence in classification for the aggregate Z-score method, we converted the raw Z-score distances into a probabilistic scale using a Softmax-based normalization. Because the no treatment group exhibited a significantly tighter distribution (*σ = 0.3*) compared to the other treatment groups (*σ = 8.3, 15.2, 11.4* for the low, mid, and high treatment groups), a simple linear distance (margin) was insufficient to represent biological certainty. The Z-scores calculated as described above were transformed into a probability distribution by applying the exponential of the negative Z-scores, where:

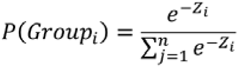

This transformation ensures that smaller Z-scores (shorter distances to a group mean) result in higher probabilities. The resulting Softmax Z-score Probability represents the model’s relative confidence in its top-ranked prediction on a 0.0 to 1.0 scale, making the confidence mathematically comparable across the Z-score, Random Forest, and XGBoost frameworks.

### Random Forest Classification Framework

To classify single cells into treatment groups based on multiplexed recording data, we implemented a Random Forest (RF) classifier. The input features consisted of the editing fractions for 291 unique amplicons per cell, for the 747 single cells across four treatment groups (“None”, “Low”, “Mid”, and “High”). Missing values (NAs) in the amplicon editing matrix resulting from stochastic sequencing coverage were handled using a Zero-fill strategy. Under this biological assumption, missing data is treated as a lack of editing signal (0). This conservative approach prevented classification being biased towards higher treatment classification and maintained the integrity of the negative control group. The RF model was trained with the following parameters:

- n_estimators = 200: The number of decision trees was set to 200 to ensure a stable "majority vote" across the forest, reducing the influence of noisy, low-copy-number amplicons.
- max_depth = None: Trees were allowed to grow to their full depth. Given the complexity of genomic recording signatures, full depth is appropriate here to capture high-order interactions between amplicons, which is necessary for resolving complex "Mid" vs. "High" signatures.
- max_features = ‘sqrt’: By restricting each split to a random subset of amplicons (square root of 291), we forced the forest to build diverse trees that do not rely solely on a few dominant features.

Confidence in classification per cell was calculated as the difference between the highest and second highest probability. Model performance was evaluated using Leave-One-Out (LOO) cross-validation. In this iterative process, each individual cell (n=747) is held out as a test case while the model is trained on the remaining 746 cells. Since every single prediction is made on a cell that the model was never allowed to see during that specific training fold, this provides a very robust estimate of how the model performs on "unseen" data. This approach ensures that every cell is tested against a model trained on all other available data, so the reported accuracy is a true reflection of the model’s ability to generalize to new single-cell data.

### Gradient Boosted Classification Framework (XGBoost)

To classify single cells into treatment groups based on multiplexed recording data, we implemented an XGBoost (Extreme Gradient Boosting) classifier. The input features consisted of the editing fractions for 291 unique amplicons per cell, for the 747 single cells across four treatment groups (“None”, “Low”, “Mid”, and “High”). Missing data/stochastic sequencing dropout was handled natively; rather than imputing missing values (NAs) with zeros or medians, the model learned an optimal "default path" for cells with missing amplicon data. This approach prevents the introduction of artificial bias and allows the model to treat the absence of sequencing data as a potentially informative biological or technical signal. The model was trained with the following parameters to ensure robust classification:

- learning_rate = 0.1: The learning rate controls how much each successive tree contributes to the ensemble; a learning rate of 0.1 balances learning efficiency against over-correction for stochastic biological noise. Higher values risk overshooting optimal solutions by over-fitting to noisy single-cell outliers, while lower values require more trees and risk underfitting subtle treatment boundaries. Setting the learning rate to 0.1 provides sufficient shrinkage to prevent any single tree from dominating predictions, allowing the model to converge on stable biological signatures across amplicons.
- n_estimators = 200: The ensemble size was set to 200 estimators, a threshold determined to allow for model convergence without reaching the point of diminishing returns or overfitting. 100 iterations were often insufficient to fully resolve the complex signatures of the upper treatment groups; accuracy was still climbing at 100. Increasing the number of trees to 300 only increased training accuracy without helping the Leave-One-Out test scores, indicating the point of overfitting was reached.
- max_depth = 7: Allowing a depth of 7 enables the model to capture complex, non-linear interactions between up to seven different genomic sites. A depth of 7 allows the model to consider interactions between up to 2^7^ = 128 leaf nodes. Biologically, this is more than enough to capture the multiplexed signatures of 291 amplicons. Unlike Random Forest models, where full-depth trees are averaged, the XGBoost depth was capped at 7 to prevent noise propagation. Since trees are built sequentially to correct prior errors, deeper trees risk amplifying outlier-driven patterns rather than signatures shared by many cells.
- colsample_bytree = 0.4: Forces each tree to ignore 60% of available amplicons at each split. This prevents high-copy-number "master" amplicons from drowning out the subtle signals provided by lower-copy genomic sites, so that no one feature dominates the rest).
- gamma = 0.1: This regularization parameter sets a minimum loss reduction threshold for creating new branches. The model will only create a new branch if the gain for the split is > 0.1, acting as a quality control filter that prevents hyper-specific branches built on single-cell noise rather than treatment-level patterns.
- subsample = 0.8: Training each tree on a random 80% subset of the cells further protected the model against overfitting to individual cell outliers.

Confidence in classification per cell was calculated as the difference between the highest and second highest probability. Validation followed a similar process for XGBoost as random forest. Model performance was validated using Leave-One-Out (LOO) cross-validation on 747 cells. This iterative process ensures that every cell is treated as an "unknown" sample, providing an accurate estimate of how the model would perform on future single-cell recording datasets.

To demonstrate that the classification accuracy was not attributable to spurious patterns in noise, we used Y-Randomization (or Permutation Testing). This process involves shuffling the "Actual Treatment" labels while keeping the amplicon data exactly the same. If the model is truly learning biology, its accuracy should drop to the level of random chance (∼25% for four groups) when the labels are scrambled. The Y-Randomization (Permutation) runs 100 iterations of the classification with shuffled labels to create a "null distribution" of accuracy. The average “null accuracy” across the 100 iterations was ∼30% with an empirical p-value of 0.0, meaning that in 100 attempts at finding patterns in "nonsense" data, the model never once came close to the real accuracy of 85.0%. Even though the null accuracy is 30%, the 55% gap between "noise" and "signal" is large. This confirms that the 85% accuracy is driven by specific biological signatures tied to the treatment groups, not just the underlying distribution or structure of the editing matrix, and that the XGBoost model is successfully identifying specific, non-random editing thresholds and multiplexed amplicon combinations that are uniquely characteristic of each treatment intensity.

### Single cell classification algorithm resolution comparison

Classification accuracy across the three approaches correlated with confidence separation between correct and incorrect predictions. Cohen’s d was calculated as the mean confidence for correct predictions minus the mean confidence for incorrect predictions, divided by the pooled standard deviation (calculations below).

Min Z-score: (0.729-0.690) / 0.102 = 0.382

Random Forest: (0.554 - 0.268) / 0.266 = 1.075

XGBoost: (0.849 - 0.562) / 0.242 = 1.186

### Simulations to predict single cell signal recording capabilities using subset of LTR5 sites

For single amplicon classification, we filtered single cells for each amplicon independently to exclude cells where that amplicon was not covered by sequencing (NaNs). For the naive minimum Z-score classification method, the local mean and standard deviation of editing fractions was first calculated for each treatment group within the covered cells. In the case that cells within a treatment group exhibited zero variance, a baseline standard deviation of 1E-9 was used instead of 0 to prevent mathematical division by zero. Then, as was done when considering all amplicons together, each cell was classified by calculating its Z-score distance to each treatment group’s mean and assigning it to the group with the minimum distance. For Random Forest (RF) and XGBoost, we used the same parameters with the scikit-learn RandomForestClassifier and XGBClassifier as was used when considering all amplicons as described above. Classification accuracy for each amplicon was then evaluated the same as it was when considering all amplicons, using leave-one-out (LOO) cross-validation across all valid cells for each amplicon independently. Then, the resulting total classification accuracies were stratified by each amplicon’s previously estimated genomic copy number.

To compare classification accuracy using the highest performing classification method (XGBoost) between an amplicon with *n* copies and *n* single-copy amplicons, we compared the results for single amplicon classification as described above to classification using *n* single-copy amplicons. For each copy number *n*, we executed 1000 independent simulation iterations. For each iteration, a cell was selected at random, and *n* single-copy amplicons were selected randomly without replacement from the single copy amplicons covered in the selected cell. Then, an XGBoost model was trained using the same parameters as defined above on the remaining cells using only the *n* sampled features to predict the held-out test cell. The proportion of correct classifications across all 1,000 simulations was used to establish the mean accuracy for each number of distinct targets. For each copy number *n*, we executed as many 1,000 simulations as there were amplicons with that copy number, so the number of data points for each copy number considered in this analysis are equal.

To evaluate the added benefits of increasing the number of sites used for recording in a single cell, we simulated classification with increasing total copies considered by tracking classification accuracy against the total cumulative copy number of randomly sampled amplicon subsets from randomly selected cells. For each simulation, a cell was selected at random, and amplicons were sampled iteratively at random from those covered for that cell until the sum of the selected amplicons individual copy numbers fell within a specified range or reached a specified maximum cumulative copy number threshold. Then, either the XGBoost or Random Forest method was used as described previously to classify the randomly held-out test cell using the randomly sampled subset of amplicons. The average classification accuracy and standard error for each total cumulative copy number was calculated from the ratio of correct to incorrect predictions across the thousands of simulations for each copy number. A minimum of 1,000 simulations were executed for each total copy number.

### Pipeline for systematic identification of RGRs (Figure S3)

The methods described in the sections below detail the methods used within the repeatable RGR-finding workflow. As a reminder, Figure S3 illustrates the full workflow at a high level.

#### Part 1: Identifying guides targeting genomic repeats

The first component of the pipeline identifies and extracts genomic repeat sequences from a reference genome of interest. To identify repeats locations, we used Repeat Masker, which identifies interspersed repeats and low complexity DNA sequences from the reference genome of an organism; pre-computed RepeatMasker annotation files are publicly available for model organisms, including all species studied in this paper^53^. The RepeatMasker output contains start and end coordinates along with detailed annotations for each identified repeat region, including repeat class and family^53^. The pipeline takes these coordinates and extracts the corresponding sequences from the reference genome using the bedtools command getfasta, producing files containing all sequence mapped to each repeat. Simple repeats (i.e. (TTTC)_n_), satellite repeats (groups of repeating short DNA sequences, primary component of centromeres), and low complexity repeats (i.e. A-rich region) were excluded from downstream analysis. (code: parseRepeatMaskerData.py).

Next, sequences extracted from repeats annotated on the reverse-complement strand of the reference genome were reverse complemented for consistency to simplify identification of spacers (code: multAlign-repeatMaskerOutput.py). The sequences for each repeat region are then systematically parsed for CRISPR guides, defined as any 20 nucleotide sequence adjacent to a NGG PAM site. The identified guides are then sorted into output files based on predicted on-target efficiency scores calculated using the CRISPRater algorithm (crisprator citation- https://github.com/crisprVerse/crisprScore). This algorithm outputs a number between 0 and 1 representing the probability that a guide RNA with a given 20 nucleotide sequence will cut at its intended target. We required a minimum CRISPRater score of 0.55 to move forward in the pipeline, eliminating guides that were unlikely to successfully target where intended in the genome. (code: parseRepeatSpacers.py). In the end, all of these were combined into a summary file listing all guides targeting repeats with the number of perfectly matching targets it has throughout the repeat genome.

#### Part 2: Define sites targeted by each guide and filter out guides targeting functional regions

Guides are then screened for off-target sites - genomic locations that may be targeted by the guide RNA despite slight deviations from the exact guide sequence. Accounting for off-targets is critical, as failure to do so could lead to gross underestimation of the number of sites being targeted, may result in missing edited sites during downstream analysis, and risks overlooking unintended exon-targeting guides that would otherwise be excluded. Off-target screening was performed using CasOFFinder^61^, to get a complete list of all genomic locations potentially targeted by each guide, allowing for up to 2 mismatches. This list was further filtered to exclude potential targets with mismatches in the seed region (the 6 nucleotides closest to the PAM site). This ran in tandem to screening all identified target sites against exon coordinates obtained from UCSC Table Browser annotation files as described below; guides with any target site overlapping an exon were excluded to avoid unintended editing of functional regions. The output contains coordinates for all sites targeted by each *non-exon* targeting guide, along with a summary file indicating the total number of expected target sites per guide. (code: run casOffinder command, then filterCasOFFinderOutput.py, getCasOFFinderData.py and getSpacerLocs.py).

Then, using annotation files for the given genome, all sites targeted by guides were evaluated for functionality. The pipeline uses UCSC known gene annotations (details below) to identify exonic and intronic regions in all three organisms, and to generate bed files annotating the 500 nucleotides upstream of all non-pseudogene transcription start sites. These annotation files were downloaded from UCSC table browser^102^ from the following tracks:

Mouse: Group = Genes and Gene Predictions, Track = GENCODE VM23, table = knownGene.

Human: Group = Genes and Gene Predictions, Track = GENCODE V49, table = knownGene.

Zebrafish: Group = Genes and Gene Predictions, Track = NCBI RefSeq, table = knownGene AND Track = Ensembl Genes, table = ensGene

To generate the exon annotation file, one BED record was produced per exon; to generate the intron annotation file, one BED record was produced per intron. To get the 500 nucleotides upstream of transcription start sites for known genes excluding pseudogenes, in the filter page the "allow selection from this table" checkbox for “knownAttrs” was selected, and the “transcriptType” field was set to “does” match “protein_coding lncRNA antisense” to exclude pseudogenes. Many of the regions in this annotation file overlapped, since it included one entry per transcript and many transcripts share transcription sites. However, this redundancy was accounted for during scoring, since only one overlap between a target site and an annotated 500 bases upstream of a known transcription start site was required for that target site to receive the corresponding annotation. Having more than one overlap does not change this annotation or compound the penalty. There were 89,740 non pseudogene transcripts for mm10, and 416,798 non pseudogene transcripts for hg38.

The pipeline also uses candidate cis-Regulatory Elements (cCRE) from ENCODE (926,535 human cCREs and 339,815 mouse cCREs), which were downloaded directly from ENCODE. Since this sCRE annotation file is absent for zebrafish, curated annotation tracks were obtained from the DANIO-CODE Consortium to get a uniquely identifiable map of curated cis-regulatory elements across the zebrafish genome. Data was accessed via the UCSC Genome Browser using the danRer11 (GRCz11, May 2017) assembly. The DANIO-CODE public track hub (https://trackhub2.genereg.net/DANIO-CODE/DANIO-CODE.hub.txt) was attached to the UCSC Table Browser to export five processed regulatory tracks in BED format: Consensus Predicted ATAC-seq-supported Developmental Regulatory Elements (cPADREs), Dynamic Orphan Predicted Elements (DOPEs), Constitutive Orphan Predicted Elements (COPEs), experimentally validated transgenic enhancers, and the Promoterome Atlas consensus promoters. The five BED files were first processed individually; the genomic coordinates were sorted positionally (bedtools sort), and any overlapping intervals within the same class were merged (bedtools merge) to prevent redundant counting of overlapping features. Then, a standardized, class-specific identifier was added to the fourth column (name field) of each file - cPADRE (Consensus Predicted ATAC-seq-supported), dope (DOPE = Dynamic Orphan Predicted Element), cope (COPEs = Constitutive Orphan Predicted Element), transgenic_enhancer, and consensus_promoter - and they were concatenated into a single dataset (zeb-DANIO-annotations.bed).

Bedtools intersect was used with the following parameters to identify overlap between target sites and functional annotation files:

To check overlap with exon annotation bed files:

bedtools window -w 20 -a casOFFinder-output-sorted.bed -b annotations-sorted.bed -u > overlaps.bed

To check overlap with introns and 500 nucleotide upstream annotation bed files:

bedtools intersect -wa -wb -a casOFFinder-output-sorted.bed -b annotations-sorted.bed -sorted > overlaps.bed

To check overlap with cCRE annotation bed files:

bedtools intersect -wa -wb -a casOFFinder-output-sorted.bed -b cCRE-annotations-sorted.bed -sorted | bedtools groupby -g 1,2,3,4 -c 10 -o distinct -delim ";" > overlaps.bed

Any of the guides targeting repeats that had a single target site overlapping an exon (+/- 20 nucleotides) was eliminated as soon as casOFFinder identified the exon overlapping target, as described in the previous step. The remaining guides’ target sites were screened against the rest of the annotation files, and the results of the screen/resulting overlaps were used to generate a functionality hazard score for each rgRNA. For human and mouse, a target overlapping with a cCRE promoter like signature (PLS), proximal or distal enhancer like signature (pELS/dELS), or the region 500 nucleotides upstream of a transcription start site (for all genes except except pseudogenes) was counted as 1, and a target not overlapping any of those but overlapping a cCRE DNase-H3K4me3 and/or CTCF bound annotation was counted as 0.5, and these numbers were added across all targets to create a functionality hazard score (FHS). For zebrafish, a target overlapping with the region 500 nucleotides upstream of a transcription start site (for all genes except except pseudogenes) or a DANIO annotated consensus promoter, transgenic enhancer, or cPADRE was counted as 1, and a target not overlapping any of those but overlapping a DANIO annotated COPE or DOPE was counted as 0.5, and the numbers were summed across all targets as before to get the FHS. The functionality hazard percentage (FHP) is calculated from the functionality hazard score (FHS) by dividing it by the total number of targets and represents the fraction of targets which can potentially interfere with a functional element (code: annotateFunctionalityScore.py).

Then, guides that shared over 60% of their target sites were merged to the top guide by the following hierarchical selection algorithm. The selection process was designed to prioritize the dual objectives of minimizing functionality hazard percentage, and maximizing the number of exact targets, using the CRISPRator score as a final tie-breaker. To do this and identify the "best" item from the list of options, candidates were evaluated through seven sequential priority tiers. The algorithm terminated as soon as a tier yielded a viable candidate or a set of identical top-performing candidates. In the first tier, the algorithm sought candidates that simultaneously achieved the absolute minimum functionality hazard percentage and the maximum number of exact targets. If no such candidate existed, the second tier relaxed the accuracy constraint slightly, looking for guides with the minimum functionality hazard percentage who had one less exact target than the maximum. The third tier further expanded the search among the minimum functionality hazard percentage guides by identifying guides with the highest percent of exact matches out of total sites targeted, within 10% of the global maximum, while still prioritizing the highest available exact value within that subset.

If the first three tiers yielded no results, the search scope shifted to a "near-optimal" functionality hazard percentage range, defined as any functionality hazard percentage within 10% of the global minimum. Within this expanded functional pool, the fourth tier prioritized maximizing the number of exact targets. If this was not met, the fifth tier looked for guides within the pool for guides with one less exact target than the maximum. The sixth tier looked for guides within the near-optimal functional range that had the highest percent of exact matches out of total sites targeted, within 10% of the global maximum and again prioritizing the highest exact score. Finally, if all previous criteria remained unfulfilled, the seventh tier selected the candidate from the near-optimal functional group that possessed the highest local exact value. This tiered architecture ensured that minimizing functionality hazard percentage remained the primary gatekeeper.

If any single tier produced multiple qualifying candidates, a secondary resolution protocol was used. Candidates were first ranked by percent of exact matches out of total sites targeted, with ties broken by CRISPRator score. If multiple guides remained indistinguishable after this step, the algorithm selected one of the top tied options randomly (code: mergeNonexonicGuides.py and mergeNonexonicGuides-pt2.py).

#### Part 3: Search for conserved regions up and downstream of target sequences and design primers to maximize amplification of target sites

For each guide, the pipeline uses Clustal Omega multiple sequence alignment^64^ to align all sequences targeted by that guide (multAlign-filteredInGuides.py). It then screens for potential conserved regions up and down stream of the target sequence that could serve as primer binding sites, for the design of primers that maximize amplification of all targeted sites (primerScreenCheck-final.py). This screen is based on a consensus sequence of the alignment with a threshold of 0.7. This means that the screen requires 70% of sequences to have the same nucleotide in a position for that nucleotide to be in the consensus sequence; positions lacking that consensus are denoted as “X”. The primer screen requires the presence of at least one 15 nucleotide stretch of consensus sequence up and downstream of the conserved Cas9 target, with 100 nucleotides between them. This ensures that for any guide passing the filter, a minimum of 40% of its target sites have conserved regions both up and downstream to serve as potential primer binding sites.

Potential primer options were identified using a custom bioinformatic pipeline designed to find potential primers 18-30 bases long, and select and pair forward and reverse primers relative to a target 20 bp spacer sequence that have the potential to maximize amplification of targets. First, the target site was identified within the Multiple Sequence Alignment (MSA) from the primer screen check step by generating a majority-rule consensus string. To account for sequence variability at the target site, a sliding window search was performed across the consensus using Hamming distance. A window was identified as the target site if it exhibited ≤2 mismatches compared to the spacer, ignoring gap characters. In cases where a strict Hamming search failed, a local alignment was utilized as a fallback to identify the target index. The local alignment assigned a score of +1 to matches, -1 to mismatches, and -1 to gaps.

Then, potential forward and reverse primers were identified within the alignment regions upstream (left) and downstream (right) of the spacer, respectively. A sliding window generated all possible k-mers with lengths ranging from 18 to 30 nucleotides. Candidates were scored based on their physicochemical properties and thermodynamic stability, and each was assigned a quantitative quality score based on its sequence composition, thermodynamic profile, and structural complexity. This score was used to differentiate primers with high binding specificity and extension efficiency from those likely to form secondary structures or exhibit non-specific binding. The score represents the sum of these individual components as described:

- Nucleotide Diversity and Complexity: Sequence diversity was calculated to penalize primers with biased base compositions or low complexity. The frequency of each nucleotide (A, C, G, T) was calculated relative to the total primer length. A "diversity score" was assigned based on these frequencies. If more than one nucleotide type was completely absent (e.g., a primer made only of A and G), a diversity score of -10.0 was applied to the total score, effectively disqualifying low-complexity sequences that are prone to non-specific binding or structural issues. If not, the final diversity score applied to the total score is assigned based on the minimum frequency observed among the four bases: if the minimum frequency< 0.02 (highly skewed), the diversity score was -3. If the minimum frequency< 0.1 (moderately skewed), the diversity score was -1. And if the minimum frequency was >= to 0.1, the diversity score is the minimum frequency (a positive decimal).
- Repetitive Elements: To prevent primer dimerization and non-specific binding in homopolymeric regions, the algorithm searched for repetitive stretches. Progressive penalties were applied for homopolymers (A, T, C, or G) exceeding 4 bases, with penalties scaling from -1.5 for 5-base repeats up to -10.5 for repeats of 7 bases or more.
- 3’ Terminus Composition and GC Content: the composition of the 3’ end was specifically weighted to optimize polymerase docking and extension. To ensure stable binding at the initiation site of extension, a bonus of +2.0 was applied to the final score if the terminal 3’ nucleotide was a G or C. The terminal 8 nucleotides were also evaluated for their GC vs AT content. Regions with moderate GC enrichment (4 to 6 GC bases) received a +1.0 bonus, while low GC content (<4 GC bases) resulted in a -1.0 penalty. Optimal global GC content was targeted between 40% and 60% (+1.5 bonus), with a secondary tier of stability between 61% and 75% (+0.8 bonus).
- Thermodynamic and Structural Parameters: Primers between 20 and 25 nucleotides were prioritized (+1.5 bonus), while those exceeding 25 nucleotides were penalized (-1.0). The intrinsic melting temperature (*T_m_*) of each k-mer was calculated using the nearest-neighbor method (via the Biopython Nearest-Neighbor model). min(T_m, 65) - 59.0 was added to the overall score, favoring primers with higher T_m_s and penalizing primers the lower their T_m_.

The pipeline utilized a two-tiered stringency approach to loosen the thresholds to suggest viable pairs even in alignments of targets and surrounding regions that were more divergent, and didn’t pass the first round of thresholding. First, initial candidates were required to have an intrinsic melting temperature Tm ≥ 55∘C and a conservation threshold ≥85% across the alignment matrix. For each potential candidate, conservation was determined by iterating through all individual sequences within the local multiple sequence alignment matrix. A sequence was defined as a ‘Match’ for a specific primer only if it satisfied two concurrent criteria:

1. 3’ Terminal Fidelity: The sequence was required to possess an identical nucleotide at the alignment column corresponding to the primer’s 3’ hydroxyl terminus, ensuring the structural prerequisite for polymerase-mediated extension.
2. Sequence Identity: For sequences passing the 3’ anchor check, a Hamming distance was calculated between the k-mer and the alignment window. Sequences were permitted a maximum of two mismatches (Hamming distance ≤ 2). To account for small genomic indels, a penalty was applied to the distance score for length differentials exceeding a single nucleotide, equal to the difference in lengths.

If the search with these parameters yielded zero final pairs, the pipeline automatically re-initiated discovery with relaxed parameters: the intrinsic melting temperature Tm threshold was lowered to 52∘C, the conservation threshold was reduced to 70% across the alignment matrix, and the sequence identity criterion for conservation was relaxed, to require a maximum Hamming distance for sequence matching of 3 (instead of 2). If none of the options with Tm ≥ 52∘C and a Hamming distance for sequence matching ≤ 3 passed the 70% conservation threshold, it returned the single Tm-mer that had the highest conservation, to provide one option to test as a potential primer.

Then to reduce redundancy, overlapping primer candidates that started within 5 bp of each other were merged to select a representative potential primer for any given potential primer binding site. Within each group, this single representative was selected based on a primary priority for conservation and a secondary priority for the physical score, ensuring that the selected candidates possessed both the sequence conservation required for target recovery and the physical characteristics required for robust PCR performance. Final primer pairs were formed by combining forward and reverse representatives that produced a minimum amplicon length of 100 bp. To ensure experimental viability, each pair was evaluated for potential secondary structures (hairpins and heterodimers). Pairs exhibiting a predicted Gibbs free energy (ΔG) lower than -9.0 kcal/mol were excluded to prevent primer-dimer formation. A threshold of -9.0 kcal/mol was used because it is a common industry heuristic used in software like Primer3 and IDT’s OligoAnalyzer. It is based on the thermodynamic principle that as ΔG becomes more negative, the stability of the dimer increases. At values more negative than -9.0 kcal/mol, the primer-dimer species becomes stable enough at typical PCR annealing temperatures (55-65^∘^C) to compete significantly with the target template for reagents, leading to reduced yield or non-specific amplification. Pairs were ranked by combined quality scores, and all results were exported for further analysis (next step). (code: findAndProcessPrimers.py)

#### Part 4: Evaluate potential primer pairs and select optimal primer pairs per guide

To optimize computational efficiency during the process of screening potential primers for the optimal pair per guide, a secondary filtering protocol was implemented to limit the number of unique primers subjected to a heuristic genome-wide search for binding sites. Primer pair candidates were generated from consensus sequences as described in part 3 above. These candidates were ranked first based on the conservation of their binding sites up and downstream of target sites. This conservation was calculated as the sum of the conservation score for the forward and reverse primers from the previous step. Candidates were then ranked based on a secondary conservation score characterizing the conservation across the three most 3’ terminal nucleotides of the primer binding sites across all target containing amplicons. This secondary conservation metric was defined as the percentage of sequences in the alignment matrix where the three most 3’ terminal nucleotides of the primer matched the template exactly. Finally, candidates were ranked based on the score of the top two primers as described in the previous step. In the case more than 50 primer pair options were suggested, only the top 50 ranked pairs per spacer were evaluated. To further prevent redundant processing, unique forward and reverse primer pairs were considered as follows. The first five unique forward and reverse primers identified in the top-ranked pairs were automatically advanced. Subsequent primers (ranked 6 through 10) were only included if their individual 3’ terminal conservation score (the secondary conservation metric described above) remained within 15% of the absolute maximum observed conservation for primers in that direction being considered. Finally, the unique forward and reverse sets were capped at a maximum of 10 primers each per guide.

To identify all potential binding sites across the genome for each forward and reverse primer, a high-throughput heuristic search was conducted using UCSC Genome Browser’s findMotif utility. All unique candidate primers were mapped to the genome using their 3’ terminal 15-nucleotide "seed" sequences as the search motif. The search was parameterized to allow a maximum of two internal mismatches (mismatch=2) within this 15-bp seed region (./findMotif genome -motif=PRIMER_LAST15NT -misMatch=2). This threshold was selected to maintain a balance between capturing high-affinity off-target sites and mitigating the excessive computational noise generated by low-identity non-specific binding. To ensure comprehensive capture of all potential binding sites, the search was executed independently against both the original primer motif and its reverse complement. This dual-pass approach is critical for identifying potential amplicon partners located on opposite strands, since findMotif only returned perfect matches on the opposite strand when executing a search for a given motif, rather than allowing for the 2 mismatches directed in the command. The resulting strand-specific hit lists, generated in BED format, were merged into a unified genomic binding map. During this integration, hits identified on the reverse complement strand underwent a sign-reversal (strand flipping) to maintain coordinate consistency relative to the reference genome, and the number of 5’ nucleotides of the primer that had been left off of the 15 nucleotide motif was added to the coordinates, so the bedfile contained coordinates for the entire potential primer binding site, not just the 3’ most 15 nucleotides. Duplicate entries arising from the perfect matches that were identified by the search on both the original and reverse complemented primer motif were identified and removed to ensure a unique set of binding candidates.

Following the initial heuristic search, potential hits were subjected to a secondary filtration for polymerase extension eligibility. The Bedtools getfasta function was used to get the sequences of those coordinates. Hits were strictly required to possess a perfect 3’ terminal match between the primer’s hydroxyl terminus and the template. If the next 3’ most nucleotide did not match, the rest of the primer sequence had to have a perfect alignment to the template to be considered a hit - meaning it was required to have a minimum alignment score of the length of the primer minus two. If the two 3’ most nucleotides matched, but the third 3’ most nucleotide did not match, the rest of the primer sequence had to have a near perfect alignment to the rest of the template to be considered a hit - meaning it was required to have a minimum alignment score of the length of the primer minus four. And finally, if all three of the 3’ most nucleotides matched, the primer sequence was only required to have a minimum alignment score of the length of the primer minus five to be considered a hit. Scoring parameters for the pairwise aligner were: mode = global, match = 1, mismatch = -1, open gap score = -1, extend gap score = -0.2, and target and query end gap scores = 0.

Following the identification of all potential genomic binding sites for all forward and reverse primers individually, potential primers were paired to form "Expected Amplicons" based on their spatial orientation. A valid amplicon required a Forward (or Left) primer hit and a Reverse (or Right) primer hit to be located on opposite strands in a convergent orientation (facing each other), within 800 base pairs of each other. For every valid predicted amplicon, its genomic sequence (including the binding sites themselves) were extracted (again using Bedtools getfasta function) to generate the "Expected Amplicon" sequence. This sequence was then used for downstream specificity and target recovery analysis. Any potential primer pairs with zero predicted amplicons are immediately eliminated.

To quantify how well each primer pair recovered the desired target sites, the pool of predicted amplicons for each primer was compared against a bedfile of target sites (the 20 nt target sites). A target site was classified as "Recovered" by a primer pair if one of its predicted amplicon’s genomic start and end coordinates completely encompassed the start and end coordinates of the target site. Any predicted amplicon for a primer pair that did not capture a target site was classified as an "Off-target" product. These off-target counts serve as a critical metric for predicting sequencing library waste and non-specific background noise. For each case where a predicted amplicon covered a target site, the binding metrics (Match/Length and calibrated *T_m_*) for both the Forward and Reverse primers to that specific amplicon were recorded and formatted as a single descriptive string (e.g., 23/23 (59.8C) | 22/23 (54.8C)). *T_m_* was calculated using standard Nearest-Neighbor parameters (using Bio.SeqUtils MeltingTemp Tm_NN(primer, target_complement, dnac1=250, Na=50)); for primer-template duplexes containing non-canonical mismatches exceeding standard thermodynamic tables, T_m_ was estimated using a calibrated penalty of 1.5°C per mismatch relative to the perfect-match Nearest-Neighbor baseline.

Identification of the final optimal primer pair for each spacer was conducted through a hierarchical, multi-objective selection algorithm designed to balance recovery of target sites (sensitivity) with sequencing efficiency (specificity). First, a binary filter was applied to ensure a minimum Covered-to-Offtarget (C:O) ratio of 0.01. This gate ensured that the selected pair would yield at least 1% on-target amplicons relative to total genomic background, preventing the selection of pairs that would completely overwhelm sequencing depth with non-target containing amplicons products. If the number of non-target containing amplicons (“off-targets”) for a primer pair was 0, its C:O ratio defaults to the num_covered value, which effectively favors a "zero off-target" pair over one with noise. If no primer pairs pass this test, the selection algorithm will still select the top option out of the existing primer pairs following these ranking steps.

Within the pool of candidates passing the C:O efficiency gate, pairs were ranked first by the maximum number of target sites expected to be captured/covered (sensitivity/prioritizing highest recovery). In the event of equal coverage, the primer pair with the minimum total off-target amplicon count was prioritized to maximize sequencing efficiency (specificity). Maximum amplicon length served as the final tertiary tie-breaker, to maximize information captured. To further optimize for precision in cases where the top-ranked candidate exhibited high background noise, a secondary "Specificity Override" was implemented. If the top-ranked candidate exhibited a C:O ratio < 0.1, the algorithm performed a secondary search through the pool of potential primers for candidates within a five-target window of the maximum coverage. If a candidate within this window demonstrated a C:O ratio > 0.5, it was selected as the optimal pair, allowing for a marginal sacrifice in sensitivity to achieve a significant increase (>5-fold increase) in on-target specificity. In the event that no such candidate existed within the five-target window, the original top-ranked candidate was maintained. This selection logic prioritizes the sensitivity of the assay (capturing all desired targets) while maximizing sequencing efficiency by minimizing non-specific amplification. Final results were exported as a sorted CSV file (fullProcessed), where the primary selection criteria determined the row order. For each spacer, the selected pair was reported alongside its recovery fraction, total off-target count, and calculated C:O ratio to provide a transparent metric of assay performance. (code: processPrimers-findMotif.py).

### Calculating other RGR characteristics

The sequence distinguishability index is defined as (*SDI*) as *SDI* = (*m* – 1)/(*n* – 1), where *m* is the number of distinct target-containing amplicon sequences and *n* is the total number of covered genomic targets. *SDI* = 1 represents the case in which all amplified targets of an rgRNA in the haploid genome have at least one single-nucleotide variation (SNV) or indel beyond the spacer that distinguishes them from all the others. *SDI* = 0 represents the case in which all amplified targets have an identical sequence and are thus indistinguishable. Values of 0 ≤ *SDI* ≤ 1 represent intermediate scenarios.

The intra-chromosomal spacing score is a quantitative assessment of how well spread targets are within individual chromosomes, and is calculated as the average intra-chromosomal spacing distance. For each chromosome with more than one target, the targets were sorted by coordinate, and the distances between adjacent targets were averaged. To ensure that extremely large genomic gaps did not disproportionately inflate the score, the average distance was log-transformed. To discourage clumping, any gap between adjacent targets on one chromosome less than 500 bp was penalized by reducing the average distance for that chromosome to 1 bp. This ensured that clustered targets contributed near-zero values to the logarithmic spacing factor, effectively lowering the dispersion score for spacers hitting repetitive or localized regions. The average of the logarithmic intra-chromosomal spacing distances across all of the eligible chromosomes was then used as this score.

The inter-chromosomal entropy score uses Shannon entropy to quantify how evenly targets are distributed across different chromosomes. This rewards spacers that distribute their targets across multiple chromosomes rather than clustering them on a single scaffold.

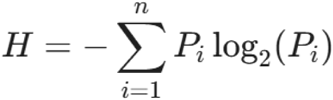

Where *n* is the number of chromosomes containing at least one target, and *P_i_* is the proportion of total targets found on chromosome *i*. With this metric, a spacer with all targets on a single chromosome results in an entropy of 0, a spacer with targets perfectly divided between two chromosomes results in an entropy of 1, and a spacer a spacer with targets perfectly divided between all n chromosomes will be log2(*n*), which for example for human (22 chromosomes + X + Y) is 4.585.

### Repeat Landscape Visualization

Repeat genome composition data and values were derived from standard RepeatMasker summary reports (the .tbl output) for mm10, hg38, and danRer11, and the pre-computed annotation files available in the bigZips directories for each assembly (e.g., hg38.fa.out.gz) from the UCSC Genome Browser database. For zebrafish (danRer11) assembly, total genome coverage was calculated based on a total assembly size of approximately 1.41 Gb. Summary statistics were extracted from the RepeatMasker Summary Table (.tbl) output. For each broad class, the percentage genome coverage was calculated as the total number of base pairs masked in the class divided by the total assembly size and multiplied by 100. The "Non-repeat" fraction was defined as the genomic remainder after subtracting the sum of all masked repeat elements from the total assembly length.

### Testing RGR predictions in HEK and N2A cells

HEK cells and N2A cells were seeded at ∼30-40% confluency on PDL-coated 24 well plates. 24 hours later, 60-80% confluent cells were co-transfected with a constitutively expressed Cas9 (lentiCRISPR v2, Addgene plasmid #52961) and an epegRNA targeting one of the selected RGRs. Each well received 300 ng Cas9 and 100 ng epegRNA. HEK cells were transfected using Lipofectamine 2000, and N2A cells were transfected using Lipofectamine 3000. Cells were passaged 24 hours post transfection, and cells were collected 3 days post transfection. RGR loci were amplified with their respective top primer pairs (**Table S4**) and sequenced with Illumina with at least 5,000 reads per sample. Amplicons were called as seen if >= 10 reads aligned to that amplicon.

### Testing mouse RGR predictions in NIH-3T3 cells stably expressing Cas9 and rgRNA

Lentivirus was produced for each LentiCRISPR v2 rgRNA construct as described. NIH-3T3 cells were seeded in a 24 well plate with ∼30,000 cells per well, and transduced 24 hours later. 48 hours post transduction, puromycin selection began. Selection media was prepared and refreshed every 48 hours. Cells were collected 72 hours later (5 days post transduction). RGR loci were amplified with their respective top primer pairs (Table S4) and sequenced on a NovaSeq X Plus instrument with at least 20,000 reads per sample.

### RGR editing quantification analysis

For each RGR, reads were aligned to the expected amplicons produced by the primer pair used to amplify them. Sequencing data, containing read counts and percent editing metrics per reference amplicon, were exported as wide-format CSV files. Data processing was performed using a custom Python script utilizing the pandas library. Sample metadata (i.e. experimental condition. biological replicate, etc) were parsed directly from column headers. To ensure robust analysis, amplicons with fewer than 1,000 total sequencing reads summed across all samples were excluded. To account for variations in sequencing depth across different libraries, raw read counts were normalized by calculating the count fraction for each amplicon, defined as the read count at a specific location divided by the total read count for that specific sample replicate.

Percent editing and count fractions were averaged across biological replicates to calculate the mean and standard error of the mean (SEM) for both the Cas9-H1 and NC conditions. Reference amplicons were then sorted in descending order based on their mean percent editing in the Cas9-H1 condition. To systematically distinguish true editing from background sequencing noise, a heuristic threshold was applied: an amplicon was classified as "significantly edited" if its mean Cas9-H1 percent editing was strictly greater than the background noise limit, defined as the mean NC percent editing plus one SEM of the NC condition. For each RGR, summary statistics were calculated based on this threshold. The total number of edited amplicons was counted as the number of unique reference amplicons exceeding the threshold. The total number of edited sites was calculated by summing the number of locations for all edited amplicons. These values were then divided by the total number of evaluated reference amplicons and the total predicted locations (sites), respectively, to calculate the overarching fraction of edited locations and the fraction of edited sites per target ID.

### AAV production, purification, and quantification

All AAVs were produced in house. 50 mM chloroquine (Cell Signaling Technology 14774) was prepared from 150 mg powder in 5.82 mL of ultra pure water, and filter sterilized through a 0.22 micron syringe filter. 1 mg/mL pH 7.0 PEI Prime™ linear polyethylenimine (Sigma-Aldrich 919012) was prepared in 100 mL of ultra pure water from 100 mg powder, and filter sterilized through a 0.22 micron syringe filter.

AAVs were generated in HEK cells and purified using an adapted iodixanol gradient purification protocol. HEK cells were seeded in 150 mm dishes at 30-40% confluency, to be at 60-70% confluency 24 hours later. Cell media was refreshed 30-60 minutes before transfection with chloroquine media. Chloroquine media was prepared by adding 22 µL of the 50 mM stock to 44 mL of media. AAVs were packaged using the AAV-PHP.eB capsid to efficiently transduce the central nervous systems (pAdDeltaF6, Addgene #112867 and pUC-mini-iCAP-PHP.eB, Addgene #103005). The ratio of cargo AAV plasmid to packaging plasmids for transfection was: 1:2:1 AAV:pUCmini:pAdDelta. PEI was used for transfection at a 3:1 ratio of PEI to DNA. Media was refreshed 8-12 hours post transfection. Cells and viral supernatant were collected 96 hours later, and AAV was harvested and purified. Genomic AAV titer was determined first by qPCR, and verified by imaging sagittal brain sections and quantifying mCherry expression of DAPI stained nuclei.

### Experimental model and subject details for recording Fos/Npas4 in mice

3-4 week old Cas9 male mice were used at the beginning of all experiments. AAV was delivered systemically via retroorbital injections. 50 uL (∼1E10 GC) were injected per mouse, and treated with 50 mg/mL Proparacaine Hydrochloride at the injection site. 50 mg/mL Proparacaine Hydrochloride (Millipore Sigma PHR3229) was prepared from 500 mg powder in 10 mL PBS, aliquoted, and stored at -20 C. Mice were observed daily for 5-7 days following injection. Animals were sacrificed 3 and 10 weeks later.

### Kainic acid induced *status epilepticus*

Mice (24-29 g) were placed on a dox diet 2 days post AAV delivery for 2 weeks to keep the recorder off. 3 weeks post AAV delivery (one week after mice were switched off of the dox diet). Mice were randomly assigned to two groups, (1) Kainic acid induced status epilepticus (SE), and (2) Healthy controls.

Kainic acid (Sigma Aldrich, 58002-62-3) was dissolved in sterile saline (2.5 mg/mL KA at up to 2.5 mg/kg). Mice were initially weighed to determine the volume of kainic acid required for the dose, and kainic acid was injected intraperitoneally (IP) at a dose range of 2.5-10 mg/kg until SE was developed^103^. The onset of SE was defined as the point at which the mice exhibited at least unceasing stage 4 or higher for 5 minutes or more on the modified Racine scale^104,105^, as outlined below:

- Stage 0: Normal activity
- Stage 1: Freezing and slight head nodding, arched body
- Stage 2: Continuous head bobbing, wet dog shakes, straub tail
- Stage 3: Partial myoclonus, occasional whole-body jerks, continuous body tremors, intensified freezing, extended/arched body with jerks
- Stage 4: Increased immobility and freezing, uncontrolled circling movement, continuous whole-body jerks lasting >5 minutes
- Stage 5: Continuous straub tail, loss of limb control followed by generalized tonic–clonic seizures, or one episode of rearing),
- Stage 6: Loss of balance, more than one episode of rearing followed by occasional falling, jumping and rolling over, generalized tonic extension of the body, cardiopulmonary collapse, and death

These seizures typically originate in the limbic system; however, they rapidly generalize, resulting in high-amplitude electrical discharges across the cortical regions^103^. Animals that did not develop SE were not included in the study.

All animals that experienced SE received 200-600 µL fluids (LRS) subcutaneously to replenish fluids 15-20 minutes post-KA administration. Mice were continuously monitored for 3 hours post KA injection, then returned to their home cages with liquid/gel food and water (Clear H2O Hydrogel, hydration supplement). They received a second injection of fluids/saline ∼4-6 hours later with daily checks. The animals were euthanized, and brain regions were harvested 4 weeks post SE-induction.

### Nuclei dissociation and staining

Brain and liver tissues from injected Cas9 mice were harvested. Then brain regions of interest (cerebral cortex, hippocampus, thalamus, and cerebellum) were dissected and flashfrozen using liquid nitrogen. Tissue was transferred to a Dounce homogenizer containing ice-cold lysis buffer (10 mM Tris-HCl pH 7.4, 10mM NaCl, 3mM MgCl2, and 0.1% Nonidet P40) and homogenized with 10 strokes of pestle A and 10 strokes of pestle B. The homogenate was incubated on ice for 5 minutes, further dissociated by pipetting 10 times using a wide-bore 1000uL pipette tip, and then filtered through a 40uM filter. The filter was washed with 1% BSA in PBS. Samples were centrifuged at 500xg for 5 minutes at 4C and then the pellet was resuspended in 1% BSA in PBS.

To remove debris, a sucrose cushion solution (1.8M sucrose, 10mM Tris-HCl pH 8.0, 3mM MgCl2, and 1mM DTT) was added to the bottom of a 2mL microcentrifuge tube. The nuclei suspension was mixed with the sucrose cushion solution for a final concentration of (1.16M sucrose, 6.43mM Tris-HCl ph 8.0, 1.93mM MgCl2, and 0.643mM DTT). This nuclei suspension and sucrose mixture was then carefully layered over the sucrose cushion layer, and centrifuged at 13,000xg for 45 minutes at 4C. The debris layer was carefully removed and pellets were washed 1x with 1% BSA in PBS.

Liver samples were incubated in blocking buffer (3% BSA Solution in PBS with 0.05% TritonX-100) for 30 minutes at room temperature with gentle rotation. Rat anti-mCherry antibody (ThermoFisher Scientific, M11217) was added at 1:500 and incubated for 1 hour at room temperature with gentle rotation. Samples were washed 3 times with 1% BSA in PBS (500xg for 3 minutes at 4C). Pellets were resuspended in blocking buffer and anti-rat Alexa Fluor 594–conjugated secondary antibody (ThermoFisher Scientific, A-21209) was added at 1:500 and incubated for 30 minutes at room temperature with gentle rotation, protected from light. Samples were washed 3x with 1% BSA in PBS and then resuspended in 1% BSA in PBS.

For brain samples, native mCherry fluorescence was used. Following debris removal, samples were resuspended with 1% BSA in PBS. Anti-NeuN antibody conjugated to Alexa Fluor 647 (Abcam, AB190565) was added at 1:1000 and incubated for 1 hour on ice, protected from light. Then samples were washed once with 1% BSA in PBS and resuspended in 1% BSA in PBS.

DAPI was added 10 minutes prior to sorting and samples were filtered through a 40uM filter immediately before sorting.

### Flow sorting for mCherry+ cells and sequencing of RGR loci

All flow cytometry experiments were performed using a Sony MA900 system with 100-uM sorting chips (Sony). Prior to each sorting session, the instrument was calibrated using the targeted setting with Auto Setup Beads (Sony) after ethanol priming. Gates were used to remove debris and doublets before sorting for cells. Briefly, an initial gate based on forward scatter area (FSC-A) versus backscatter area (BSC-A) was used to exclude debris (events with high FSC-A and/or BSC-A signal) (**Figure S7b**). Nuclei were identified using FSC-A vs DAPI-A to select for events with high DAPI signal. Doublets were removed using FSC-A vs FSC-H and FSC-A vs FSC-W gating (**Figure S7b**). For brain samples, cells were gated for NeuN positive signal (Alexa Fluor 647) and mCherry expression, and 100 NeuN⁺/mCherry⁺ cells were sorted per replicate per brain region (**Figure S7c**). For liver samples, 100 mCherry⁺ cells were sorted per replicate (**Figure S7c**).

Cells were sorted directly into 4 µL of QuickExtract (Lucigen QE09050). As described in the post sort steps for library preparation of the LTR5-RGR1 single cell sequencing library, DNA was extracted by incubating the 96-well plates for 10 minutes at 65°C followed by 5 minutes at 98°C to inactivate the QuickExtract. PCR1 and PCR2 were performed as described in that section as well, with SBS3 and SBS9 adaptors added to the amplicons in PCR1 and indexing pairs and Illumina sequencing adaptors were added in PCR2. Following these steps for library preparation, MMERVK10C-int-RGR1 was amplified using its corresponding primers (Table S4) and sequenced on a NovaSeq X Plus instrument with at least 50,000 reads per 100 cell sample.

### Calculating relative Fos expression following kainic acid induced seizures in the cerebellum versus the hippocampus

The area under the curve (AUC) of AP-1 binding over time in the hippocampus and cerebellum was calculated from published data^89^, and the ratio of the AUC in hippocampus to AUC in cerebellum was measured. AP-1 AUC in the cerebellum was found to be ≥25-fold lower than hippocampus in the hours after KA injection.

### Statistical analysis of kainic acid induced seizure data

All statistical analyses and data visualizations were performed using R software. Boxplots were generated using the ggplot2 and ggpubr packages. For all pairwise comparisons between the KA+ and KA- conditions, the negative control (NC, no AAV) group was excluded from the statistical testing. Statistical significance was defined as p < 0.05. Differences in editing percentages between the KA+ and KA- conditions were evaluated independently for each brain region. Due to the non-normal distribution of the data, as well as the a priori hypothesis that the KA+ condition increases editing rates, an unpaired one-sided Wilcoxon rank-sum test (Mann-Whitney U test) was performed for each region. The alternative hypothesis tested was that editing percentages in the KA+ group were strictly greater than those in the KA- group. All data points within a condition were treated as independent observations for these pairwise comparisons.

To evaluate global shifts in editing efficiency between KA+ and KA- conditions, statistical comparisons were performed on the per amplicon editing observed within each isolated brain region. Raw sequencing counts from technical and biological replicates were first aggregated to calculate a single mean percent edited value for both the KA+ and KA- conditions at every targeted genomic location. To determine if the KA+ condition induced a significant global change in editing, a Paired Wilcoxon Signed-Rank Test was employed. This specific statistical framework was selected intentionally because it is both paired and non parametric. Locus-specific pairing was needed because editing efficiency is heavily dependent on locus-specific factors, such as local chromatin accessibility and sequence-specific guide RNA binding dynamics (i.e. because of mismatches between target sequence and gRNA). By treating the genomic location as the pairing variable, each amplicon serves as its own internal control. This isolates the effect of the KA condition from the inherent baseline variability between different loci. It was important to have a non-parametric data structure because the distribution of editing efficiencies was both highly skewed and zero-inflated, violating the assumption of normality required for parametric tests (e.g., the paired t-test). The Wilcoxon Signed-Rank Test relies on rank differences rather than raw means, making it robust to these extreme distributions and outliers. By evaluating the paired differences at each targeted amplicon, the Wilcoxon Signed-Rank Test allowed for the evaluation of systematic, landscape-wide shifts in editing efficiency associated with the KA+ environment. All statistical analyses were performed in R, utilizing standard corrections for exact zero-ties.

### Statistical analysis of single neuronal nuclei

To compare the distributions of neuronal editing between the mice that received kainic acid induced seizures vs the mouse that did not, we employed non-parametric statistical methods suitable for single-cell datasets with potential skewness and heterogeneity. Statistical significance was determined using the two-sample Kolmogorov-Smirnov (K-S) test, which assesses differences in both the location and the shape of the empirical cumulative distribution functions (ECDF). Unlike tests of central tendency, the K-S test is sensitive to differences in the tails of the distribution, making it appropriate for detecting sub-populations of high-activity neurons, given our model where only a sub-population of neurons respond to seizures, and the highest responders likely die immediately. Quantile-quantile (Q-Q) plots were generated to visualize the relationship between the editing distributions of the two groups by plotting their respective quantiles against each other. Empirical cumulative distribution function (ECDF) plots were utilized to visualize the maximum vertical distance (*D* statistic) between the groups. All analyses and visualizations were performed in R using the ggplot2 and ComplexHeatmap packages.

### Ethical compliance

We have complied with all relevant ethical regulations for animal testing and research. The study protocol was approved by the Johns Hopkins University Animal Care and Use Committee (ACUC).

## Acknowledgements

The authors would like to acknowledge Dr. Theresa Loveless and Dr. Keith Dveirin for comments on the manuscript, Dr. Jesse Handler, Jinwoo Jun, James Forsmo, Dr. Weixiang Fang, Dr. Dustin Shigaki, and Dr. Nikhil Hajirnis for feedback on the project. This work was supported by the National Institutes of Health (NIH) (R01HG012357, R.K.), and the David & Lucile Packard Foundation (2020–71380, R.K.). R.K.D and J.D.L. were supported by NSF Graduate Research Fellowships. Parts of this work were carried out at the Advanced Research Computing at Hopkins (ARCH) core facility, which is supported by the National Science Foundation (NSF) grant number OAC 1920103.

## Author Contributions

R.K.D. and R.K. conceived the study and designed experiments; R.K.D. carried out experiments and analyzed data. J.D.L. assisted experimental design, single-cell and single-nucleus cell sorting, dissections, and AAV injections. P.V. and S.K. helped design and implement neuronal recording in the epilepsy model. Y.Y. assisted in AAV production. J.L. conducted xenograft experiments. J.J.L. and S.K.R. performed RGR tests in NIH/3T3. X.D. provided access and expertise in cell sorting. B.L. provided sequence analysis expertise. B.L. and S.K.R. provided critical feedback throughout the course of the project. R.K.D. and R.K. interpreted the data and wrote the manuscript with input from B.L. and S.K.R. R.K. supervised the project.

## Data visualization and schematics

Heatmaps were generated in R using ComplexHeatmap^106^ 2.20.0. Unless otherwise noted, all other plots were either generated in R using ggplot2^107^ 3.5.1 or Adobe Illustrator. Schematics were created using Adobe Illustrator.

## Data Accessibility

All sequencing data generated in this study have been submitted to the National Center for Biotechnology Information (NCBI) Sequence Read Archive (SRA) under project ID PRJNA1414787. All other raw data is available upon reasonable request.

## Supplementary Figures

**Figure S1:**
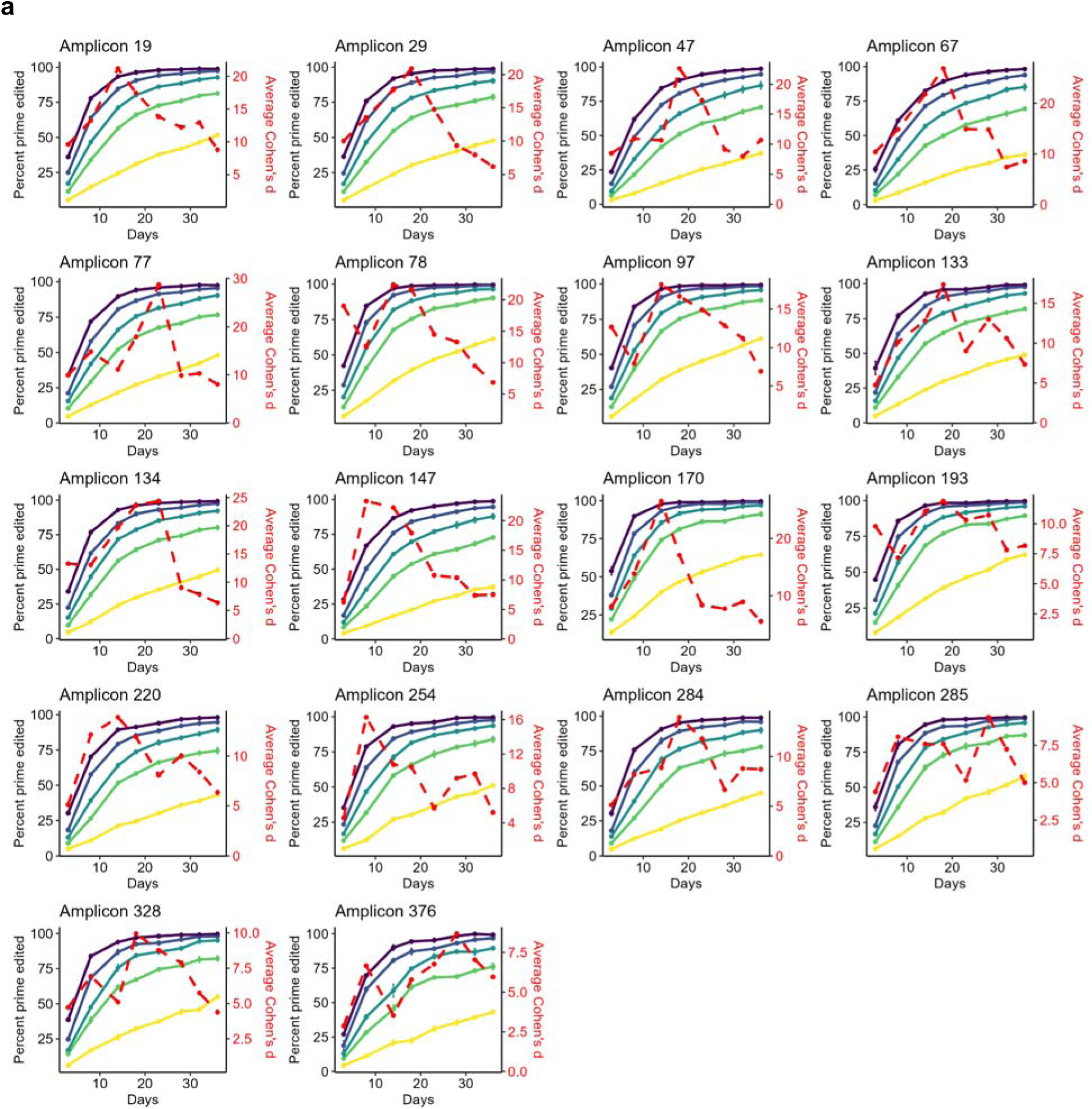
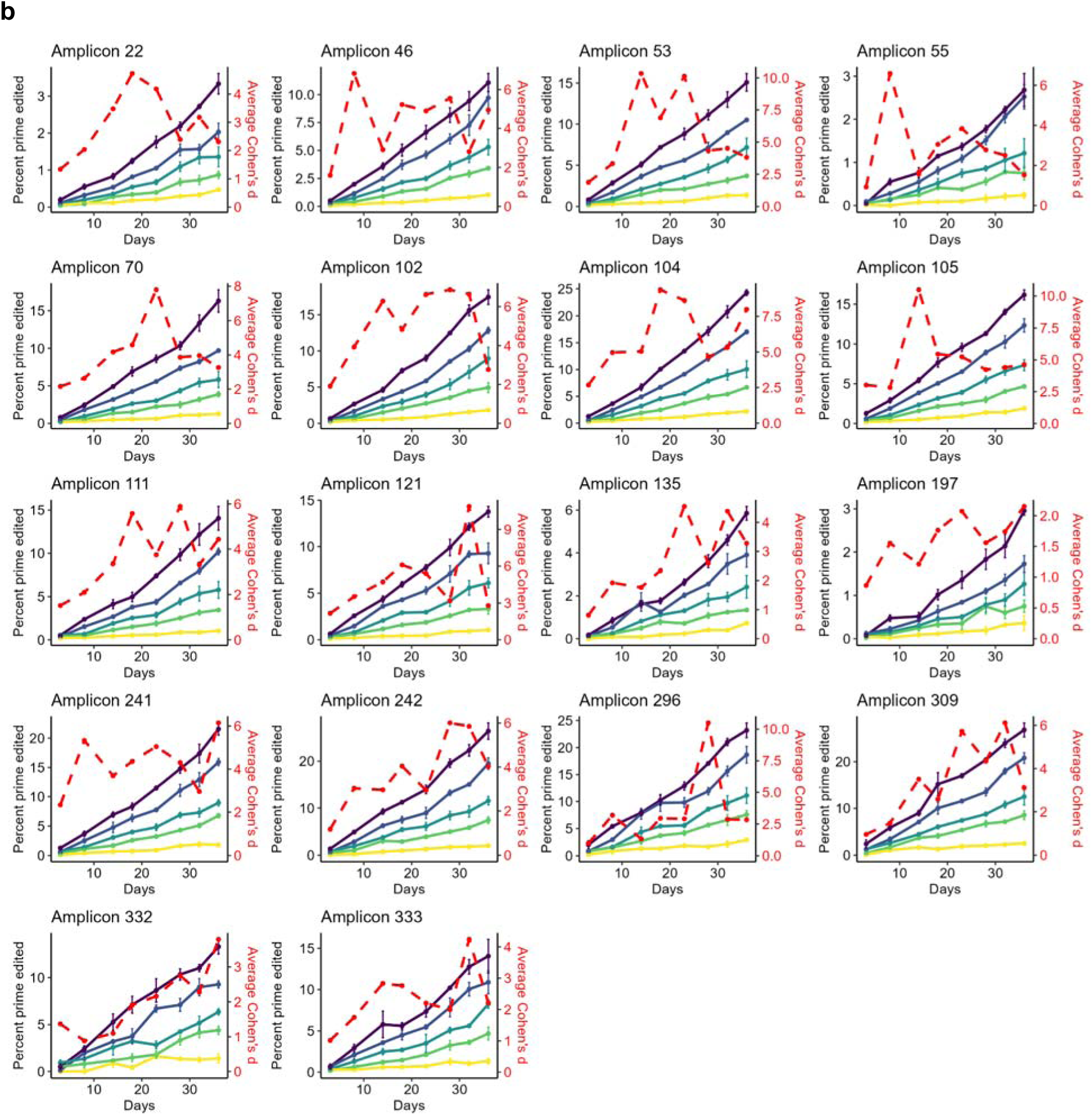
Diverse editing kinetics of LTR5-RGR1 amplicons. **a**, Representative fast-editing amplicons. Individual plots show prime editing accumulation over time for amplicons with rapid editing kinetics. Left y-axis (solid lines): percent prime edited at each Shield1 concentration (color-coded by treatment condition as in Figure 3a). Right y-axis (red dashed line): average Cohen’s *d* across all pairwise treatment comparisons at each time point, reflecting discriminatory power between conditions. Fast-editing amplicons reach high editing levels early but saturate across conditions with prolonged exposure, resulting in declining Cohen’s d values over time. **b**, Representative slow-editing amplicons. Individual plots show prime editing accumulation over time for amplicons with slow editing kinetics. Axes as in **a**. Slow-editing amplicons accumulate edits gradually and do not saturate, maintaining or increasing discriminatory power (Cohen’s *d*) over extended exposure times. However, these amplicons lack sufficient separation between conditions at early time points.

**Figure S2:**
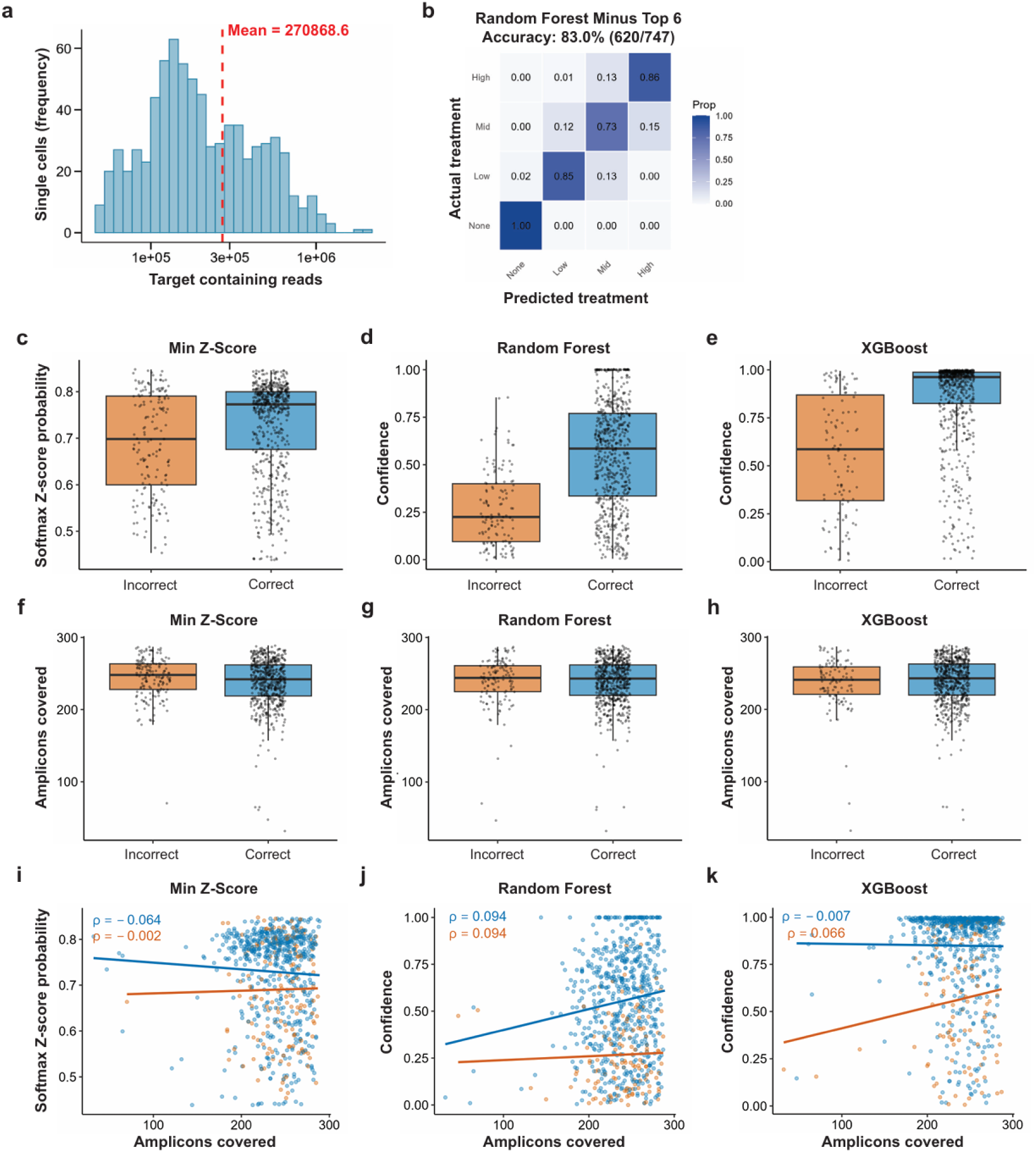
Single-cell classification performance and quality metrics across algorithmic frameworks. **a**, Distribution of target-containing sequencing reads per single cell. Red dashed line indicates mean coverage (270,868.6 reads). **b**, Confusion matrix for single-cell classification using Random Forest after excluding the six most informative amplicons. Rows indicate actual treatment group; columns indicate predicted treatment group. Cells are colored by the proportion of cells in each category. Overall accuracy: 83.0% (620/747 cells). **c-e**, Classification confidence for correctly versus incorrectly classified single cells. Box plots show the distribution of confidence scores for cells classified correctly (blue) or incorrectly (orange). For minimum Z-score (**c**), confidence was calculated as the softmax probability of the predicted class. For Random Forest (**d**) and XGBoost (**e**), confidence was calculated as the difference between the highest and second-highest class probabilities. (**c**) Minimum Z-score: mean 0.729 (correct) vs. 0.690 (incorrect), *P* = 5.31 × 10⁻⁵. (**d**) Random Forest: mean 0.554 vs. 0.268, *P* = 6.44 × 10⁻³³. (**e**) XGBoost: mean 0.865 vs. 0.562, *P* = 6.70 × 10⁻¹⁶. *P* values from two-sided Student’s t-test. **f-h**, Sequencing coverage for correctly versus incorrectly classified single cells. Box plots show the number of amplicons covered per cell for cells classified correctly (blue) or incorrectly (orange) using (**f**) minimum Z-score (*P* = 0.013), (**g**) Random Forest (*P* = 0.94), and (**h**) XGBoost (*P* = 0.42). *P* values from two-sided Student’s t-test. **i-k**, Relationship between amplicon coverage and classification confidence. Scatter plots show the number of amplicons covered (x-axis) versus model prediction confidence (y-axis) for correctly (blue) and incorrectly (orange) classified cells using (**i**) minimum Z-score, (**j**) Random Forest, and (**k**) XGBoost. Solid lines represent linear regression fits. Spearman’s ρ is annotated for each group.

**Figure S3:**
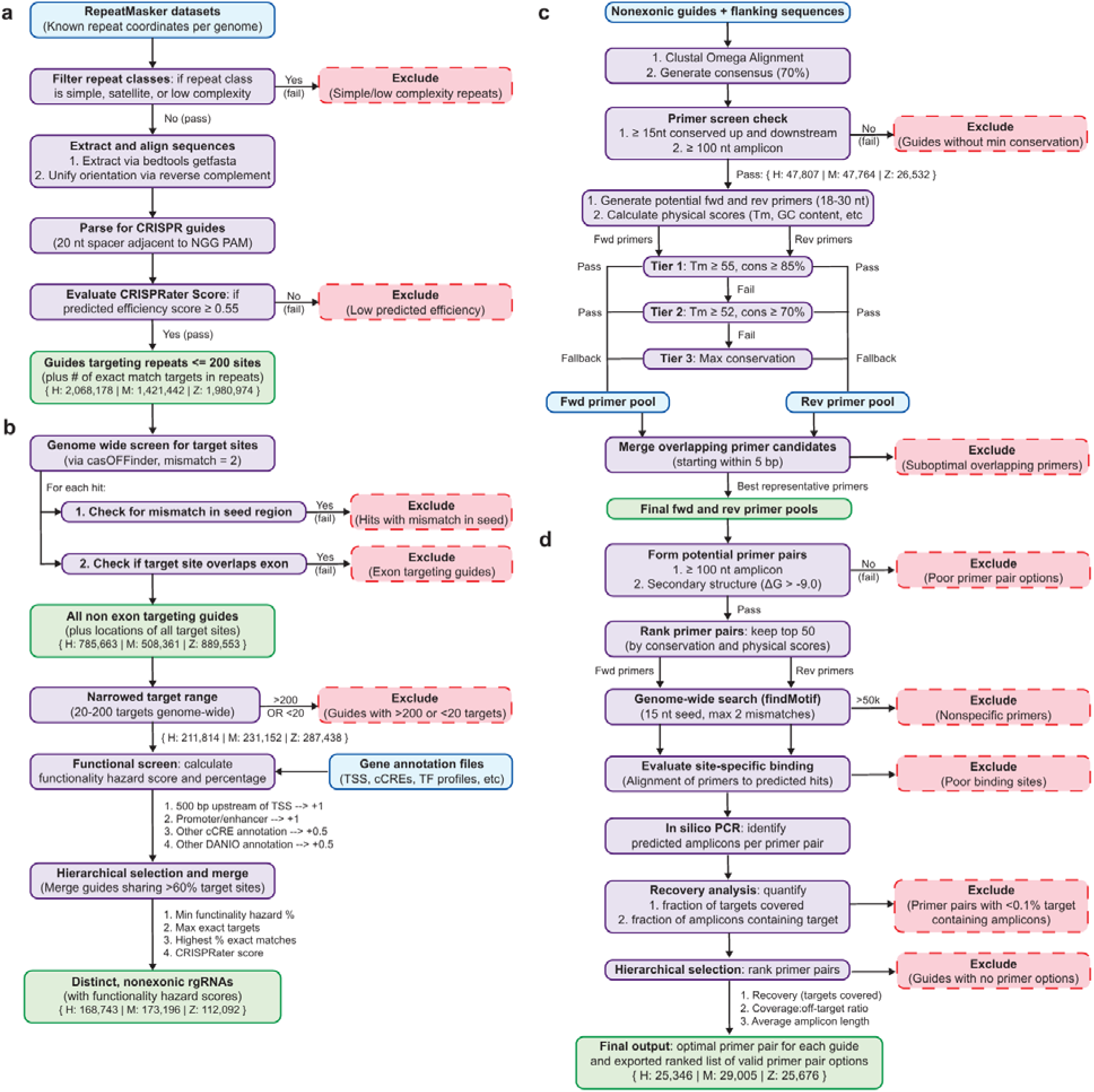
Computational pipeline for RGR identification. **a,** Guide RNA identification from repetitive elements (Part 1). RepeatMasker annotations are filtered to exclude simple and low-complexity repeats. Using the remaining RepeatMasker coordinates, sequences are extracted from reference genome, aligned within each repeat element, and parsed for CRISPR-compatible spacers (20 nt adjacent to NGG PAM). Guides with up to 200 targets are evaluated using CRISPRater, and those with predicted efficiency scores ≥0.55 are retained. Numbers indicate guides targeting repeats identified per genome (H: human; M: mouse; Z: zebrafish). **b,** Genome-wide target site identification and functional screening (Part 2). Target sites are identified using casOFFinder (allowing up to 2 mismatches). Hits with seed-region mismatches are excluded. Guides with targets overlapping or within 20 nt of an exon are excluded. Guides are filtered to retain those with 20–200 target sites and subsequently screened for potential functional disruption based on proximity to transcription start sites (TSS), promoters, enhancers, and cis-regulatory elements (cCREs). Guides sharing >60% of target sites are merged hierarchically, yielding distinct, non-exonic rgRNAs with associated functionality hazard scores. **c,** Primer pool generation (Part 3). Flanking sequences for each guide are aligned (Clustal Omega) to generate a consensus (70% threshold). Guides with ≥15 nt conserved sequence upstream and downstream with ≥100 nt amplicon length between them are retained. Forward and reverse primers (18–30 nt) are generated and filtered through a tiered system based on conservation across flanking sequences and physical properties like melting temperature (Tm) and nucleotide composition. Overlapping primer candidates (starting within 5 bp) are merged to a superior candidate, yielding final forward and reverse primer pools. **d,** Primer pair optimization and recovery analysis (Part 4). Potential primer pairs are formed and filtered based on amplicon length (≥100 nt) and secondary structure (ΔG > −9.0); pairs not meeting these are filtered out. Top 50 pairs are ranked by conservation and physical scores. Genome-wide binding specificity is evaluated using findMotif (15 nt seed, ≤2 mismatches), and pairs with >50,000 predicted hits or poor site-specific binding are excluded. In silico PCR identifies predicted amplicons per primer pair. Recovery analysis quantifies the fraction of targets covered and fraction of amplicons containing targets. Primer pairs with <0.1% target-containing amplicons are excluded. Final output: optimal primer pair for each guide with a ranked list of valid alternatives. Final RGR counts: H: 25,346; M: 29,005; Z: 25,676.

**Figure S4:**
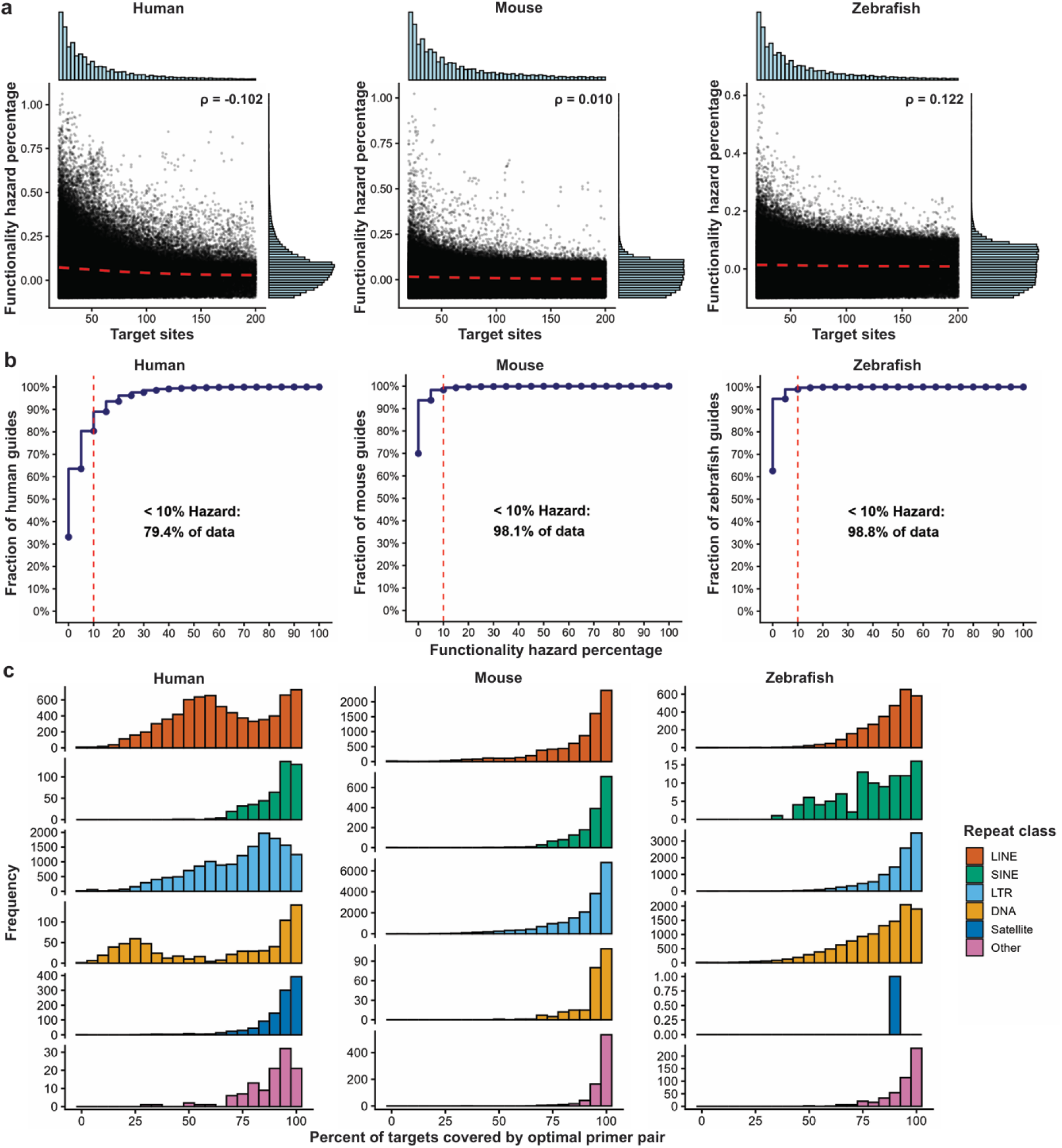
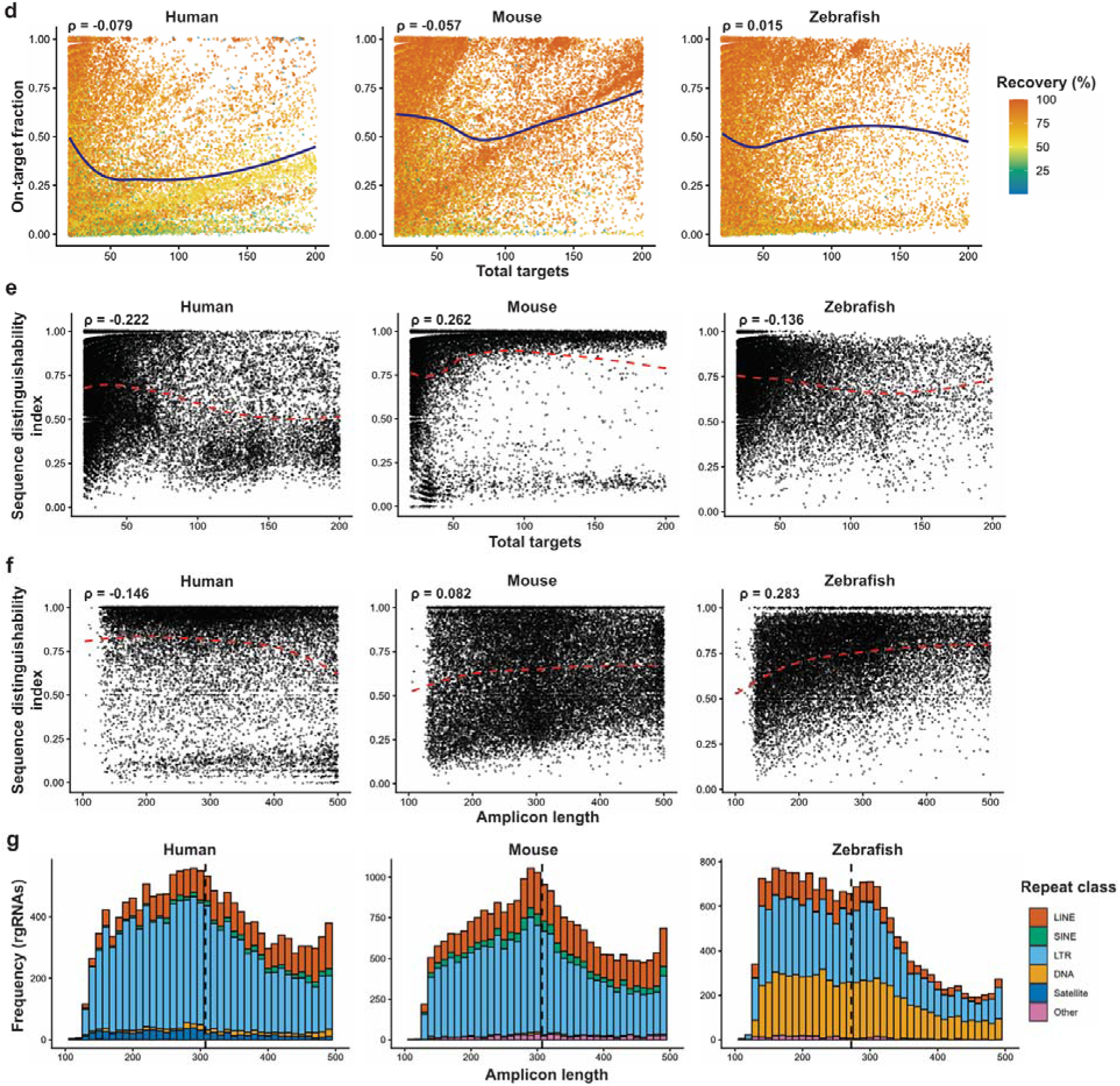
Comparative analysis of RGR properties across human, mouse, and zebrafish genomes. **a**, Relationship between target site count and functionality hazard percentage for non-exonic rgRNAs with 20–200 targets. Scatter plots display each rgRNA’s total target count (x-axis) versus functionality hazard percentage (y-axis). Marginal histograms show the frequency distributions of both variables. Red dashed lines represent LOESS regression fits. Spearman’s ρ is indicated for each species. **b**, Cumulative distribution of functionality hazard percentage across rgRNAs. Plots show the fraction of rgRNAs (y-axis) with functionality hazard percentage at or below a given threshold (x-axis). Red dashed vertical line indicates the 10% hazard threshold. Annotations indicate the percentage of rgRNAs with <10% functionality hazard (human: 79.4%; mouse: 98.1%; zebrafish: 98.8%). **c**, Distribution of optimal primer pair coverage stratified by repeat class. Histograms show the frequency of rgRNAs across primer coverage bins for each repeat class (LINE, SINE, LTR, DNA, Satellite, Other) in human, mouse, and zebrafish. **d**, Relationship between target site count and on-target fraction. Scatter plots show total target count (x-axis) versus on-target fraction (y-axis) for each rgRNA with its optimal primer pair. Points are colored by recovery percentage. Blue dashed lines represent LOESS regression fits. Spearman’s ρ is indicated for each species. **e**, Relationship between target site count and sequence distinguishability index. Scatter plots show total target count (x-axis) versus sequence distinguishability index (y-axis) for each rgRNA. Red dashed lines represent LOESS regression fits. Spearman’s ρ is indicated for each species. **f**, Relationship between amplicon length and sequence distinguishability index. Scatter plots show amplicon length (x-axis) versus sequence distinguishability index (y-axis) for each rgRNA. Red dashed lines represent LOESS regression fits. Spearman’s ρ is indicated for each species. **g**, Distribution of average amplicon lengths stratified by repeat class. Stacked histograms show the frequency of rgRNAs across amplicon length bins, colored by repeat class (LINE, SINE, LTR, DNA, Satellite, Other). Black dashed lines indicate the mean amplicon length for each species.

**Figure S5:**
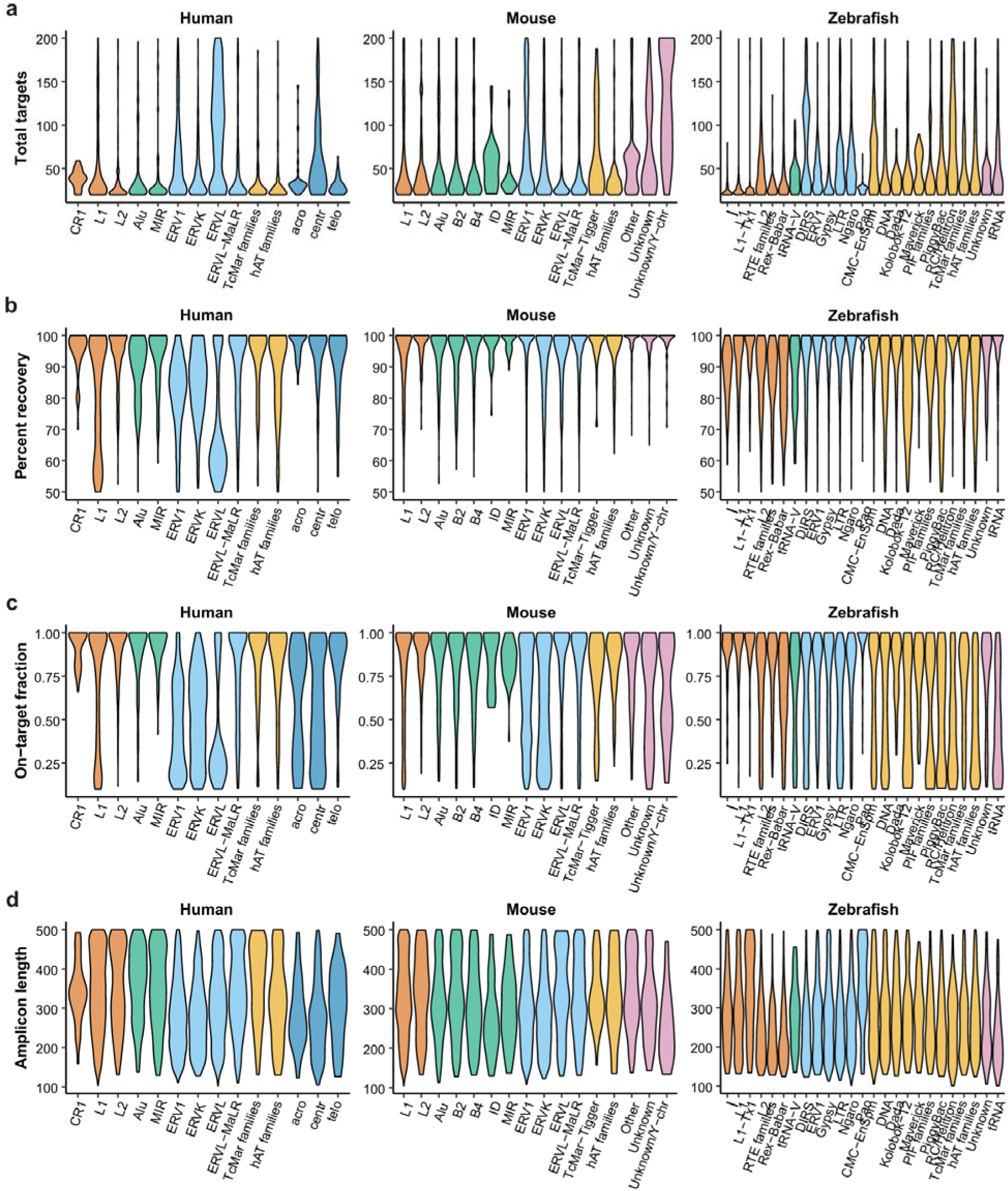
RGR properties per repeat family across human, mouse, and zebrafish genomes. **a**, Distribution of total target sites across repeat families. Violin plots show the distribution of target site count (y-axis) for rgRNAs within individual repeat families (x-axis) in human, mouse, and zebrafish. Violins are colored by repeat class following the same color scheme as figure 6 (LINE = red-orange, SINE = green, LTR = light blue, DNA = yellow-orange, Satellite = dark blue, Other = pink, in listed order). **b**, Distribution of primer recovery across repeat families. Violin plots show the distribution of percent recovery (y-axis), defined as the fraction of target sites expected to be covered by the optimal primer pair, for rgRNAs within individual repeat families (x-axis), colored by repeat class. Repeat families are ordered along the x-axis within each species by average total targets. **c**, Distribution of on-target fraction across repeat families. Violin plots show the distribution of on-target fraction (y-axis), defined as the proportion of predicted amplicons that contain a target site, for rgRNAs within individual repeat families (x-axis), colored by repeat class. Higher values indicate greater primer specificity for target-containing sequences. **d**, Distribution of amplicon length across repeat families. Violin plots show the distribution of amplicon length (y-axis) for rgRNAs within individual repeat families (x-axis), colored by repeat class. **a-d**, Repeat families are ordered along the x-axis within each class alphabetically.

**Figure S6:**
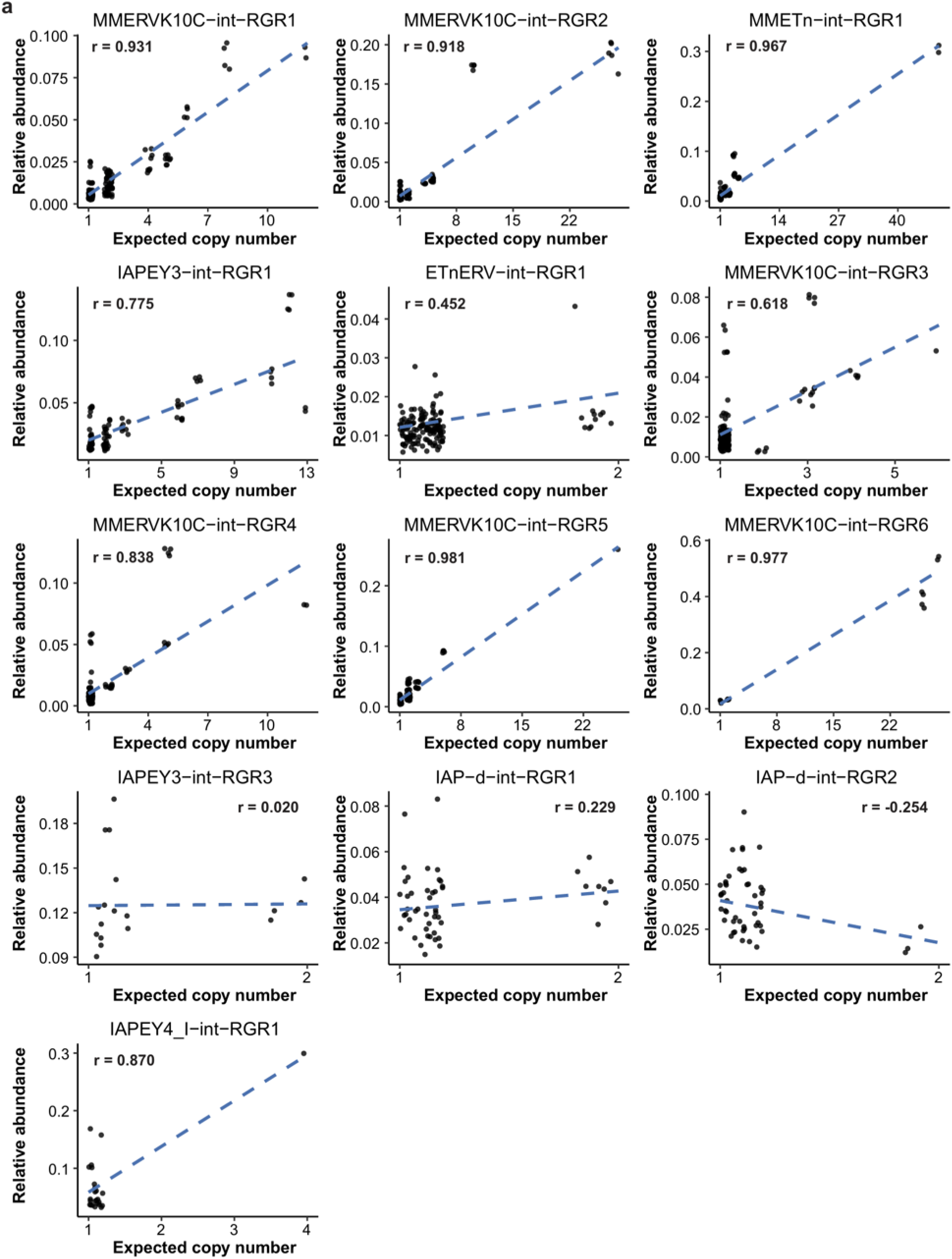
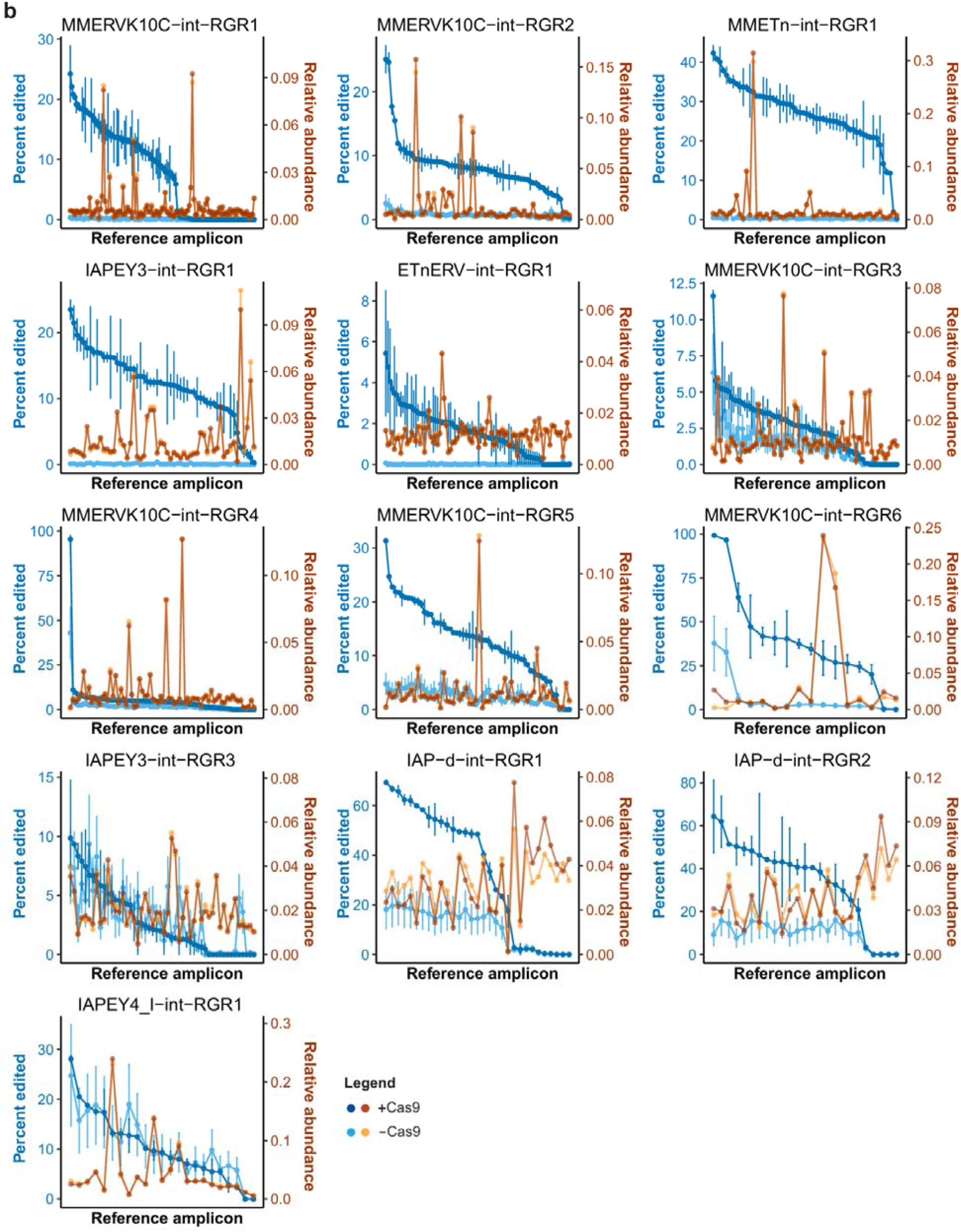
Distribution of editing and amplicon recovery across tested RGRs. **a**, Correlation between predicted haploid copy number and sequencing abundance for distinguishable amplicons targeted by mouse rgRNAs in stable cell lines. Scatter plots show the relationship between predicted copy number (x-axis, jittered by 0.2) and relative sequencing abundance (y-axis) for each distinct amplicon sequence. Each data point represents one amplicon in one sample (up to 4 samples per amplicon: 2 +Cas9, 2 −Cas9 controls). Pearson’s r is indicated for each rgRNA. Data correspond to experiments shown in Figure 7d,e. **b**, Editing efficiency and amplicon recovery across individual RGR target amplicons. Line plots show, for each rgRNA, the editing efficiency (blue lines; left y-axis: percentage of reads aligned to each reference amplicon that are edited) and relative amplicon abundance (orange lines; right y-axis: proportion of total reads aligned to each reference amplicon). +Cas9 samples (dark blue, dark orange) and −Cas9 negative controls (light blue, light orange) are shown. Amplicons are ordered by decreasing editing efficiency along the x-axis. Data points represent mean ± s.e.m.

**Figure S7:**
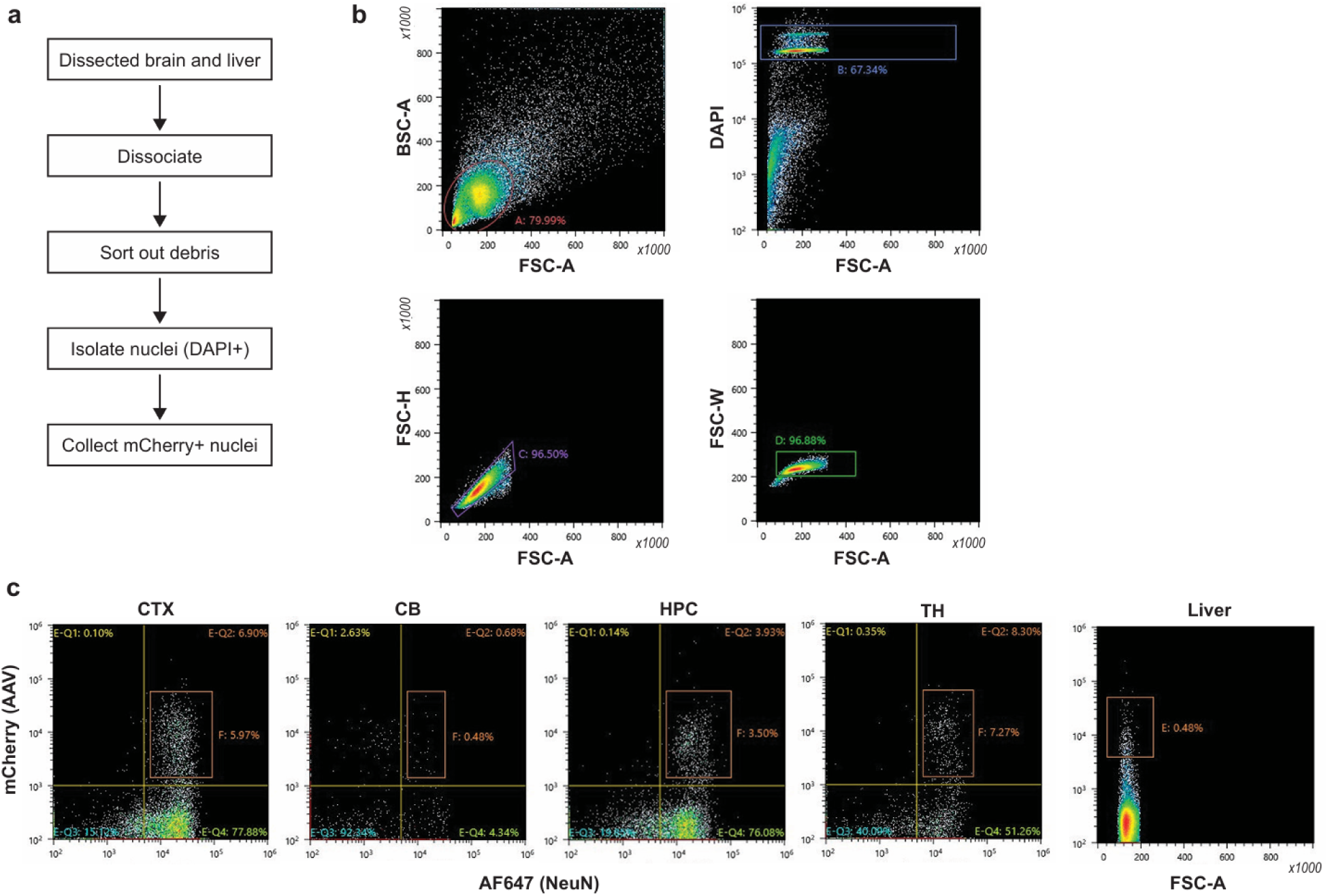
Fluorescence assisted sorting of neuronal and liver nuclei. **a**, Schematic. Mouse liver and brains were dissected, dissociated, and sorted via FACS. Gates were applied to remove cellular debris before gating for mCherry positive (AAV) and AlexaFluor647 (AF647) positive (NeuN) cells (except liver, mCherry only). **b**, Representative FACS plots from cortex to demonstrate isolation of nuclei and sorting out debris. **c**, Plots of the cell populations that were collected for all five tissues. NeuN⁺/mCherry⁺ population collected for all four brain regions (the four leftmost plots, mCherry as a measure of AAV expression plotted against AlexaFluor647 as a measure of NeuN expression), mCherry⁺ population for liver (far right plot, mCherry as a measure of AAV expression plotted against FSC-A).

## Supplementary Tables

**Table S1. Genome-wide LTR5-RGR1 guide target sites and amplicon coverage**. This table lists all 609 LTR5-RGR1 guide target sites identified across the human genome (hg38), followed by the 33 non-target containing amplicons predicted to be amplified by the LTR5-RGR1 sequencing primers used. Index: arbitrary row identifier (1–609 for target sites; OT1–OT33 for off-target amplicons). Chromosome: chromosome location. TargetSeq (+ PAM): 20-nucleotide target sequence plus 3-nucleotide PAM; lowercase letters indicate mismatches relative to the guide sequence; "NA" for off-target amplicons. Strand: genomic strand (+ or −). Mismatches: number of mismatches between the guide and target sequence (0, 1, or 2); "NA" for off-targets. Start Coord / End Coord: genomic coordinates of the target sequence (for target sites) or the predicted amplicon (for off-targets). Covered: whether the site is predicted to be amplified by sequencing primers. Amplicon Length: predicted amplicon size in base pairs; "NA" if not covered. Fwd/Rev Primer Binding Score: predicted binding score for forward and reverse sequencing primers; "NA" if not covered. PE Score: conservation score (0–29) for the 29 nucleotides downstream of the PAM; "NA" for off-targets. Prime Editable (with common pegRNA): "Yes" if PE Score > 27, indicating compatibility with a common pegRNA design; "NA" for off-targets. Unique: "Yes" if the amplicon contains distinguishing sequence variants allowing unique identification in amplicon sequencing; "No" if indistinguishable from other amplicons; −1 if not covered or off-target. Functionality: genomic annotation of the target site (Exonic, Intronic, or None/Intergenic); "NA" for off-targets.

**Table S2. LTR5-RGR1 reference amplicon sequences and estimated copy numbers.** This table lists the 387 reference amplicons identified from bulk amplicon sequencing of negative control samples in the selected monoclonal iiPE2 + LTR5-RGR1 pegRNA cell line (see **Methods**). Reference amplicon ID: unique identifier (1–387). Sequence: amplicon sequence with primer sequences trimmed. Abundance: total read count aligned to each reference amplicon across negative control samples used to build the reference list. Target: the 20-nucleotide sequence corresponding to the guide target site within the amplicon. Guide alignment score: pairwise alignment score between the 20-nucleotide target sequence and the LTR5-RGR1 guide sequence. PAM: the 3-nucleotide protospacer adjacent motif immediately downstream of the target sequence. On-target: "on" if the amplicon contains a functional CRISPR target site (≤2 mismatches with the guide sequence outside the 6-nucleotide seed region and PAM ending in "GG"); "off" otherwise. Average relative abundance: average fraction of reads aligning to each reference amplicon across negative control samples. Sequence length: length of the trimmed amplicon sequence in nucleotides. Estimated copy number: predicted number of genomic loci producing each amplicon sequence in the haploid genome, estimated from single-cell amplicon sequencing data in the D7 clonal line (see **Methods**).

**Table S3. Summary of repeat-targeting guide RNA (rgRNA) identification pipeline.** This table summarizes the number of candidate guide RNAs retained at each stage of the rgRNA identification pipeline (see **Figure S3** and **Methods**) for human, mouse, and zebrafish genomes. Rows indicate sequential filtering criteria: initial identification of guides targeting repetitive elements with moderate or better predicted efficiency (row 2), removal of guides with exonic targets and narrowing to 20–200 genome-wide targets (rows 3–5), merging of guides with >60% overlapping target sites (row 6), primer design and screening (rows 7–8), and final selection of rgRNAs meeting ≥50% recovery and ≥10% on-target fraction thresholds (row 9). Row 10 reports the overall yield—the fraction of initial repeat-targeting guides (row 2) that passed all filtering criteria to become validated rgRNAs. All counts reflect targets per haploid genome.

**Table S4. Catalog of repeat-targeting guide RNAs (rgRNAs) across human, mouse, and zebrafish genomes.** This table contains all rgRNAs identified by our computational pipeline (see **Figure S3** and **Methods**) that have 20–200 genome-wide targets, no exonic targets, and at least one primer pair achieving ≥50% recovery and ≥10% on-target fraction. Separate sheets are provided for human (hg38; n = 15,201), mouse (mm10; n = 25,606), and zebrafish (danRer11; n = 18,822). All values are per haploid genome. Spacer sequence: 20-nucleotide guide RNA spacer sequence. Repeat element: RepeatMasker annotation of the targeted repetitive element; if target sites are present in multiple repeat elements, all elements are listed. Repeat family: repeat family classification. Repeat class: repeat class classification. Total targets: number of genome-wide target sites (allowing up to 2 mismatches outside the seed region). Exact-match targets: number of target sites with perfect spacer complementarity. CRISPRater score: predicted guide efficiency score (range 0–1; all rgRNAs have scores ≥0.55). Functionality hazard score: cumulative score quantifying potential overlap with regulatory elements across all target sites (see **Methods**). Functionality hazard percent: hazard score divided by total target count, representing the average regulatory risk per target. Optimal primer pair: forward and reverse primer sequences separated by an underscore. Targets covered: number of target sites amplified by the primer pair. Targets missed: number of target sites not amplified by the primer pair. Off-target amplicons: number of amplicons produced that do not contain the rgRNA target sequence. On-target fraction: fraction of total amplicons containing the rgRNA target (Targets covered / [Targets covered + Off-target amplicons]); a value of 1 indicates no off-target amplicons are expected. Distinguishable amplicons: number of different target-containing amplicon sequences (at least one SNV or indel distinguishing them other amplicon sequences). Amplicon length: length of the expected amplicon in base pairs. SDI: sequence distinguishability index, calculated as (m−1)/(n−1) where m is the number of distinguishable amplicons and n is the total number of covered targets; SDI = 1 indicates all amplified targets are uniquely identifiable, SDI = 0 indicates all are identical.

**Table S5. Oligonucleotides used for cloning.** This table lists all oligonucleotides used for cloning in this study, including primers for PCR amplification, spacer oligos, and 3’ extension oligos. All oligonucleotides were ordered from Integrated DNA Technologies (IDT). Name: descriptive name indicating the oligo’s purpose. Sequence: oligonucleotide sequence in 5’ to 3’ orientation. Some oligos contain overhangs or homology arms for Gibson assembly, Golden Gate assembly, or other cloning methods.

